# A conserved RWP-RK transcription factor VSR1 controls gametic differentiation in volvocine algae

**DOI:** 10.1101/2023.03.26.534280

**Authors:** Sa Geng, Takashi Hamaji, Patrick J. Ferris, Minglu Gao, Yoshiki Nishimura, James Umen

## Abstract

Volvocine green algae are a model for understanding the evolution of mating types and sexes. They are facultatively sexual, with gametic differentiation occurring in response to nitrogen starvation (-N) in most genera, and to sex inducer hormone (SI) in *Volvox*. The conserved RWP RK family transcription factor (TF) MID is encoded by the *minus* mating type (*MT*) locus or male sex-determining region (SDR) of heterothallic volvocine species and dominantly determines *minus* or male gametic differentiation. However, the factor(s) responsible for establishing the default *plus* or female differentiation programs have remained elusive. We performed a phylo transcriptomic screen for autosomal RWP-RK TFs induced during gametogenesis in unicellular isogamous *Chlamydomonas reinhardtii* (Chlamydomonas) and in multicellular oogamous *Volvox carteri* (Volvox) and identified a single conserved ortho-group we named Volvocine Sex Regulator 1 (VSR1). Chlamydomonas *vsr1* mutants of either mating type failed to mate and could not induce expression of key mating-type-specific genes. Similarly, Volvox *vsr1* mutants in either sex could initiate sexual embryogenesis, but the presumptive eggs or androgonidia (sperm packet precursors) were infertile and unable to express key sex-specific genes. Yeast two-hybrid assays identified a conserved domain in VSR1 capable of self-interaction or interaction with the conserved N terminal domain of MID. *In vivo* co-immunoprecipitation experiments demonstrated association of VSR1 and MID in both Chlamydomonas and Volvox. These data support a new model for volvocine sexual differentiation where VSR1 homodimers activate expression of *plus*/female gamete-specific-genes, but when MID is present MID-VSR1 heterodimers are preferentially formed and activate *minus*/male gamete-specific-genes.

**Significance Statement:** Sex and recombination are conserved features of eukaryotic life cycles, but sex determination mechanisms are diverse, and are poorly understood in most major taxa. Our study identified a long-sought regulator of sexual differentiation in volvocine green algae—the RWP-RK family transcription factor (TF) VSR1— leading to the first complete paradigm for mating type or sex determination in this lineage. Our results support a model where gametically expressed VSR1 homodimerizes and activates *plus*/female specific genes. When the dominant sex-linked *minus*/male RWP-RK family TF MID is present MID-VSR1 heterodimers are preferentially formed and activate *minus*/male genes. The widespread association of RWP-RK TFs with gamete differentiation in the green lineage suggests that a similar paradigm may operate throughout the plant kingdom.

## Introduction

Sexual reproduction is a ubiquitous part of eukaryotic life cycles. A key feature of eukaryotic sex is the specification of gamete types which can help ensure that two non-related individual gametes fuse during fertilization and enable the transition to zygotic differentiation (1). In isogamous species, which include many single celled protistan lineages, gametes are defined by mating types whose numbers can range from two in most taxa to many thousands in some fungi (2). Many multicellular taxa are anisogamous or oogamous with two sexes that are defined as male and female based on size and motility of their gametes, with males producing many small motile gametes and females producing fewer large and sometimes immotile gametes. An astonishing diversity of mechanisms have evolved for specifying mating types and sexually dimorphic gametes, including genetic mechanisms governed by mating-type loci or sex chromosomes, and epigenetic mechanisms where a single individual can produce gametes of either mating type or of either sex. Genetic programs that specify gamete differentiation have been well studied in animals and fungi, but much less is known about the logic of gamete specification in other taxonomic groups. Green algae and land plants share common ancestry, but knowledge of mechanisms for gamete differentiation and sex determination in this group is incomplete, especially in green algae where molecular information is only known for a few species (3).

Green algae in the volvocine lineage have mating type and sex determination systems that are some of the best understood among algae and other protists. Volvocine algae include *Chlamydomonas reinhardtii* (Chlamydomonas), a bi-ciliate unicellular species with two mating types, and *Volvox carteri* (Volvox), a multicellular species with dimorphic sexes and UV sex chromosomes (4–6). Like most green algae, volvocines are haplontic with mating type or sex of heterothallic species determined in the haploid phase by mating type or sex determining loci . Sex in volvocine algae is facultative with sexual reproduction triggered by absence of nitrogen (-N) in Chlamydomonas and most other volvocine genera, or triggered by a sex inducer hormone (SI) in Volvox. Upon fertilization a diploid zygote is formed which immediately differentiates into a dormant spore. When germinated the spore undergoes meiosis to produce new haploid progeny (Figures 1A and 1B).

**Figure 1.**
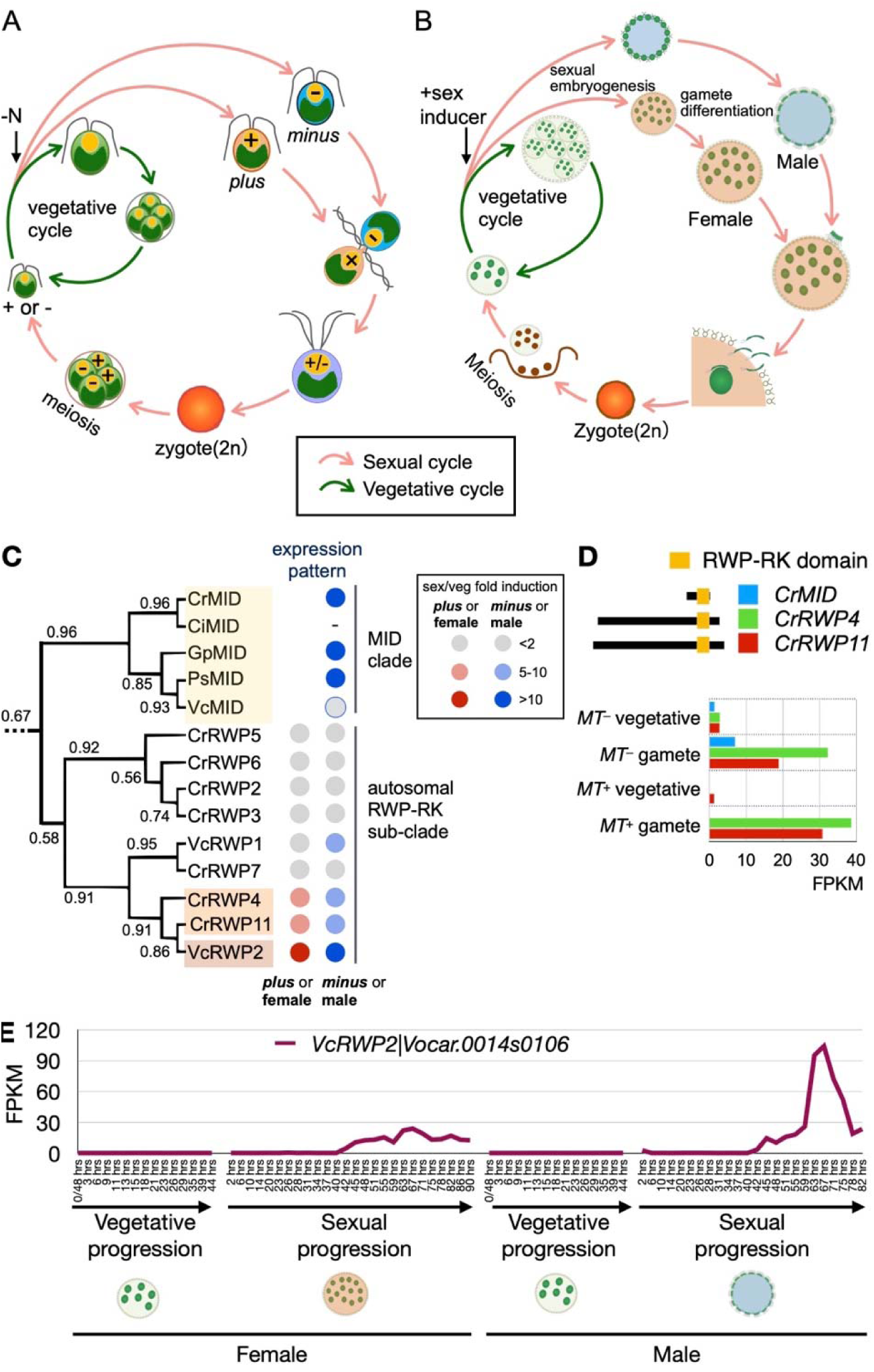
Life cycles of *Chlamydomonas reinhardtii* (Chlamydomonas) and *Volvox carteri* (Volvox) (A, B) and phylogenomic screening of volvocine RWP-RK family genes expressed during sexual reproduction (C-E). A. Chlamydomonas life cycle. During vegetative reproduction (green arrows) cells grow and divide by multiple fission. The sexual cycle (pink arrows) is triggered when nitrogen is absent (-N). Gametic differentiation occurs under the control of the *MT^+^* and *MT^−^* mating-type loci that specify *plus* or *minus* gametes. Ciliary agglutination is followed by fertilization and differentiation into a diploid zygospore. Upon germination meiosis occurs yielding four recombinant haploid progeny. B. Volvox life cycle. During vegetative reproduction of both sexes (green arrows) large stem cells in each spheroid called gonidia undergo embryogenesis and produce a new generation of spheroids. Exposure to sex inducer triggers the sexual cycle (pink arrows) leading to dimorphic embryogenesis in males and females under the control of the male (*MTM*) or female (*MTF*) sex determining regions (SDRs). Post embryonic maturation of sexual germ cell precursors leads to their differentiation into sperm packets or eggs. Sperm packets produced by males swim to females, enter the female extracellular matrix (ECM) and fertilize the large immotile egg cells that then develop into dormant diploid zygospores. Upon germination a single meiotic product survives and develops into a new haploid recombinant progeny. (C) Phylogenetic subtree of selected RWP-RK family genes in volvocine algae. MID genes (restricted to *minus*/male) are indicated at the top and autosomal RWP-RK genes at the bottom. For the mRNA expression patterns, gray circles indicate weak or no induction (<2-fold) in sexual versus vegetative phases; pink indicates 5-10 fold induction and red >10-fold induction in *plus*/female sexual phase; pale blue indicates 5-10 fold induction and dark blue indicate >10 fold induction in *minus*/male sexual phase. Colored rectangles highlight gene families of special interest. (D) Expression patterns of Chlamydomonas sexually expressed RWP-RK genes. FPKM values are extracted from (16). Protein domain schematics with RWP-RK domain in pale orange are shown next to the color key for mRNA expression of each gene. (E) Volvox *RWP2* expression pattern across the vegetative and sexual development cycles. Arrows indicate time progression in the four sample types starting from pre-cleavage spheroids for vegetative samples and from addition of sex inducer for sexual samples: female sexual, female vegetative, male sexual, and male vegetative.

First discovered in the mating type *minus* (*MT^−^*) locus of Chlamydomonas, the *MID* (*<u>MI</u>NUS <u>D</u>OMINANCE*) gene encodes a small RWP-RK family transcription factor (TF) and is responsible for *minus* gamete differentiation: introduction of a *MID* gene into a *MT^+^* strain, or its presence in a heterozygous diploid dominantly cause *minus* differentiation (7, 8). Conversely, in *mid* mutants a default program of *plus* gamete differentiation occurs. *MID* is conserved across volvocine species and is found in the *MT^−^* locus or male SDR of all heterothallic species (6). In Volvox *MID* was found to be a dominant determinant of spermatogenesis, with oogenesis occurring in its absence, though other aspects of dimorphic sexual development were not controlled by MID (9). Although direct targets of MID activation or repression are not established, they likely include gamete-type specific genes including *minus* genes *GSM1* and *SAD1* which encode a homeodomain TF required for zygote specification and the *minus* sexual agglutinin respectively, and *plus* genes *GSP1* and *SAG1* which encode a homeodomain TF partner for GSM1 and the *plus* sexual agglutinin binding partner for SAD1 (10–12).

Although presence/absence of *MID* is the key determinant of *minus* or male gamete differentiation in volvocine algae, it has remained unclear how the default or ground state *plus* or female differentiation program is specified in its absence. Although extensive mutagenesis screens have been done in Chlamydomonas, no mutants that specifically interfere with *plus* differentiation have been identified (13, 14). Moreover, mutants that impact gametogenesis of both mating types are difficult to investigate because they are sterile (15), so to date no factors that impact *plus* or female gamete differentiation have been positively identified in volvocine algae and information about this important aspect of their sexual life cycle has remained missing.

Here we used a phylo-transcriptomic search to identify a key missing gamete differentiation factor in volvocine algae, a conserved RWP-RK TF we designated as VSR1 (<u>v</u>olvocine <u>s</u>ex regulator). VSR1 was induced in both mating types of Chlamydomonas in response to -N, while in Volvox its expression was induced by SI in both sexes. *vsr1* mutants were validated or created for Chlamydomonas and Volvox where they caused sterile phenotypes in both mating types or sexes and failed to express key gamete differentiation genes. Yeast two-hybrid experiments identified a region of VSR1 that could mediate both self-interaction and interaction with the N-terminal domain of cognate MID proteins. Interactions between MID and VSR1 in Chlamydomonas and in Volvox were further validated *in vivo* using co-immunoprecipitation. Our data support a model where VSR1, when expressed alone, self-dimerizes and activates *plus* or female differentiation genes, and that when co-expressed with MID preferentially forms VSR1-MID heterodimers which can activate *minus* or male gamete differentiation genes. These data provide a new paradigm for mating type or sex determination by RWP-RK TFs and have implications for the evolution of gamete differentiation that is likely controlled by RWP-RK TFs in other green lineage species.

## Results

### Phylogenomics search for conserved volvocine gametogenesis regulators

We reasoned that the default gametic differentiation state of volvocine algae that is adopted when gametogenesis cues are present (-N for Chlamydomonas and +SI for Volvox), but when MID is absent, could be controlled by an autosomal TF that is expressed in both mating types or sexes. The activity of this hypothetical TF would need to be modified or inhibited by MID in *minus* or male gametes since MID is dominant (7, 9). We further guessed that this TF might be in the RWP-RK family. In Chlamydomonas extensive transcriptome responses to -N treatment and gametogenesis have been documented, including the induction of several RWP-RK TFs, but most of the transcriptome changes in -N are related to starvation and growth arrest and not to gametogenesis or sex (16–19). Having an orthogonal set of data for Volvox where gametogenesis occurs in nutrient replete conditions provided a potential phylogenetic filter to identify conserved gametogenesis genes. Using time series data from *V. carteri* (Figure S1; SRA Accession No. PRJNA977067; Hamaji *et al*., in preparation) and previously published data on *plus* and *minus* gamete gene expression from *C*. *reinhardtii* (16, 17) we searched for related RWP-RK TFs that were upregulated in both species by their respective cues for gametogenesis. A single candidate sub-family of autosomal RWP-RK TFs fit this pattern. This subfamily had a single member in Volvox, *VcRWP2* (Vocar.0014s0106), and two closely related paralogs in Chlamydomonas, *CrRWP4* (Cre03.g149350) and *CrRWP11* (Cre03.g149400) that are adjacent to each other on Chromosome 3 (Figures 1C and S1A). All three of these genes showed increased expression in response to the respective gametogenesis cues in each species (Figures 1C-1E). We subsequently identified single *RWP2* candidate orthologs in other multicellular volvocine genera (a lineage comprising Tetrabaenaceae, Goniaceae, and Volvocaceae a.k.a. the TGV clade), and the same tandem duplication in two other *Chlamydomonas* species that are closely related to *C. reinhardtii*—*C. incerta* and *C. schloesseri*—but only a single ortholog in *Edaphochlamys debaryana* (Figure S2B-F) (20). These findings indicate a likely origin of the duplication within Metaclade-C of the volvocine algae but not in the TGV clade that contains most of the multicellular volvocine genera (Figure S2E). The paralogous duplicates in the Metaclade-C *Chlamydomonas* species are highly similar in their predicted C-terminal RWP-RK domains and share different degrees of similarity in their long N terminal domains (Figure S2F).

### *Chlamydomonas rwp11/vsr1* mutants are mating defective

Chlamydomonas *rwp4* and *rwp11* mutants were obtained from the CLiP collection of insertional strains which are in a *MT*^−^ strain background (21) (Figure 2A-B; Figure S3). The *rwp4 MT*^−^ strain was confirmed by genotyping and found to mate and produce viable offspring. Both *MT^+^* r*wp4* and *MT^−^ rwp4* progeny mated successfully with strains of opposite mating type, and this mutant was not pursued further (Table S1). The *MT*^−^ *rwp11* mutant strain could not mate and was blocked prior to agglutination with wild-type *MT^+^* gametes. It was also unable to agglutinate or mate with wild-type *MT^−^* gametes meaning that it had not adopted a pseudo-*plus* mating program as is observed in *mid* mutants (7).

**Figure 2.**
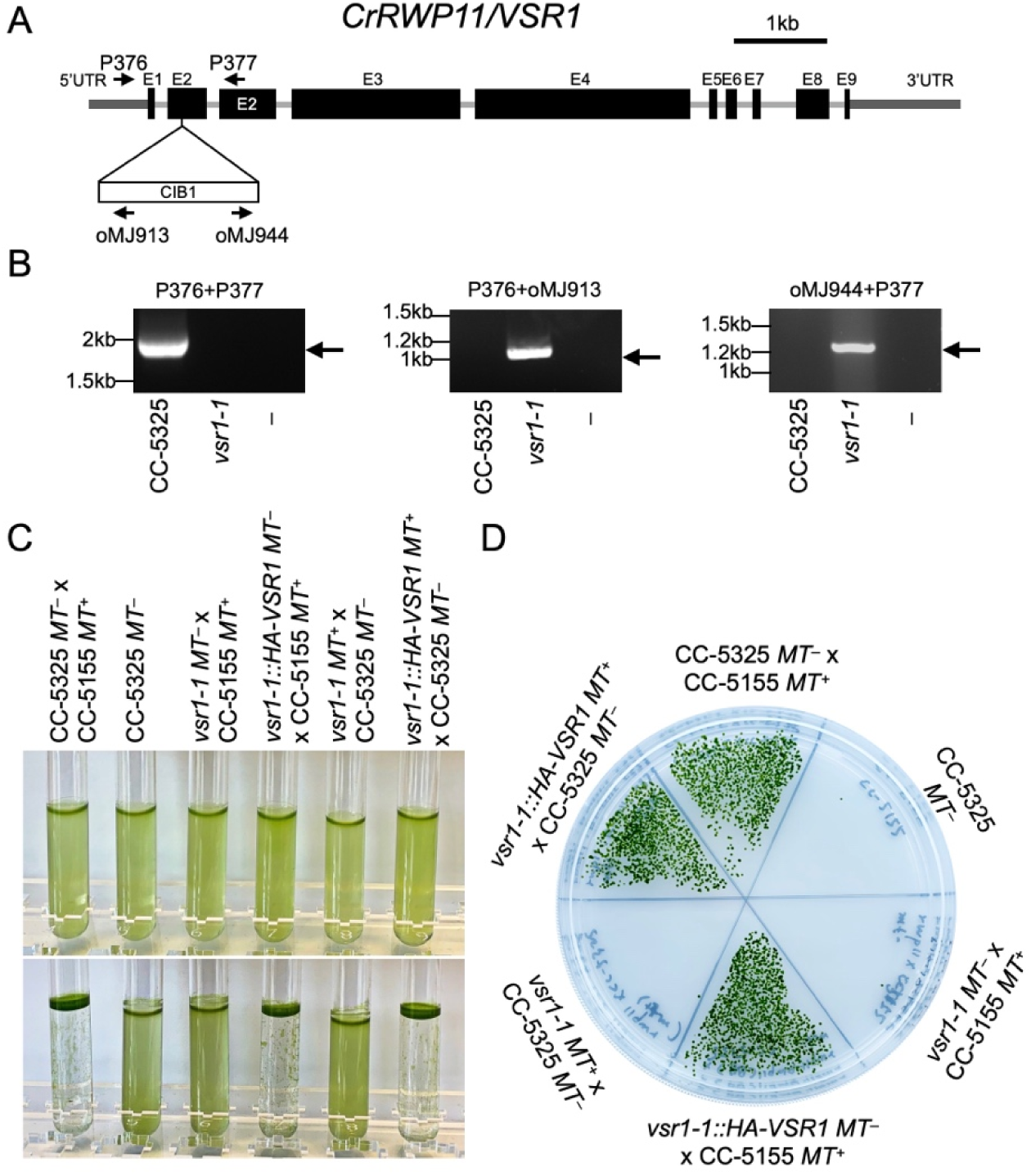
Sterility phenotype of Chlamydomonas *vsr1* mutants. A. Diagram of *RWP11* (*VSR1*) locus and location of CIB1 cassette in exon 2 (E2) of mutant strain LMJ.RY0402.189640 whose mutant allele is designated as *vsr1-1*. Locations and names of PCR primers used for genotyping are shown above. Exons are indicated by thick black boxes and labeled E1-E9. 5’ and 3’ untranslated regions (UTRs) are indicated by thinner dark gray lines. B. PCR genotyping of *VSR1* and *vsr1-1* alleles. PCR products were run on agarose gels and imaged after staining with RedSafe dye. Strain CC-5325 *MT^−^* is the parent strain of *vsr1-1* and serves as a wild-type control. A no-template control for each primer pair is designated as –. C. Mating reactions between pairs of indicated strains (or CC-5325 as a negative control). Top row of tubes shows gamete suspensions just after mixing. Bottom row shows tubes after an overnight incubation. Successful mating results in buoyant mats of zygote pellicle on the surface and sides of the tube and a clear appearance. D. Matings from panel C were spread in sectors on an agar plate prior to pellicle formation, allowed to mature, and chloroform treated to kill unmated gametes prior to germination. Colonies are formed by germinated zygospores

In order to pursue the *rwp11* mutant further a rescuing construct containing a N-terminal hemagglutinin tagged version of the *RWP11* gene driven by a strong promoter/terminator from the *RPL23* gene (22) (pHA-VSR1; Figure S4A) was introduced into the *rwp11* mutant strain by transformation. Multiple independent transformants were recovered that expressed the tagged transgene constitutively (Figure S4B) and these strains had restored ability to mate with an isogenic *MT^+^* parental strain CC-5125 (Figure 2C). Segregation behavior of the *rwp11* mutation and rescuing construct were as expected, enabling the characterization of the *rwp11* phenotype in both mating types (Table S2). *rwp11* mutant progeny of both mating types were sterile, but those which also inherited the rescuing transgene could mate as could all progeny that inherited a wild-type *RWP11* allele (Table S3, Figure S5). During the process of outcrossing and genotyping *rwp11*, a second insertion cassette and paromomycin resistance marker was discovered in the original *rwp11* CLiP strain in an unlinked gene, *SYP1* (Cre13.g588550), encoding a predicted vesicle trafficking protein. The presence/absence of the *syp1* mutation did not affect the mating phenotypes of any of the progeny strains that were tested (Table S3) meaning that this second mutation is unrelated to the mating defect of *rwp11*. Progeny in which the *syp1* mutation was segregated away from the *rwp11* mutation were chosen for further study. We renamed the *RWP11* gene *VSR1* (volvocine sex regulator 1).

### Volvox *vsr1* mutants are defective for gamete differentiation

To test whether the *Volvox carteri* VSR1 homolog (originally *RWP2*, Vocar.0014s0106) was involved in sexual differentiation or mating we used a gene editing method based on CRISPR-Cas9 to make frameshift alleles (23). We made multiple independent edits in either the fourth or seventh exons of *VSR1* in a male strain (Figure 3A, Figure S6). All *vsr1* frame-shift male strains tested had normal vegetative phenotypes that were indistinguishable from wild type (Figure 3B,E). In response to SI, *vsr1* mutants underwent a typical male early embryogenesis program to produce spheroids with 128 small somatic cells and 128 large cells that resembled uncleaved androgonidia (Figure 3C,F). However, unlike wild-type males, the androgonidial-like cells in *vsr1* mutants did not cleave into sperm packets, but instead underwent a vegetative embryogenesis program and reentered the vegetative life cycle (Figure 3D). This novel phenotype is different from that seen in sexually induced males with *mid* knockdowns where the post-embryogenesis large cells trans-differentiated into eggs (9).

**Figure 3.**
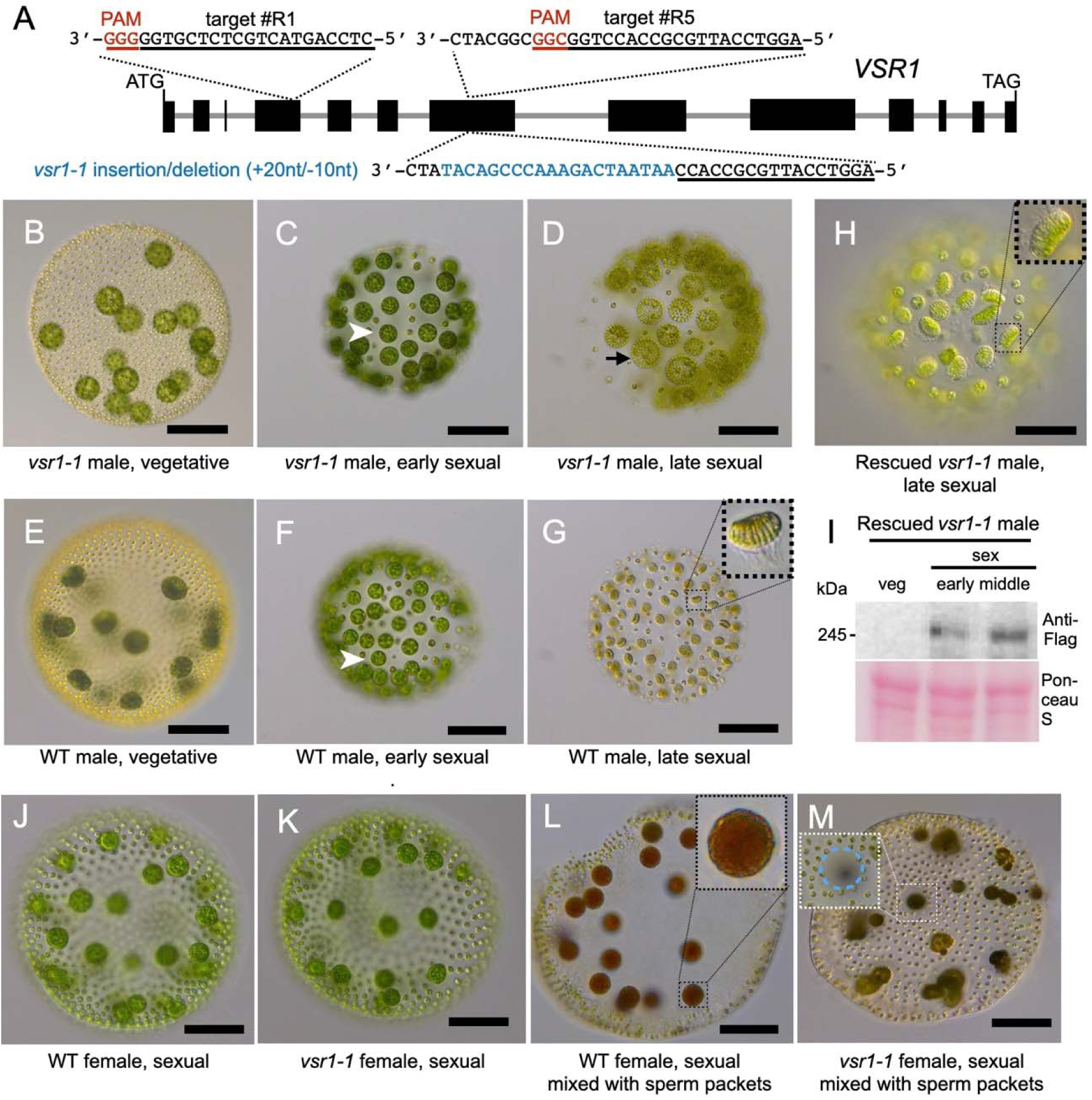
Isolation and characterization of Volvox *VSR1* edited mutants and rescued strains. (A) Diagram of Volvox *VSR1* locus showing two different editing sites in detail including the PAM site that specifies cleavage by targeted Cas9 guide RNA RNP complexes. The *vsr1-1* insertion/deletion allele is shown below in blue lettering. Dark boxes represent exons. Start and termination codons are also shown. (B-D). Micrographs of male *vsr1-1* mutant taken from vegetative cultures (B), from post-embryonic sexual males with uncleaved presumptive androgonidia with an example shown by a white arrowhead (C), and from a stage when the presumptive androgonidia had undergone vegetative embryogenesis to produce vegetative juveniles (black arrowhead) instead of sperm packets (D). (E-G) Micrographs of a wild-type male strain taken at equivalent stages as the *vsr1* mutant in panels B-D, respectively. A mature male with sperm packets (G) is shown with a magnified sperm packet in the inset. (H). Rescued *vsr1-1::SF-VSR1* male strain at sexual maturity with restoration of sperm packets. A magnified sperm packet is shown in the inset. All scale bars are 50 µm. (I). Immunoblot detection of FLAG epitope tagged VSR1 in a rescued *vsr1-1* strain. Samples were taken from vegetative (veg) and from early (48 hrs), and middle (59 hrs) stages of sexual (sex) development as depicted in Figure S1. (J) Wild-type sexual female with eggs. (K) Sexually induced *vsr1-1* female mutant spheroids with egg-like cells. (L) Fertilized wild-type female with maturing zygotes. Inset in (L) shows a magnified view of a zygote. (M) Post-mating *vsr1-1* sexual females with fertilization pore at sperm entry point (inset) but no zygotes. The egg-like cells behave as unfertilized *vsr1-1* mutants and undergo vegetative embryogenesis.

Using a representative allele, *vsr1-1* (*vsr1*^#5_4^, Figure S6), a rescue experiment was performed by transforming the mutant with construct pSF-VSR1 containing the *VSR1* gene and its native promoter with a terminator sequence from the *NitA* gene. Tandem StrepII and FLAG epitope tags were inserted in the N-terminus of *VSR1* in pSF-VSR1 to facilitate detection of the VSR1 protein (Figure S7). Several of the transformants produced with this construct showed a rescued phenotype where sperm packet development occurred normally after sexual induction (Figure 3H). The rescued strains also produced detectable tagged VSR1 protein of the expected size on immunoblots in sexually induced spheroids (Figure 3I). These results indicated that defects in *VSR1* were causative for the male gametogenesis defect that was observed in the *vsr1* mutant strain. Immunofluorescent staining of sperm cells from the rescued *vsr1-1 SF-VSR1* male strain with anti-StrepII antibody showed that tagged transgenic VSR1 localized to nuclei as expected for a predicted TF (Figure S8).

The rescued *vsr1-1* male strain was then crossed to a wild-type female strain (Figure 3, Figure S9). Normal segregation patterns were observed in progeny, some of which were *vsr1-1* mutant females (Table S4, Figure S10). Like the case in males, all female *vsr1-1* progeny without a co segregating rescuing construct had normal vegetative phase phenotypes, but were sterile. Upon sexual induction they underwent a normal sexual early embryogenesis program to make spheroids that had around 2000 somatic cells and 32-48 large egg-like cells. However, the egg like cells in the female *vsr1-1* mutants could not be fertilized when incubated with mature sperm from a wild-type male, despite the sperm being able to enter the females through a fertilization pore and adhere to the egg-like cells, sometimes remaining associated for days (Figure 3K,M; Figure S9D-F). Eventually the egg-like cells underwent vegetative embryogenesis and reentered the vegetative life cycle, though with different timing than unfertilized wild-type eggs which experience a delay before differentiation (Figure S9D,E). Though we think it unlikely, we cannot rule out that some of the *vsr1-1* eggs underwent fertilization but were blocked post-zygotically due to lack of *GSP1* expression (see below). We also noted that the vegetative embryos produced by the egg-like cells of *vsr1-1* mutants frequently failed to invert, though this phenotype disappeared in all subsequent vegetative generations (Figure S9D,E). This one-time inversion phenotype may be related to the size or expansion capacity of the vesicle surrounding the egg-like cells, a trait which is known to influence inversion (24). As expected, male *vsr1-1* mutants behaved as they did in the parental mutant strain, and all female spheroids that received a rescuing construct or that inherited the wild-type *VSR1* allele underwent normal female sexual development (Figure 3L; Figures S9G and S10).

### *VSR1* is required to express mating type/sex specific genes

Chlamydomonas gametes express several mating-related genes that are specific to either *plus* (e.g. *SAG1*, *GSP1*) or *minus* (e.g. *SAD1*, *GSM1*) gametes (10–12). Quantitative reverse transcription and PCR (qRT-PCR) was used to determine whether these marker genes were expressed normally in *vsr1* gametes. These experiments revealed that *vsr1-1* mutants, but not parental or rescued strains, were defective for expressing mating-type specific genes (Figure 4A, B). We also noted that the pHA VSR1 rescued strains, which express *VSR1* constitutively, did not show ectopic expression of mating-related genes in either mating type (Figure 4A, 4B). Together our data show that VSR1 in Chlamydomonas is necessary for gametic differentiation and mating of both *plus* and *minus* mating types, and the sterile phenotype of *vsr1-1* mutants is wholly or partly due to defects in activating mating-type specific gene expression. *VSR1* is the first gene identified in Chlamydomonas that is necessary for differentiation of both gamete types.

**Figure 4.**
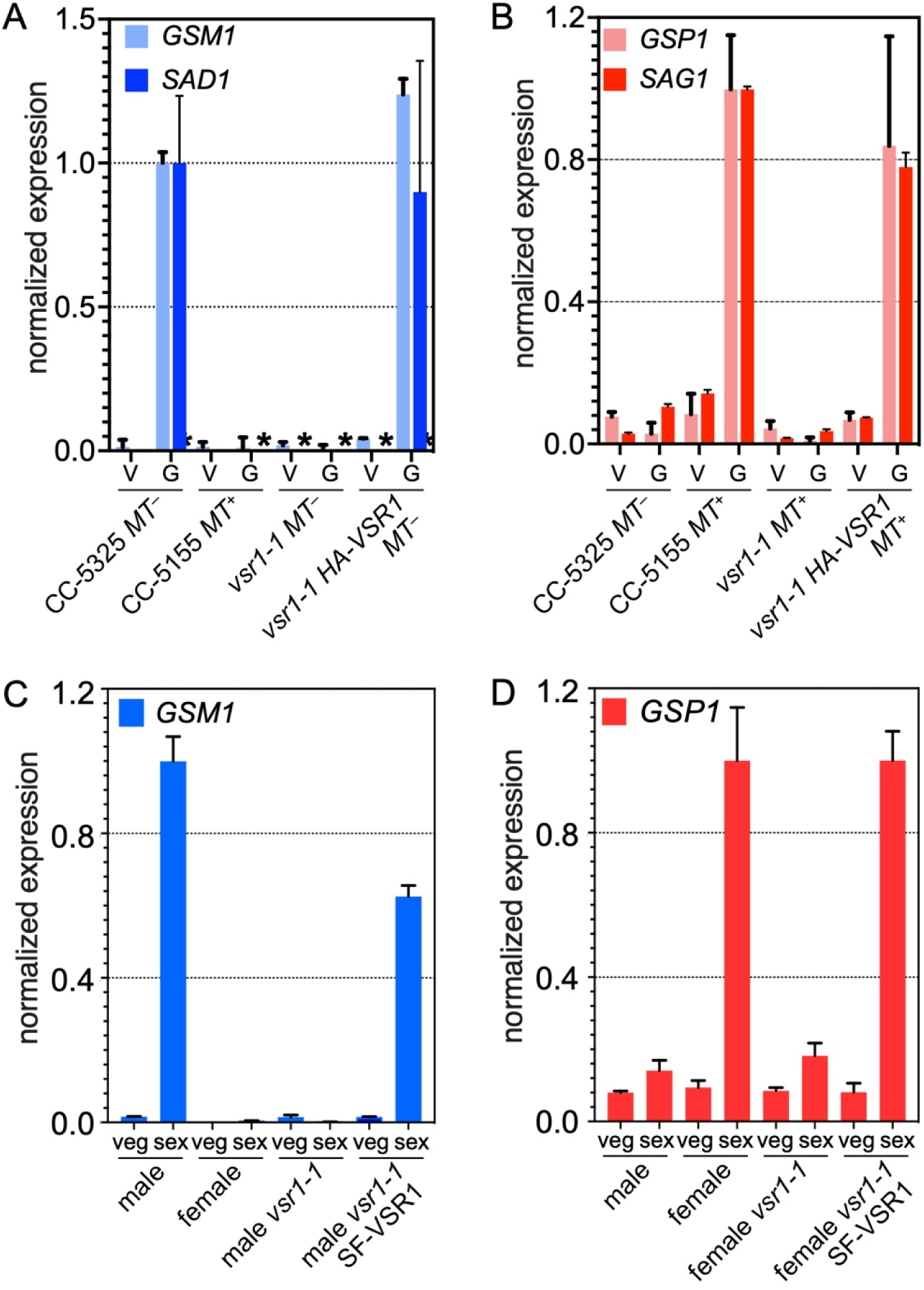
Impact of *vsr1* mutants on cell-type-specific gamete transcript accumulation. (A, B) Expression patterns of *minus* and *plus* gamete specific genes in Chlamydomonas wild type (CC-5325 or CC-5155), *vsr1-1* mutants, and rescued *vsr1-1* mutant strains expressing HA-VSR1. Transcript levels of vegetative cells (V) or gametic cells (G) were measured using quantitative RT-PCR (qRT-PCR) with values normalized to internal control ribosomal protein gene *RPL36a* and then re-scaled as relative expression to wild type *minus* or *plus* gamete values, respectively. A, *minus* genes *GSM1* and *SAD1*; B, *plus* genes *GSP1* and *SAG1*. * No expression detected. (C, D) Expression patterns of Volvox *GSP1* (female-specific) and *GSM1* (male-specific) orthologs from qRT-PCR normalized to 18S rRNA, and then re-scaled to wild type male or female values, respectively. RNA samples were derived from vegetative cultures (veg) or mature sexual cultures (sex) from indicated genotypes.

Volvox has single orthologs of *GSP1* and *GSM1* homeodomain protein coding genes that were designated *VcGSP1* and *VcGSM1* (Vocar.0026s0144 and Vocar.0053s0055, respectively; Figure 4C, D). Volvox has no detectable SAG1 homolog, but the female SDR has a *SAD1*-related gene (5) whose function is unknown but is unlikely to be essential for mating since male *MID* knockdown lines make functional eggs in its absence (9). Using qRT-PCR we determined that *VcGSP1* was expressed only in female sexual spheroids and *VcGSM1* only in male sexual spheroids, indicating that members of this heterodimeric pair retained the same expression polarity as in Chlamydomonas with respect to *MT* or SDR haplotype (Figure 4C, D). Expression of each of these Volvox genes was lost in the *vsr1-1* mutant strains but restored in rescued mutant strains. Expression timing of *VSR1* before *GSP1* or *GSM1* in Volvox was also consistent with VSR1 being an activator of these downstream TFs (Figure S1). In summary VSR1 is a conserved volvocine putative RWP-RK family TF that is essential for gametic differentiation and controls both mating-type-specific or sex-specific gene expression.

### VSR1 proteins interact with themselves and with MID proteins via a novel protein interaction domain

A simple model that could explain bi-stable and mutually exclusive gamete differentiation programs involves VSR1 forming homodimers (or other homomeric forms) in the absence of MID to activate expression of *plus* or female gamete differentiation genes. When MID is co-expressed with VSR1, MID-VSR1 heterodimers (or higher order heteromeric forms) with *minus*/male promoter specificity would outcompete VSR1 homodimers to activate *minus* or male gamete differentiation genes. This model predicts that MID and VSR1 associate with each other, and that VSR1 can self-associate.

We tested VSR1 and MID associations with themselves and each other using a yeast two-hybrid (Y2H) assay (Figure 5A-D, Figure S11). Full length MID produced a negative result when co expressed as bait and prey (Figure 5A,B), so lacks the ability to homodimerize in this assay. In the case of CrVSR1 or VcVSR1, full length bait constructs showed auto-activation, so could not be used to test self-association. However, Using CrMID as the bait construct, a positive interaction was detected with CrVSR1 as prey (Figure 5A). Truncations of CrVSR1 prey constructs showed that a 261 aa region in its N-terminus (CrVSR1^431-691^) was sufficient to interact with CrMID. We termed this region of VSR1 as its dimerization domain (DD). The DD sequences of VSR1 are conserved across volvocine algae (Figure S2C) but are not a previously defined domain detectable with NCBI BLAST conserved domain search. Using Volvox constructs we found VcMID-bait and VcVSR1-prey or VcVSR1^485-694^-prey (DD only) exhibited interaction, confirming that this domain and its interaction with a cognate MID protein is functionally conserved (Figure 5B). MID proteins are small, with a C terminal RWP-RK domain and a conserved N terminal region. Using Chlamydomonas constructs with just the MID N domain or MID C domain as bait, VSR1 was found to interact only with the N domain and this interaction may explain why the N domain of MID is so well conserved (Figure 5D).

**Figure 5.**
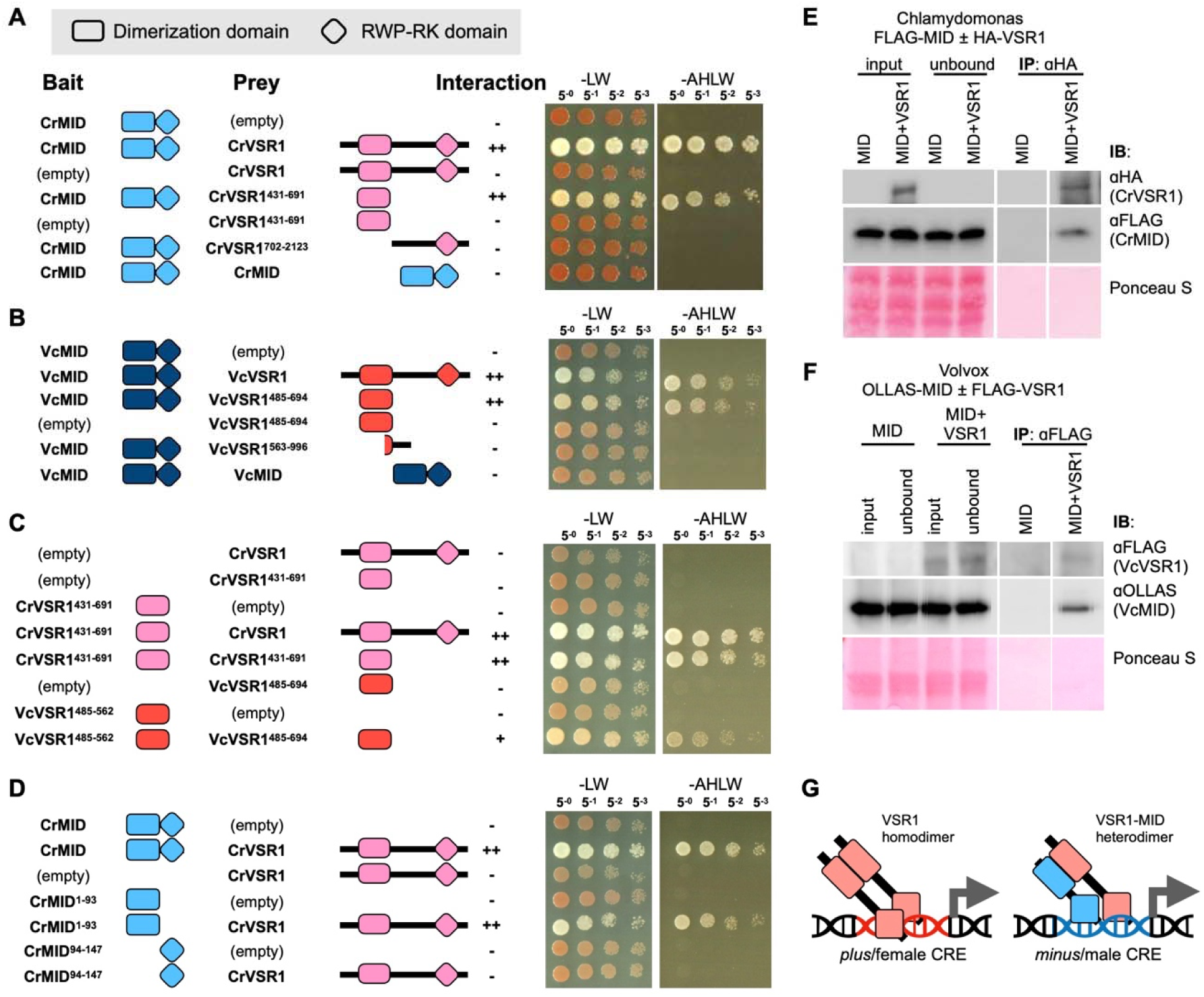
VSR1 interacts with itself or with MID via an N-terminal domain. (A-D) Yeast two hybrid experiments with schematics of bait and prey constructs shown on the left, and dilutions of yeast expressing indicated construct pairs spotted on the right. Results are summarized as - (no interaction), + (weak interaction) or ++ (strong interaction). The RWP-RK domains are depicted as diamonds and the protein dimerization domains (DDs) are depicted as rectangles. VSR1 is shown in pink (Chlamydomonas) or red (Volvox), and MID is shown in light blue (Chlamydomonas) or dark blue (Volvox). Empty indicates empty vector controls. Growth of two independent co-transformants spotted in 1:5 serial dilutions on permissive media (-LW) or interaction-dependent media (-AHLW) are shown. White colonies on -LW and growth on -AHLW indicate interaction between introduced bait and prey proteins. (E,F) Co-immunoprecipitation (IP) of VSR1 and MID in Chlamydomonas (E) and Volvox (F). (E) Co-IPs for Chlamydomonas were done in two strains, both expressing a FLAG-epitope tagged transgenic MID protein rescuing a *mid-2* deletion allele (7). One strain also expressed a transgene encoding a HA-epitope tagged VSR1 protein as described in Figure S4A. Immunoblots (IBs) with indicated antibodies are shown in each row with the Ponceau S stained membrane below. Input and unbound lanes were loaded with 1/60 of the total sample. Pellet lanes were loaded with 1/10 of the total sample. (F) Co-IP with _α_FLAG from sexually induced Volvox pseudo-males expressing OLLAS-tagged MID with and without FLAG tagged VSR1. Loading amounts were the same as in (E) for input, unbound and pellet fractions. (G) Model for bi-potential gamete differentiation based on VSR1-VSR1 homodimers binding to and activating promoters of *plus*/female genes and VSR1-MID heterodimers binding to and activating *minus*/male genes. See main text for additional details.

Because VSR1 autoactivates when used as bait in Y2H assays, the full-length protein could not be used as a bait. We therefore tested the VSR1-DD region (CrVSR1^431-691^) as bait since this fragment did not autoactivate. When co-expressed with full length CrVSR1 as prey, a positive interaction was detected indicating that CrVSR1 can self-interact through its DD (Figure 5C). When just the VcVSR1 DD was expressed in both bait and prey constructs the interaction was also detected, and a similar result was found for the DD of VcVSR1. These data support the idea that VSR1 can homo-dimerize through its DD, and this same domain can interact with the MID N terminal domain.

To determine if the MID-VSR1 interaction occurs *in vivo* we used a co-immunoprecipitation (Co IP) assay from algal lysates (Figure 5E, F; Figure S12). We constructed strains of Chlamydomonas or Volvox expressing epitope tagged *MID* and *VSR1* transgenes. For Chlamydomonas we prepared whole cell lysates from gametes of a double transgenic strain *MT^−^ mid-2 MID-FLAG, vsr1-1 HA-CrVSR1* or a strain with only the *MID-FLAG* transgene as a control. IPs were performed using anti-HA antibodies to capture VSR1 and immunoblots were probed with anti-FLAG antibodies to detect MID. MID-FLAG was detected only in IPs from the strain co expressing the HA-VSR1 transgene and MID-FLAG, but not the control strain indicating that these two proteins associate *in vivo* (Figure 5E). Similar results were obtained for Volvox male strains co-expressing a FLAG and StrepII tagged VcVSR1 protein and an OLLAS-tagged MID protein, or expressing only OLLAS-tagged MID (Figure 5F). In summary, our results indicate that VSR1 contains a conserved dimerization domain (DD) in its N terminal region and this domain facilitates self-interaction, most likely to form homodimers, as well as interaction with the N terminal domain of MID, most likely to form heterodimers.

## Discussion

### VSR1 is a key missing regulator for volvocine algal gametic differentiation and sex Determination

Several decades ago mutant screens were performed to identify mating and gametogenesis genes in Chlamydomonas (13, 14, 25). These screens identified mating defective or *imp* (impotent) mutants of various types, including *imp-11* which was later found to have a mutation in the *MID* gene causing a *plus* differentiation phenotype (7, 14). However, further work on several *imp* mutants was not feasible because they were unable to mate or mated very poorly. This conundrum precluded further analysis on genes that might be required for *plus* gamete differentiation. Here we made use of phylogenomics and phylotranscriptomics to identify a key missing factor for *plus*/female gamete differentiation, the conserved RWP-RK TF VSR1. The success of this approach rested on the different induction conditions for gametogenesis in Chlamydomonas versus Volvox which served as a powerful filter against the myriad stress and starvation genes induced in Chlamydomonas by -N. While our search for VSR1 was narrowly focused on one TF family, it could easily be generalized to find a larger set of genes that might have conserved functions in volvocine gametogenesis. While additional factors besides VSR1 may be required for volvocine gametic differentiation, VSR1 fulfills a key role of helping to establish the gametic default or ground state of *plus* or female in two distant volvocine cousins— *C. reinhardtii* and *V. carteri*. By extension, it seems likely that this role for VSR1 is conserved across the entire volvocine clade where we have identified candidate VSR1 orthologs in species with sequenced genomes (Figure 1; Figure S2).

The tandem duplication of *VSR1* (*RWP11*) and *RWP4* that seemed to have occurred only in *C. reinhardtii* and two close relatives is somewhat puzzling (Figure S2). The RWP-RK domains of these two paralogs are highly similar, though their N terminal regions are more diverged. However, their mutant phenotypes are completely different, thus ruling out the possibility of redundant function: *rwp4* mutants had no detectable phenotype in the sexual cycle, while *rwp11* (*vsr1*) mutants were completely sterile. Importantly, some DD residues which are conserved in volvocine VSR1 proteins are absent from RWP4 which may render RWP4 incapable of interacting with itself or with MID (Figure S2F). Nonetheless, the retention of this duplication over several speciation events and the induction of RWP4 expression by -N suggests some role in the sexual cycle or nitrogen starvation response. On the other hand, most volvocine species have only a single *VSR1*-related gene, so whatever role RWP4 acquired seems specific to Chlamydomonas and its closest relatives and this paralog is not a core component of volvocine mating-type or sex determination.

The identification of VSR1 as a factor required for gametogenesis in both mating types or sexes indicates a dual role in promoting expression of mating-type or sex-specific genes whose expression patterns are mutually exclusive. One way to reconcile these mutually exclusive roles would involve modification of VSR1 by association with MID, converting it from either a homodimer in *plus*/female (or heterodimer with another factor) to MID-VSR1 heterodimers in *minus*/males with presumably different DNA binding specificity (Figure 5G). Data from Y2H and Co-IP data are consistent with this simple model since the VSR1 DD we identified has the capacity for homodimerization with the N terminus of MID and with itself, indicating potential mutual exclusivity of VSR1 self interaction versus interaction with MID (Figure 5). This model further predicts that the MID-VSR1 interaction is stronger than the VSR1-VSR1 interaction so that VSR1 homodimers are not present or functional in *minus*/male. While these predictions— including competitive binding of MID and analysis of VSR1 homodimers *in vivo*—remain to be directly tested, our finding that VSR1 is required for expression of key mating type or sex specific genes for both mating types or sexes of Chlamydomonas and of Volvox support its role as a dual function TF for gametogenesis. Although the VSR1 and MID DDs are not detectably conserved in other species, studies with plant RWP-RK TFs identified PB1 domains in NLP type RWP-RKs that mediate interactions between NLPs and between NLPs and TCP type TFs (26). Thus it seems that dimerization and combinatorial DNA binding is a feature of at least some RWP-RK proteins.

### A new paradigm for control of sexual differentiation in volvocine algae

Molecular mechanisms for sex or mating type determination have been studied extensively in fungi and metazoans where a great diversity of epigenetic and genetic sex determination systems have evolved (1). The model we propose here for MID and VSR1 (Figure 5G) is different from previously described systems as the VSR1 protein is required in both mating types or sexes, and interacts directly with a *minus*/male TF MID to modify its function. In many fungi each haploid mating-type locus encodes a TF idiomorph that is required for mating type specific gene expression, and for activating the diploid program after fertilization (27). In the ascomycete budding yeast *S. cerevisiae*, a variant system has arisen that is superficially similar to volvocine algae. **a** mating-type differentiation is the default when *MAT*_α_ genes _α_*1* and _α_*2* are absent, just as *plus* is the default in Chlamydomonas when MID is absent. However, unlike the case in Chlamydomonas where the presence of MID dominantly converts a *plus* gamete to a *minus* gamete, the *MAT*_α_ locus is co-dominant with *MATa* and the coexpression of a1 from the *MATa* locus with _α_*1* and _α_*2* represses haploid cell type gene expression programs (28). In eutherian mammals the Y-chromosome gene *SRY* is somewhat analogous to *MID* in being a dominant specifier of male differentiation with the ground state or default being female. However, the regulatory logic is different since the male differentiation regulator on the Y chromosome SRY and its coregulator SF1 are at the top of a hierarchy in which male sexual differentiation is maintained by pushing gonadal precursor cells into a state of stable *SOX9* expression; but none of these three male-expressed TFs plays a role in maintaining female gonadal differentiation which occurs when *SOX9* expression is turned off (29). Although no evidence is available, it seems likely that MID plays a direct role in activating *minus* or male genes as a heterodimer with VSR1, which is unlike the case for SRY whose only target may be *SOX9* (29). Interestingly, the dominance relationship between MID and VSR1 would make this system compatible with a transition to a diploid XY sex determination system with MID as the Y linked male determining factor, though no natural diploid species have been found among volvocine algae.

### Deep Conservation of RWP-RK TFs in green lineage gamete specification

RWP-RK TFs are associated with proposed mating type loci, sexual reproduction and/or gametogenesis in diverse members of the Viridiplantae including several Chlorophyte algae (30–33), the Charophyte alga *Closterium peracerosum–strigosum–littorale*(Closterium) (34) and embryophytes (Table S5). Among them, conserved RWP-RK TFs in the RKD family are key factors for egg cell differentiation in embryophytes. In Arabidopsis, RKD family members are required for egg cell differentiation (35–37), and the single RKD homolog in the liverwort *Marchantia polymorpha* (MpRKD) is required for egg cell differentiation and also plays a role in spermatogenesis (38, 39). Taken together, the available information across the green lineage suggests an ancestral role for RWP-RK TFs in governing sexual differentiation in addition to their roles in nitrogen uptake and utilization (40). We speculate that a common set of RWP-RK TFs may have mediated gamete differentiation and mating-type specification at the base of the Viridiplantae or even earlier, but that rapid evolution of these genes, as has been demonstrated directly for MID (41), might obscure their common origin among modern descendant lineages where RWP-RK proteins have diversified into distinct sub-groups (40).

### Mating type specification and the emergence of sexes

In species with mating types, gametes are usually isogamous with little or no morphological differentiation between types. In many fungi where mating-type-specific gene expression is required for differentiation of mating types, each *MAT* locus has its own unique TF that directly regulates mating-type-specific genes (42). In multicellular taxa with separate sexes, two additional features often evolve. One is more complex gamete differentiation to produce male and female gamete types, and the second is integration of gametogenesis into a body plan where males and females produce distinct types of support tissues and organs. In these sexually dimorphic systems sex determination mechanisms can control both somatic and gametic differentiation through hierarchical or parallel developmental mechanisms that may govern thousands of genes either directly or indirectly (43). This emergent property of sexes in multicellular organisms is evident even in Volvox where at least two levels of control over sexual differentiation have been observed. Male and female sexual spheroids each have their own unique embryogenesis programs that produce a characteristic pattern of sexual somatic cells and germ cell precursors. Previously it was shown that the MID pathway was largely responsible for differentiation of gametic precursor cells, but that it did not control sexual embryogenesis which instead must be controlled by the Volvox U and V sex chromosomes (9, 44). Our data for VSR1 further supports this emergent role for sex chromosomes in dimorphic embryogenesis in Volvox. Volvox *vsr1* mutants caused both male and female germ cell precursors to revert to a vegetative reproductive program, but did not impact the early sexual embryogenesis patterning. The added complexity of Volvox sex-related gene expression provides an opportunity to understand how the core gametogenesis and mating type programs coevolved with multicellularity and sex chromosomes.

Direct targets of VSR1 homodimers or MID-VSR1 heterodimers are currently unknown. Molecular analyses of target genes and sequence binding specificity of VSR1 homodimers and MID-VSR1 heterodimers in Chlamydomonas and Volvox will be useful to understand whether the core recognition sequence changed during the transition to oogamy in Volvox and whether the MID VSR1 protein interaction coevolved as well. It is notable that Volvox MID did not function in Chlamydomonas (44). Reciprocally, Chlamydomonas MID or chimeras between the N terminal and C terminal domains of Chlamydomonas and Volvox MID did not function in Volvox (9). We speculate that part of this incompatibility may be due to inability of MID and VSR1 to undergo cross species interactions, though there may also be changes in DNA binding specificity for their respective RWP-RK domains. Notably, *Gonium* and *Pleodorina* MID proteins showed partial function in Volvox, so MID in these species is predicted to retain some interaction with Volvox VSR1 (44). A second aspect of VSR1 regulation that must have changed during the evolution of the Volvox lineage is the cue for VSR1 expression changing from nitrogen starvation to sex inducer. Little is known about the perception and signaling mechanism of sex inducer, but it likely involves activation of one or more TFs that control expression of VSR1 and other sex-inducer dependent genes.

## Materials and Methods

### *V. carteri and C. reinhardtii* strains and culture conditions

*Eve* (*Volvox carteri.* f. *nagariensis* UTEX 1885) and *AichiM* (*Volvox carteri.* f. *nagariensis* NIES 398) were obtained from stock centers http://web.biosci.utexas.edu/utex/ and http://mcc.nies.go.jp/, respectively. All *Volvox* strains were cultured in Standard Volvox Medium (SVM) at 32°C (unless otherwise specified) and synchronized on a 48 hour developmental cycle with a 16hr:8hr light:dark regime (45) and with a combination of 100 _μ_E blue (465 nm) and 100 _μ_E red (465 nm) LED lights with bubbling aeration. Female *VcMID-GO* transgenic lines were grown at 28°C for vegetative cultures and 32°C to obtain sexual cultures. Sexual development was induced by adding pre-titered sex inducer to juveniles 36 hours prior to embryonic cleavage. Life-cycle cultures including vegetative phase and sexual phase for deep sequencing were collected according to the marked time points in Figure S1. Sexual induction and mating tests were done as described previously (9, 44).

The *Chlamydomonas reinhardtii* strains used in this study were obtained from the Chlamydomonas Resource Center (http://www.chlamycollection.org/): CC-3712 (*mid* deletion mutant), CC-1690 (21gr wild-type *MT*^+^), CC-1691 (6145C wild-type *MT^−^*), and CLiP strains from the Chlamydomonas Library Project (21): CC-5325 (*MT^−^*, parental line of CLiP mutant), CC-5155 (*MT*^+^, isogenic to CC-5325), *rwp4 MT^−^* mutant (LMJ.RY0402.060132), and *rwp11* (*vsr1*) *MT^−^* mutant (LMJ.RY0402.189640). Strains were grown in liquid Tris-acetate-phosphate medium at ∼25°C or on TAP plates with 1.5% agar (46).

### Gene IDs, gene structure and annotations

*C. reinhardtii* and *V. carteri* gene IDs are from Phytozome V13 (47). *RWP4* and *RWP11* orthologs from other Chlamydomonas species (20) with NCBI accession numbers (*CiRWP4*: KAG2440096; *CiRWP11*: KAG2440097; *CsRWP4*: KAG2454850; *CsRWP11*:KAG2454851). VSR1/RWP2 genes from other volvocine species in Figure S2 are available from NCBI (6, 20) (*E*. *debaryana EdaRWP*: KAG2495229; *Y*. *unicocca Yu.g2625.t1*: XXXXXX; *Eudorina* sp. *Eu.g6369.t1*: XXXXXX).

### Cloning *VcVSR1* genomic DNA, *VcVSR1* cDNA and *CrVSR1* genomic DNA

The expression vector construction procedures are described in Supplementary information text 1.

### Transformation, gametogenesis, whole cell extract preparation and immunoblotting of Chlamydomonas extracts

Transformation of Chlamydomonas and preparation of gametes was done as described previously (44). Whole cell extract preparation and immunoblotting of Chlamydomonas cultures were performed as described previously (46, 48) with the following modifications: Antibody dilutions were: rat-anti-HA (1:5000) (3F10, Roche, Switzerland), Anti FLAG rabbit polyclonal (Rockland 600-401-383) was used at 1:5,000 to detect the FLAG epitope tag. Signal was detected by chemiluminescence (Luminata Forte Western HRP Substrate, Millipore) using an Azure c300 Gel Imaging System (Azure Biosystems chemiluminescence).

### *V. carteri vsr1* mutant generation by CRISPR-Cas9 and rescue

CRISPR-Cas9 editing to make *vsr1* mutants was done as described previously using an *in vivo* expression system that co expresses Cas9 enzyme along with a guide RNA (gRNA) in a plasmid vector (23). Oligos used to generate gRNA sequences and their insertion into the expression vector are described in Table S6. All transformations of Volvox were done as described previously (9) with 10 _μ_g/ml Hygromycin B used for selection of edited transformants. The AichiM *vsr1* mutant candidates from the Hygromycin B selection medium were tested for sexual differentiation, and multiple independent candidates were found that did not develop sperm packets and were characterized further by amplifying the edited region using PCR (Table S7) and sequencing. Edited male mutant line *vsr1*^#5_4^ (*vsr1-1*) was used for rescue experiments. Construct pSF-VSR1 was co transformed into *vsr1-1* along with pPmr3 which encodes a paromomycin resistance marker (49) with selection of rescued strains in medium containing 10 _μ_g/ml of paromomycin. Eve *vsr1-1* mutants were generated by crossing Eve and a selected rescued male strain *vsr1::SF-VSR1*. Around 70 meiotic progeny were scored by genotyping (sex determining region, *VSR1* locus, and presence/absence of pSF-VSR1 construct) and phenotypic analysis of sexual development. Genotyping *vsr1-1* was facilitated by use of a *Hpy1*88I restriction site polymorphism created by the edited mutation (Figure S10, Table S4). Strains for Co-IP experiments with MID and VSR1 were generated by transforming either wild-type Eve or rescued female strain *vsr1::SF-VSR1*#1 with plasmid VcMID-GO and selecting for paromomycin resistance carried by the plasmid. Pseudo-male transformants were identified by sexual induction and scoring for sperm packet formation as described previously (9), and MID expression was confirmed by immunoblotting. Selected female VcMID-GO expressing and female *vsr1::SF-VSR1* VcMID-GO expressing transformants were used for Co-IP experiments.

### Chlamydomonas RNA extraction and cDNA synthesis

Vegetative and gamete RNA isolation were done as previously described (50) with the following modifications: *C. reinhardtii* cultures were grown to confluence on TAP plates (50, 51) for four days under continuous light at room temperature. Cells were washed off of the plates with TAP medium and placed immediately into 200ml TAP media (for vegetative samples) with density of ∼5.0x10^6^ cells/mL at 25°C℃ for 4 hours with air bubbling Erlenmeyer, or with nitrogen-free (N-free) HSM media and placed immediately into 200ml nitrogen-free (N-free) HSM media with density of ∼5.0x10^6^ cells/mL at 25℃ for 4 hours in large unshaken Erlenmeyer flasks. For each sample, 200 mL of cells were collected in 250mL centrifuge bottles and Tween-20 was added to a final concentration of 0.005%. The samples were centrifuged at 3,000x*g* for 3 minutes, the supernatant decanted and pellet was resuspended in 1ml TAP or N-free HSM medium, and the suspension was moved to 1.5ml Eppendorf tubes with 5x10^7^ cells/tube. Suspensions were centrifuged at 3,000x*g* for 3 minutes, supernatants removed, and cell pellets were snap frozen in liquid nitrogen and stored at -80C. RNA was extracted from frozen pellets with Trizol (Invitrogen, Carlsbad CA) according to the manufacturer’s protocol. RNA was further purified using RNEasy columns with DNAse digestion (Qiagen) according to the manufacturer’s protocol. cDNA preparation was performed as described previously (44).

### Volvox PCR genotyping, RNA preparation and cDNA synthesis

PCR genotyping, PCR amplification conditions, RNA and cDNA preparation and RT-PCR on *V. carteri* strains were performed as described previously (9, 22) using primers listed in Table S7. RNA was purified using RNEasy columns (Qiagen) according to the manufacturer’s protocol.

### Quantitative RT-PCR (qRT-PCR)

cDNAs were diluted 1:20 in H_2_O. Each 10 µL RT-PCR contained 2.5 µL of diluted cDNA, 1 µL 10 X Choice Taq buffer (Denville Scientific, Cat# C775Y44), 1X SYBR Green I (Invitrogen), 0.6 µL MgCl_2_ (50mM MgCl_2,_ Invitrogen), 1 µL dNTP mix (2mM each dNTP, OMEGA BIO-TEK, Cat#101414-958), 1U Choice Taq (Denville Scientific, Cat# C775Y44), 0.8 µL primers (mix with 5µM each primer). Real-time RT-PCR analysis was carried out using a LightCycler-® 480 Instrument II (Roche Diagnostics, USA) with the following cycling conditions: 95C 3”, 57C 20’, 72C 20’ for 50 cycles. Melt curves and gel electrophoresis were used to confirm the presence of a single amplification product of the correct size in each reaction. For all primer sets, a standard dilution curve was prepared using cDNAs pooled from all samples (Table S7). Relative cDNA levels were calculated using the best-fit curve from the standard dilution of each primer set using supplied software from Roche and then normalized against the internal reference gene cDNA signal. Each reaction was run with three biological and two technical replicates.

### Chlamydomonas immunoprecipitation (IP)

An outcrossed *vsr1-1 MT*^−^ CLiP mutant (LMJ.RY0402.189640) expressing HA-VSR1 was crossed with *MT^+^* wild type strain 21gr to generate a *MT^+^* progeny expressing HA-VSR1. This progeny strain was then crossed with CC-3712 *MT^−^ mid-2::CrMID-6XFLAG* (44) to generate a double tagged strain *MT^−^ mid-2:: HA-VSR1::CrMID-6XFLAG* for Immunoprecipitation (IP) along with the rescued *mid-2 CrMID-6XFLAG* parental strain used as a negative control. Gametes of Chlamydomonas double tagged strain *vsr1*::HA-VSR1 MID-FLAG and control strain CC 3712::MID-FLAG were collected and used to prepare whole cell lysates that were subjected to IP with anti-HA magnetic beads (Pierce™ 88837) as described previously (44, 48, 52). Bound proteins were eluted with 2xSDS sample buffer by boiling at 95℃ for 5 min, separated by SDS PAGE, and detected by immunoblotting as described above.

### Volvox extract preparation and immunoblotting

Preparation of whole-cell lysates and immunoblotting of *V. carteri* cultures were performed as described previously (9) with the following modifications: culture pellets were resuspended in PBS with protease and phosphatase inhibitors (P9599, Sigma-Aldrich; 10 mM PMSF, 10 mM benzamidine, 5 mM EDTA, 5 mM EGTA, 50 mM MG-132, 10 mM ALLN, 1 mM NaF, and 1 mM Na2VO4). Anti-FLAG rabbit polyclonal (Rockland 600-401-383) was used at 1:5,000 to detect the FLAG epitope tag. OLLAS epitope tag antibody (L2) (Novus Biologicals LLC, NBP1-06713) was used at 1:3,000 to detect the OLLAS epitope tag. Antigens were detected by chemiluminescence (Luminata Forte Western HRP Substrate, Millipore) using an Azure c300 Gel Imaging System (Azure Biosystems chemiluminescence) with a Chem Mode.

### Volvox Immunoprecipitation (IP)

Two pseudo-male strains of Volvox (females expressing MID GO) were generated–one coexpressing a pSF-VSR1 tagged transgene in a *vsr1* mutant background, and a control strain expressing only MID-GO in a *VSR1* wild-type background. Upon sexual induction both strains produced sperm packets. Whole cell lysates were prepared from mature sexually induced spheroids with sperm packets and were subjected to IP with anti-FLAG magnetic agarose beads (Pierce,A36797) using a similar procedure as the Co-IP done in Chlamydomonas. Bound proteins were eluted, separated by SDS-PAGE, and detected by immunoblotting as described above.

### Immunofluorescent Staining and Microscopy

Immunofluorescent staining and microscopy on sperm packets were performed as described previously (9) with the following modifications: male *vsr1::SF-VSR1* or wild type male control samples were processed in parallel and imaged under identical conditions. After fixation, blocking and washing, cover slips with adhered spheroids were incubated overnight in anti-StrepII 1:500 (Abcam, ab76949).

### Phylogenomic analysis

Alignments of RWP-RK domains from all predicted family members in *C. reinhardtii* and *V. carteri* plus MID proteins from additional volvocine species in Figure 1 were done using the MUSCLE algorithm in MEGAX (53, 54). MEGAX was used to identify the best fit model, LG+G with gamma shape 1.34. An unrooted maximum likelihood phylogeny was estimated using PhyML with aLRT values calculated for branch support. The resulting tree was converted to a cladogram and a subtree containing RWP-RK members that were sex or mating related was used to create Figure 1C. Estimates of sex induced gene expression were derived from previously published data from Chlamydomonas (16, 17) and from unpublished Volvox transcriptome data shown in Figure 1E and Figure S1. Amino acid similarity between *VSR1* paralogs of core *Reinhardtinia* (Dataset S1) were done with the PLOTCON software contained in the EMBOSS package (55). *Yamagishiella unicocca* and *Eudorina* sp. VSR1 homologs were generated by WebAUGUSTUS (56) based on the published genomes (6).

### Yeast two hybrid assay

Plasmids for yeast two-hybrid experiments were constructed using NEBuilder DNA assembly (NEB), Ligation High kit (TOYOBO), or T4 DNA ligase (NEB) to assemble PCR and restriction fragments as summarized in Table S7 and Dataset S2. The pGBKT7 or pGADT7 backbone vectors were linearized with EcoRI/SalI or EcoRI/XhoI, respectively. Required fragments were amplified with the KOD FX Neo polymerase (TOYOBO) following the manufacturer’s instruction. Isolated colonies were inoculated and mini-prepped with the plasmid mini kit (QIAGEN). Inserted fragments were sequenced and confirmed (GENEWIZ). Yeast two-hybrid experiments were conducted using Matchmaker Yeast Two-Hybrid Kit (Clontech) following the product instruction. *Saccharomyces cerevisiae* strain AH109 was co transformed with bait and prey plasmids and selected on synthetic complete (SC) medium agar plates without leucine or tryptophan (-LW). Resultant colonies were assayed in serial dilution for production of adenine and histidine to grow on SC plates without adenine/histidine/leucine/tryptophan (-AHLW).

## Acknowledgments

We thank Ms. Masumi Taniguchi for maintaining Chlamydomonas strains.

This work was supported by Grants-in-Aid for Scientific Research on Innovative Areas (Grant No. 17H05840 to T.H. and Y.N.) and Scientific Research (C) (Grant No. 20K06766 to T.H.) from the Ministry of Education, Culture, Sports, Science and Technology (MEXT)/JSPS KAKENHI, and National Science Foundation Grant IOS 1755430 to J.U.

## References

1. L. W. Beukeboom, N. Perrin, The Evolution of Sex Determination (Oxford University Press, 2014).

2. J. Heitman, Evolution of sexual reproduction: a view from the Fungal Kingdom supports an evolutionary epoch with sex before sexes. Fungal Biol. Rev. 29, 108-117 (2015).

3. J. Umen, S. Coelho, Algal Sex Determination and the Evolution of Anisogamy. Annu. Rev. Microbiol. 73, 267-291 (2019).

4. S. M. Coelho, J. Gueno, A. P. Lipinska, J. M. Cock, J. G. Umen, UV Chromosomes and Haploid Sexual Systems. Trends Plant Sci. 23, 794-807 (2018).

5. P. Ferris, et al., Evolution of an expanded sex-determining locus in Volvox. Science 328, 351-354 (2010).

6. T. Hamaji, et al., Anisogamy evolved with a reduced sex-determining region in volvocine green algae. Commun. Biol. 1, 17 (2018).

7. P. J. Ferris, U. W. Goodenough, Mating Type in Chlamydomonas Is Specified by mid, the Minus-Dominance Gene. Genetics 146, 859-869 (1997).

8. W. T. Ebersold, Chlamydomonas reinhardi: heterozygous diploid strains. Science 157, 447-449 (1967).

9. S. Geng, P. De Hoff, J. G. Umen, Evolution of sexes from an ancestral mating-type specification pathway. PLoS Biol 12, e1001904 (2014).

10. P. J. Ferris, et al., Plus and minus sexual agglutinins from Chlamydomonas reinhardtii. Plant Cell 17, 597–615 (2005).

11. V. Kurvari, N. V. Grishin, W. J. Snell, A gamete-specific, sex-limited homeodomain protein in Chlamydomonas. J. Cell Biol. 143, 1971–1980 (1998).

12. J.-H. Lee, H. Lin, S. Joo, U. Goodenough, Early sexual origins of homeoprotein heterodimerization and evolution of the plant KNOX/BELL family. Cell 133, 829–840 (2008).

13. C. L. Forest, R. K. Togasaki, Selection for conditional gametogenesis in Chlamydomonas reinhardi. Proc. Natl. Acad. Sci. U. S. A. 72, 3652–3655 (1975).

14. R. E. Galloway, U. W. Goodenough, Genetic analysis of mating locus linked mutations in Chlamydomonas reinhardii. Genetics 111, 447–461 (1985).

15. T. Saito, Y. Matsuda, Isolation and characterization of Chlamydomonas temperature sensitive mutants affecting gametic differentiation under nitrogen-starved conditions. Curr. Genet. 19, 65–71 (1991).

16. S. Joo, et al., Gene Regulatory Networks for the Haploid-to-Diploid Transition of Chlamydomonas reinhardtii. Plant Physiol. 175, 314–332 (2017).

17. D. Lopez, et al., Dynamic changes in the transcriptome and methylome of Chlamydomonas reinhardtii throughout its life cycle. Plant Physiol. 169, 2730–2743 (2015).

18. S. Schmollinger, et al., Nitrogen-sparing mechanisms in Chlamydomonas affect the transcriptome, the proteome, and photosynthetic metabolism. Plant Cell 26, 1410–1435 (2014).

19. U. Goodenough, et al., The path to triacylglyceride obesity in the sta6 strain of Chlamydomonas reinhardtii. Eukaryot. Cell 13, 591–613 (2014).

20. R. J. Craig, A. R. Hasan, R. W. Ness, P. D. Keightley, Comparative genomics of Chlamydomonas. Plant Cell 33, 1016–1041 (2021).

21. X. Li, et al., A genome-wide algal mutant library and functional screen identifies genes required for eukaryotic photosynthesis. Nat. Genet. 51, 627–635 (2019).

22. C. López-Paz, D. Liu, S. Geng, J. G. Umen, Identification of Chlamydomonas reinhardtii endogenous genic flanking sequences for improved transgene expression. Plant J. 92, 1232– 1244 (2017).

23. J. A. Ortega-Escalante, R. Jasper, S. M. Miller, CRISPR/Cas9 mutagenesis in Volvox carteri. Plant J. 97, 661–672 (2019).

24. N. Ueki, I. Nishii, Controlled enlargement of the glycoprotein vesicle surrounding a volvox embryo requires the InvB nucleotide-sugar transporter and is required for normal morphogenesis. Plant Cell 21, 1166–1181 (2009).

25. U. W. Goodenough, C. Hwang, A. J. Warren, Sex-limited expression of gene Loci controlling flagellar membrane agglutination in the chlamydomonas mating reaction. Genetics 89, 235– 243 (1978).

26. Y. Sakuraba, M. Zhuo, S. Yanagisawa, RWP-RK domain-containing transcription factors in the Viridiplantae: their biology and phylogenetic relationships. J. Exp. Bot. 73, 4323–4337 (2022).

27. S. C. Lee, M. Ni, W. Li, C. Shertz, J. Heitman, The evolution of sex: a perspective from the fungal kingdom. Microbiol. Mol. Biol. Rev. 74, 298–340 (2010).

28. A. D. Johnson, Molecular mechanisms of cell-type determination in budding yeast. Curr. Opin. Genet. Dev. 5, 552–558 (1995).

29. R. Sekido, R. Lovell-Badge, Sex determination involves synergistic action of SRY and SF1 on a specific Sox9 enhancer. Nature 453, 930–934 (2008).

30. T. Yamazaki, et al., Genomic structure and evolution of the mating type locus in the green seaweed Ulva partita. Sci. Rep. 7, 11679 (2017).

31. X. Liu, et al., Transcriptional dynamics of gametogenesis in the green seaweed Ulva mutabilis identifies an RWP-RK transcription factor linked to reproduction. BMC Plant Biol. 22, 19 (2022).

32. R. Blanc-Mathieu, et al., Population genomics of picophytoplankton unveils novel chromosome hypervariability. Sci. Adv. 3 (2017).

33. A. Z. Worden, et al., Green evolution and dynamic adaptations revealed by genomes of the marine picoeukaryotes Micromonas. Science 324, 268–272 (2009).

34. H. Sekimoto, et al., A divergent RWP-RK transcription factor determines mating type in heterothallic Closterium. New Phytol. 237, 1636–1651 (2023).

35. D. Koszegi, et al., Members of the RKD transcription factor family induce an egg cell-like gene expression program. Plant J. 67, 280–291 (2011).

36. F. Tedeschi, P. Rizzo, T. Rutten, L. Altschmied, H. Bäumlein, RWP-RK domain-containing transcription factors control cell differentiation during female gametophyte development in Arabidopsis. New Phytol. 213, 1909–1924 (2017).

37. T. Waki, T. Hiki, R. Watanabe, T. Hashimoto, K. Nakajima, The Arabidopsis RWP-RK protein RKD4 triggers gene expression and pattern formation in early embryogenesis. Curr. Biol. 21, 1277–1281 (2011).

38. S. Koi, et al., An evolutionarily conserved plant RKD factor controls germ cell differentiation. Curr. Biol. 26, 1775–1781 (2016).

39. M. Rövekamp, J. L. Bowman, U. Grossniklaus, Marchantia MpRKD regulates the gametophyte-sporophyte transition by keeping egg cells quiescent in the absence of fertilization. Curr. Biol. 26, 1782–1789 (2016).

40. C. Chardin, T. Girin, F. Roudier, C. Meyer, A. Krapp, The plant RWP-RK transcription factors: key regulators of nitrogen responses and of gametophyte development. J. Exp. Bot. 65, 5577–5587 (2014).

41. P. J. Ferris, C. Pavlovic, S. Fabry, U. W. Goodenough, Rapid evolution of sex-related genes in1Chlamydomonas. Proc. Natl. Acad. Sci. U.S.A 94, 8634–8639 (1997).

42. J. A. Fraser, J. Heitman, Fungal mating-type loci. Curr. Biol. 13, R792–R795 (2003).

43. L. W. Beukeboom, N. Perrin, “The evolution of sex chromosomes” in The Evolution of Sex Determination, (Oxford University Press, 2014), pp. 89–114.

44. S. Geng, A. Miyagi, J. G. Umen, Evolutionary divergence of the sex-determining gene uncoupled from the transition to anisogamy in volvocine algae. Development 145 (2018).

45. D. L. Kirk, M. M. Kirk, Protein synthetic patterns during the asexual life cycle of Volvox carteri. Dev. Biol. 96, 493–506 (1983).

46. E. H. Harris, The Chlamydomonas Sourcebook: Introduction to Chlamydomonas and Its Laboratory Use (Academic Press, 2008).

47. D. M. Goodstein, et al., Phytozome: a comparative platform for green plant genomics. Nucleic Acids Res. 40, D1178–86 (2012).

48. Y. Li, D. Liu, C. López-Paz, B. J. Olson, J. G. Umen, A new class of cyclin dependent kinase in Chlamydomonas is required for coupling cell size to cell division. Elife 5, e10767 (2016).

49. T. Jakobiak, et al., The bacterial paromomycin resistance gene, aphH, as a dominant selectable marker in Volvox carteri. Protist 155, 381–393 (2004).

50. P. L. De Hoff, et al., Species and population level molecular profiling reveals cryptic recombination and emergent asymmetry in the dimorphic mating locus of C. reinhardtii. PLoS Genetics 9, e1003724 (2013).

51. E. H. Harris, The Chlamydomonas Sourcebook: A Comprehensive Guide to Biology and Laboratory Use (Elsevier, 2013).

52. B. J. S. C. Olson, et al., Regulation of the Chlamydomonas cell cycle by a stable, chromatin-associated retinoblastoma tumor suppressor complex. Plant Cell 22, 3331–3347 (2010).

53. S. Kumar, G. Stecher, M. Li, C. Knyaz, K. Tamura, MEGA X: Molecular Evolutionary Genetics Analysis across Computing Platforms. Mol. Biol. Evol. 35, 1547–1549 (2018).

54. R. C. Edgar, MUSCLE: multiple sequence alignment with high accuracy and high throughput. Nucleic Acids Res. 32, 1792–1797 (2004).

55. P. Rice, I. Longden, A. Bleasby, EMBOSS: the European Molecular Biology Open Software Suite. Trends Genet. 16, 276–277 (2000).

56. K. J. Hoff, M. Stanke, WebAUGUSTUS—a web service for training AUGUSTUS and predicting genes in eukaryotes. Nucleic Acids Research 41, W123–W128 (2013).

