## Supplementary material for "A conserved RWP-RK transcription factor VSR1 controls gametic differentiation in volvocine algae": Dataset S1

```
CLUSTAL W 2.0 multiple sequence alignment

CrRWP11             ---------------------------------------------------------------------- 70
CiRWP11             ---------------------------------------------------------------------- 70
CsRWP11             ---------------------------------------------------------------------- 70
CrRWP4              ---------------------------------------------------------------------- 70
CiRWP4              ---------------------------------------------------------------------- 70
CsRWP4              ---------------------------------------------------------------------- 70
EdaRWP/KAG2495229   ---------------------------------------------------------------------- 70
Yu.g2625.t1         ---------------------------------------------------------------------- 70
Eu.g6369.t1         ---------------------------------------------------------------------- 70
VcRWP2              MNTGKGLKARRKFHGLPNSLMRRASKTWLGPNKCTSKPAGRGTLLAPSLYFYYVDQFRQWFTEYEPPSLK 70


CrRWP11             -------------------MDLDVPDLLADFNPGAGLVVPIASVGQALLPLARNLPKEE----------- 140
CiRWP11             -------------------MDPDVCELLAEFNPGAGRVAPIASVGQALLPLARKIPEED----------- 140
CsRWP11             -------------------MDPDVCELLAEFNPGAGRVAPIASVGQALLPLARKIPEED----------- 140
CrRWP4              ---------------------------------------------------------------------- 140
CiRWP4              ---------------------------------------------------------------------- 140
CsRWP4              ---------------------------------------------------------------------- 140
EdaRWP/KAG2495229   ----------------MAAVPLELASILEDFHPTDGKVQALEPVGQDQLPLARE---------------- 140
Yu.g2625.t1         ---------------------MSYGGHNHHHQPSAALGTHSATRQGLDLLMLR----------------- 140
Eu.g6369.t1         ---------------------------------------------------------------------- 140
VcRWP2              GTTLQEPLCSARRRRACRRLMEDLAEVLTEFHPSTGKVPHLDTLSQAHLPIARELPNNDSLRLEEYLSAG 140


CrRWP11             ---------------------------------------------------------------------- 210
CiRWP11             ---------------------------------------------------------------------- 210
CsRWP11             ---------------------------------------------------------------------- 210
CrRWP4              ---------------------------------------------------------------------- 210
CiRWP4              ---------------------------------------------------------------------- 210
CsRWP4              ---------------------------------------------------------------------- 210
EdaRWP/KAG2495229   ---------------------------------------------------------------------- 210
Yu.g2625.t1         ---------------------------------------------------------------------- 210
Eu.g6369.t1         ---------------------------------------------------------------------- 210
VcRWP2              SFAVLDAALHSICESQATLPETNAGSSAPSSYYHSHQDQNALTGQGTNPMLATRQGFTSGQLAASTTGQT 210


CrRWP11             ----------------LENVDVTGLDALLSSDPSALNDVTFQRSSPSAPTPQATPT-APSTSARPGSSTT 280
CiRWP11             ----------------QHEVDGTGLDALLSSDP-ALADVTLQRYSPSAPTPQATPT-APVTSR---SSQP 280
CsRWP11             ----------------QHEVDGTGLDALLSSDP-ALADVTLQRYSPSAPTPQATPT-APVTSR---SSQP 280
CrRWP4              --------------------MDTGSAAI---------------------AADAAVSKDSNDSQ---VAAE 280
CiRWP4              -------------------MDNVGSG-----------------------DDSAAAATDPRAFK---MSAE 280
CsRWP4              -------------------MDNVGSG-----------------------DDSAAAATDPRAFK---MSAE 280
EdaRWP/KAG2495229   ----------------LSNADSLKLGELFSAG--SLSAVEAALGVVAGPGSEVIAQGAPRSSS---GAEQ 280
Yu.g2625.t1         --------------------QGSGAAASAFAHLAASYAITQQQQQQQQQQQQQQQQQQQQQLQ---QERS 280
Eu.g6369.t1         ----------------MVQAAQSG------------------PPLPTPPPPLQTHQHTLQHHQ---LQHQ 280
VcRWP2              LQQQTQQHQQQQQEHQHRQAADGGQAAASGTGSSGLFPRQMNPQLAQPPTPQQQQQQQPSQHQLQELQHQ 280


CrRWP11             LAAGASVHQPYLR-EQQRQT---------------------------LPHLPSLSGFASAGAGQQPACPA 350
CiRWP11             DGAGASCSQQCCALNQQGTT---------------------------LPHLPSLSTASGAAADQHPACRA 350
CsRWP11             DGAGASCSQQCCALNQQGTT---------------------------LPHLPSLSTASGAAADQHPACRA 350
CrRWP4              IAHP------------------------------------------------------------------ 350
CiRWP4              IVDG------------------------------------------------------------------ 350
CsRWP4              IVDG------------------------------------------------------------------ 350
EdaRWP/KAG2495229   HAHDAMAAKPFSFLDPAGQA---------------------------QPSFHSLSSPIDEDGAPQPSAPA 350
Yu.g2625.t1         EGPAHAHGSGCNHVHFPGQ------------------------------HQQQAQQPQQAQQAQQPPQPF 350
Eu.g6369.t1         LQSGVGHFHHH--HHHHNP-------------------------------------------------LV 350
VcRWP2              HLHQHQHQHQHQQHQQQQQQQLLQGTRTSPAQQPSAPAQQPQQQHSLQNHQQQQQQHQQHQQQKRPSLAS 350


CrRWP11             GSH---------LGPAAALGGTQACLSKISDPGCGKQTTSNRSSLSQMDEDGGDDAGAPEHEAQDVRNGD 420
CiRWP11             GGKSPRPAVPFGASDAPAVGSVPACLSKISDPGCGSQSTSNRPSLSQMDADGG-DAGRPEDPV-----GS 420
CsRWP11             GGKSPRPAVPFGASDAPAVGSVPACLSKISDPGCGSQSTSNRPSLSQMDADGG-DAGRPEDPV-----GS 420
CrRWP4              -----------------------GC-----------------------------GYQATQKCK------D 420
CiRWP4              -----------------------SC-----------------------------QICGPSQGQ------S 420
CsRWP4              -----------------------SC-----------------------------QICGPSQGQ------S 420
EdaRWP/KAG2495229   RGY--------------------SC-----------------------------DASAAAAQ-------L 420
Yu.g2625.t1         ANA--------------------APIGGLSPSNSNGSNTSSK------------DAAAPALH-------D 420
Eu.g6369.t1         GVA--------------------ACSGCATRSDSNSNSKNGSISSTR-------DAAVPVPHH------E 420
VcRWP2              GGA--------------------ACSRGPSRSESNSNGSNQISR----------DATVPPPH-------E 420
                                           .                                  .

CrRWP11             SGRGDGGGGGHMYASEPP--RGPEPMTLISGCSETASGHRIDGGEAPSVSHSPSGAGGGASAPGAAGGAS 490
CiRWP11             AGDGD----ARMYCSEPPPQQGLAPMAVTAGCSGSDSRQPGTIALVPSASPSPSGSGGAAAA-DTLGGAG 490
```

### Dataset S1

```
CsRWP11              AGDGD----ARMYCSEPPPQQGLAPMAVTAGCSGSDSRQPGTIALVPSASPSPSGSGGAAAA-DTLGGAG 490
CrRWP4               SGTGR---------------------------------------------PGEVDLGGDVKDTQLVKAPA 490
CiRWP4               GGARQ---------------------------------------------PTFTTIGSKVAD---ISQPG 490
CsRWP4               GGARQ---------------------------------------------PTFTTIGSKVAD---ISQPG 490
EdaRWP/KAG2495229    TAAGD---------------NGPATATASAAASERSAGS-----------GGRHGSGSGPAS-GASGGPG 490
Yu.g2625.t1          GSTGE-----------------------SAQRAEDGDGRAV---------GARATEGAAVADTNAAGGPD 490
Eu.g6369.t1          SSTGE---------------SGP---RAELSCMDVPMSRPA---------SNTVVVAAGAAAAAAASAAG 490
VcRWP2               SSTGE---------------SGP---RADSGPMEVQVARAT---------GATAAAVVTTVAAGAAGGGG 490
                      .

CrRWP11              GATRPEQAETAPSRMSANHQLPGADPTAGASPRNGSGIAAADGDAEGSGDGDGDEEEEEEDDEELGDALA 560
CiRWP11              EIKGPESAPGAPSRMSGNNQSSGAKGDGTGSPPQAEGGNMAADQSEDSGDGEGADEEEGEEAPCGEEAAG 560
CsRWP11              EIKGPESAPGAPSRMSGNNQSSGAKGDGTGSPPQAEGGNMAADQSEDSGDGEGADEEEGEEAPCGEEAAG 560
CrRWP4               ---------------------------------------------------------------------- 560
CiRWP4               ---------------------------------------------------------------------- 560
CsRWP4               ---------------------------------------------------------------------- 560
EdaRWP/KAG2495229    ------------------------PGDAGGGGGSSGGPAAAD---------------------------- 560
Yu.g2625.t1          ---------------------------GGGPP-------------------------------------- 560
Eu.g6369.t1          ---------------------------GGGSGIGRGDTAAATGRSSHEAQQPTHQPPPPPQPPPPPPAMP 560
VcRWP2               ---------------------------GSGASGSRGDGHVASGPAAGPAGGASDQTHPPPPPPEVST--- 560


CrRWP11              GAPAEGAVSGSRPAKAAR-SKAQPRGRRPKAAKQDGGGEGGSDSED-EDGGESRHLGRRVIEYAEKIVEQ 630
CiRWP11              GEEAAGG--RSRRAKAARTTKVQAKRRKQKPAKQEVDGDGDSDSDDEEEGGENRHLGRRVIEYADKIVEQ 630
CsRWP11              GEEAAGG--RSRRAKAARTTKVQAKRRKQKPAKQEVDGDGDSDSDDEEEGGENRHLGRRVIEYADKIVEQ 630
CrRWP4               ---------------------------------------------------------------------- 630
CiRWP4               ---------------------------------------------------------------------- 630
CsRWP4               ---------------------------------------------------------------------- 630
EdaRWP/KAG2495229    ---------GSRPGRKGR----------------------EAEEAD-----------------IARVVDA 630
Yu.g2625.t1          ---------VPAESRRLR----------------------EDEQRD-----------------VDEFVNL 630
Eu.g6369.t1          PPAPPDVNTPPAEGRKLR----------------------EEEQRD-----------------VEEFVNL 630
VcRWP2               ---------PPAEGRKLR----------------------EEEQRD-----------------VEEFVNL 630


CrRWP11              AWAANQLSDVPHTATLDCNELPGIGFKKAGRILATAAVRLLVRGEEAQKKGTLRT-LLRSAENAGILAQ- 700
CiRWP11              AWAANQLSEVPHTAKLDCNELPGIGFKKAGRILATAAVRLLVRGEEAHRNGTLRP-LLRTAEDTGLLTQ- 700
CsRWP11              AWAANQLSEVPHTAKLDCNELPGIGFKKAGRILATAAVRLLVRGEEAHRNGTLRP-LLRTAEDTGLLTQ- 700
CrRWP4               --------DAANSAMNTC------GRLSGPKAPGGGGPPLPLKAMKEAGSG------------------- 700
CiRWP4               -----KLEDVAAEMPSTC-------------------SPAVHAETIKDRTG------------------- 700
CsRWP4               -----KLEDVAAEMPSTC-------------------SPAVHAETIKDRTG------------------- 700
EdaRWP/KAG2495229    AMDANYNQGVVPTVAFDG---PYGGPRTTGRLIATTSAAMFRKGVAASSAGATLPKTLDLALRAQIEKHA 700
Yu.g2625.t1          AWEANHKHGESVRLEFTG---PNVGARTTGRLMAAATAQLMRRGKAAAQSG-QIT-GLNSTIRAAMERH- 700
Eu.g6369.t1          AWNANYKQGDAVRLEFTG---PNVGARTTGRLMAAATAQLMRRGVAAAQAG-QIT-GLNTTIRSAMDRH- 700
VcRWP2               AWNANYKQGDAVRLEFTG---PNVGARTTGRLMAAATAQLMRRGVAAAQAG-QIT-GLNTSIRSAMERH- 700
                                                                       *

CrRWP11              --QGGPAGAAGIAAGGGAARGTAGGADMGASAAAAAAAAAAVASMPNLASMDPETLRVCLAAAAAAAAAA 770
CiRWP11              --QQGAGGAAGVEAA------------QTADGAASASAAVAGAGDPKRTGLNSQALRACMAAAATAAAAA 770
CsRWP11              --QQGAGGAAGVEAA------------QTADGAASASAAVAGAGDPKRTGLNSQALRACMAAAATAAAAA 770
CrRWP4               ---------AGLDGE--------------SRVHFLQEAFLTISEELAAQGVNPHAVIA-----AASNSPT 770
CiRWP4               ---------ASLDEE--------------SRMHCLQAAFLTISEELAAQGVNPHAVIAAAAAAAASNSPT 770
CsRWP4               ---------ASLDEE--------------SRMHCLQAAFLTISEELAAQGVNPHAVIAAAAAAAASNSPT 770
EdaRWP/KAG2495229    ANLSGTNHVGPLDKD--------------RRTAFLMEAYLQISEELARSGVD-------IAAAAATAAAA 770
Yu.g2625.t1          --LPAKKEEAHLEGN--------------KRTEFLTEALRQITEQLISAGVDPRTIID------GSAATA 770
Eu.g6369.t1          --LPAKRDKAQLEGD--------------KGTELLTEALRHITEQLVSRGVDPRTIIDSAGNPTASGGAA 770
VcRWP2               --LPVKRDKAHLEGD--------------KGTELLTEALRHITEQLMSRGVDPRSIIE---------NHS 770
                              . :                         *    :      .::                 :

CrRWP11              AGTPSATSGVTQQPQTHAQANAIVNLASELQQHGLGQWAVQAAAEVAMKAAAAAVLQQQQHAAAAAAADP 840
CiRWP11              SASAAASAAAS-------------SSAAHAQQHGLSQWAVQAAAAAAVQVAAAAALQQQQQAAAALAA-- 840
CsRWP11              SASAAASAAAS-------------SSAAHAQQHGLSQWAVQAAAAAAVQVAAAAALQQQQQAAAALAA-- 840
CrRWP4               ASRGASSGGVA----------------------------------AAVAAAAFILPAVRPLSAPTTAAIP 840
CiRWP4               ASRGGSGGV------------------------------------AAVVAAAALISPTRPPASATHAPAA 840
CsRWP4               ASRGGSGGV------------------------------------AAVVAAAALISPTRPPASATHAPAA 840
EdaRWP/KAG2495229    ASSPSAS-------------------------------------------------PNRASASAAVASAF 840
Yu.g2625.t1          NQQPGNNNN------------------------------------------------NNNNNNNNNNNRN 840
Eu.g6369.t1          AAAPSGSAVAA-------------AG--------------GLSPTATTAAAAASGSLNRAHSIGDYSGNM 840
VcRWP2               NGTPPGGGG------------------------------------AATAVAAAAAAANGGAGGSKTSLGS 840


CrRWP11              RGAGSSVDAG---PS------SLNLLTAAAVGPPDMRHFGGFAAAPSHAAA--GGNMATLYSAQGNPRVE 910
CiRWP11              -AAGVGSGGG---PS------SASLFSAAAVGAPSMHGLGGCAPPFPHHQHHTGGDVAMFFPAAQPHSAD 910
CsRWP11              -AAGVGSGGG---PS------SASLFSAAAVGAPSMHGLGGCAPPFPHHQHHTGGDVAMFFPAAQPHSAD 910
CrRWP4               VVMAGGGSGV-----------VPAPLPTMPSLTPSQAQTQSCANAAADAASPTAASGGGGPP-------- 910
CiRWP4               AGSPTSASGA------------PPLPSLTPMPPPPLLANGACSAAATESAASLAVPSEGDTAA------- 910
CsRWP4               AGSPTSASGA------------PPLPSLTPMPPPPLLANGACSAAATESAASLAVPSEGDTAA------- 910
```

### Dataset S1

```
EdaRWP/KAG2495229   AAAAAAAPGG-----------------------------GGMVDP------------------------- 910
Yu.g2625.t1         SGGGGGDAGR------------APLL------LPSLLAQDTPAPQPQSAQPHYAGALEMLQSH------- 910
Eu.g6369.t1         ARKLSSPPAQLTVPSQQQQQQQQPQQQPQQQLQPPMVNAMGLQAHLHHHHQQLAAAMQQQQAQ------- 910
VcRWP2              IGDNSADTAGLKLPS------PPPVAAGAMVGMPPPGGAMDLIQLQQQQQLQQQLQLQMQMQQAP----- 910
                         .  .

CrRWP11             PAAAPGPFQQMVLEAALA-----------DDCNPDGGAGAGLTTGGELECTLSGPRSSTDELLQAVFDGM 980
CiRWP11             AAVVPGPLQQMLLEAVLE--GSTADTPAPAVTAPDAGVGPSLAEVGR---SLSGPRSSTAEMLQAIFDVM 980
CsRWP11             AAVVPGPLQQMLLEAVLE--GSTADTPAPAVTAPDAGVGPSLAEVGR---SLSGPRSSTAEMLQAIFDVM 980
CrRWP4              ------ASAPLLPDGPAA------------PAAPAIGGGNGGGDAAAMGDAAAAAAAGGFDFLFDAFHAF 980
CiRWP4              ------ASPQAQPEGPAAPPPAAAAAAGDNDDGTDGCGGHGNPAGAS---SGAGDAAEGFDFLFDAFHAF 980
CsRWP4              ------ASPQAQPEGPAAPPPAAAAAAGDNDDGTDGCGGHGNPAGAS---SGAGDAAEGFDFLFDAFHAF 980
EdaRWP/KAG2495229   ------PLLQQVLDAAAI--------------SHDGMYGAGGAAG----------GVGGTDILLEALAAF 980
Yu.g2625.t1         ------HFQQQMMDAIRE------------QAGHNGYEADGGFAAH------AGQGGPDPDPLHQAFQAF 980
Eu.g6369.t1         ------PFQQMMLEATRE--GPMAAAAAGTDPRVSGMGYYGCPGGAE---QVPIIGASVSDLLQEAFRAF 980
VcRWP2              ------QFQQLMLEATREHHQPTGPMGSTGGPPLVDYGGFFSPTGETGPVGVPAAGGGVSDILQEAFRAF 980
                                 :.                                             : *   :  :

CrRWP11             GDPGGARALTGNSGGGPSGGSNVVAG------------VAEPRKSDGAAWDLLGQMFDSYCGGGAPAAAT 1050
CiRWP11             GEPGTSAGAMRAGTGSGSGGAGSSGGASGAISGADGGADAQRSDNAAADWDFLGQAFDRYCGAAVAAVPE 1050
CsRWP11             GEPGTSAGAMRAGTGSGSGGAGSSGGASGAISGADGGADAQRSDNAAADWDFLGQAFDRYCGAAVAAVPE 1050
CrRWP4              VHHDGPTADANTVAAEASPASKARPT-------------------------------------------- 1050
CiRWP4              VSHDGPNTAASNVAAAAGGEATAPRK-----------------------------------------LPR 1050
CsRWP4              VSHDGPNTAASNVAAAAGGEATAPRK-----------------------------------------LPR 1050
EdaRWP/KAG2495229   SS------------GAADGGVHA----------------------------------------------- 1050
Yu.g2625.t1         VAE-----------GDAAGGADGRGD-------------------------------------------- 1050
Eu.g6369.t1         ASEQEAAAAAAEAAGANGGGNGDVVV-------------------------------------------- 1050
VcRWP2              AAEQEAAAVAAAAAAAAAEGGDGSGG-------------------------------------------- 1050
                                  .

CrRWP11             DTQASVSLPHRQHHQPLQLP---LPHVSLQHPALAMIAAAEAAGPTGGF-RLNGGSITMLDAWVDAQNDE 1120
CiRWP11             P-QAPISPVRHQLQGQPQPP-RQLPLAPMQHPALAAIAAAEAVAPTGCYGRMHGGSITMLDAWVDAQNDE 1120
CsRWP11             P-QAPISPVRHQLQGQPQPP-RQLPLAPMQHPALAAIAAAEAVAPTGCYGRMHGGSITMLDAWVDAQNDE 1120
CrRWP4              ----SQQPTAVQTADRDQSTCRAAGMQPPPPQSALSGVAT--------AARANGRSVTMFDVWAA----- 1120
CiRWP4              PSQQHQALPAVKTEAEVAVPVGVVHHRPPLHPVLSAVVAA--------VTRPNGGSVTMLDAWAA----- 1120
CsRWP4              PSQQHQALPAVKTEAEVAVPVGVVHHRPPLHPVLSAVVAA--------VTRPNGGSVTMLDAWAA----- 1120
EdaRWP/KAG2495229   -------LQDFTHAAAAAAA-SAAAAAAAAAAAGLTFPVP--------YNRTT-GSVTMLDAWVD----- 1120
Yu.g2625.t1         --SAHVRMMDFNDMQATELP-RSLPAAPLLHPTAAALAVP--------YQRAT-GSVTMIDVWVN----- 1120
Eu.g6369.t1         ---VNASMMDYHAATIAGAGLSGTAMMP--------------------YHRTA-GSVTMIDVNWE----- 1120
VcRWP2              DMGATATMMDFQAAAAAAAS-SVLPNAATAAVPL--------------YQRTT-CSVTMIDVGHD----- 1120
                                               .                      *    *:**:*.

CrRWP11             HVDAAVVAAMVAG---DDDVPRLHDTLGPVVGGAATAACADGAGLSTASRQHQH-QHPQQQA--APTPTL 1190
CiRWP11             HVDAAVVADMVGGGDDDDDLPRLHETLGPAAAAAVARGAAPRAAGAVGIPSTPSGAQPQLQPAVAAGPTL 1190
CsRWP11             HVDAAVVADMVGGGDDDDDLPRLHETLGPAAAAAVARGAAPRAAGAVGIPSTPSGAQPQLQPAVAAGPTL 1190
CrRWP4              ----------------GEE---------GGSDGGNGAPATDTDPIQQPSAA------PPTAE--GDGTAQ 1190
CiRWP4              ----------------GDD--------------------ESGGQDQTEPPE------PTLVA--AGGQAD 1190
CsRWP4              ----------------GDD--------------------ESGGQDQTEPPE------PTLVA--AGGQAD 1190
EdaRWP/KAG2495229   ----------------DTDVTSPAKPLPGAAAGGSAATAATSGGGAAPAPS------TSANG--TSSSSD 1190
Yu.g2625.t1         ----------------DNDGILPPMPPAAAAADANANSYAAAGGSTAPHPQ------PINFS--SLGPGA 1190
Eu.g6369.t1         ----------------NTE--------QHAAAAAVAAAAAAAGANAAAADA------EAQAAASAQHAAG 1190
VcRWP2              ----------------ESELSSPH----AAAAAAAAAAAAAAAGGTLLSEG--------MQG--ACGPPG 1190
                                      :

CrRWP11             LAPMPVRGGGGAGVSALADVSPGDHSPGSDGGGLMAPPQPRPPICI---SPPPEDLDPSE---------- 1260
CiRWP11             HAPVPFRGGIVSAFAAVSPTAEHSPGSGDGSGGSMGPPAPRAPICI---SPPPEDLDPTE---------- 1260
CsRWP11             HAPVPFRGGIVSAFAAVSPTAEHSPGSGDGSGGSMGPPAPRAPICI---SPPPEDLDPTE---------- 1260
CrRWP4              QQQADSHGGAAAGSTAVEAIAG-----------------ASSKTLA---APKPVRLSPFS---------- 1260
CiRWP4              PS---SAGATAAGRNRAPLAAP-----------------KALHRCP---LPLPAATAASG---------- 1260
CsRWP4              PS---SAGATAAGRNRAPLAAP-----------------KALHRCP---LPLPAATAASG---------- 1260
EdaRWP/KAG2495229   VSNGAWQGTAAATVASASAVSA--------------APAAAAPLTP---AAPLPLLSPTP---------- 1260
Yu.g2625.t1         SS--PW-GGGGGVPGPQPGLQQ--------------PPSSSSPLQPPSASPPARLVHPRP---------- 1260
Eu.g6369.t1         FGAGSWGCGRGAGGKTVPP-----------------QHLQQQQQQPPHQQQQQHLHHPTPTSAAGGGGRG 1260
VcRWP2              FRSGQWIPTGRSMVGGKAATLPSSSSSLQTTASAAAAAAAAAAAALPHQQQQQMLHHSAA---------- 1260
                               .                                             .

CrRWP11             ----LRDDFPDLPLTASMFRISNMSIADGP---------PAPGSARPSASAFGA-NPFGQFGAA--GGVG 1330
CiRWP11             ----LRNEFQDLPLTASMFRISNMSIADVP---------PAPGTARPSATAFGANNPFGSFGAL-PGGGG 1330
CsRWP11             ----LRNEFQDLPLTASMFRISNMSIADVP---------PAPGTARPSATAFGANNPFGSFGAL-PGGGG 1330
CrRWP4              ----GGG------GAVSAFAGPATTCISPAPMDAMD-GSDGDGIGRGDSEGGKGFGGASRRASRRGGGAA 1330
CiRWP4              ----AAT------GVVSAFASPPPTCVSPAPLDAMDAGSGGEGLGRSDGEGYGGYGGSESRRASRRGGGG 1330
CsRWP4              ----AAT------GVVSAFASPPPTCVSPAPLDAMDAGSGGEGLGRSDGEGYGGYGGSESRRASRRGGGG 1330
EdaRWP/KAG2495229   ----QRP------GVFSPFGAAGA-CDSPP---------PIEGHLGPDAPCNRGGMGVSPRPSN---VVT 1330
Yu.g2625.t1         ----SV-------GMVSAFAAPG--CISPQALDDADQ--QHPAHERLASPP----------------RQG 1330
Eu.g6369.t1         SGGGGGGTSGGGNGMMSAFAGRAT-CISPQL------GAPEGCDGRPLSPV-----------------HT 1330
VcRWP2              ----AGCAAG---GMMSSFGGPPT-CISPPP-------GPETSDKRPLSPS-----------------RF 1330
```

Dataset S1

```
                                    * *        .                    .

CrRWP11             GGGT-----PNRLSNALSILR---AGS-AGCMDMLMSND-FMDALAAADPLLAAEVSAGGAAGGGGRASH 1400
CiRWP11             GGGT-----PIRLSNAMSILR---AGS-AGCMDMLMSND-FMDTLAATDPLLAAEV---GGAAGGGRPSH 1400
CsRWP11             GGGT-----PIRLSNAMSILR---AGS-AGCMDMLMSND-FMDTLAATDPLLAAEV---GGAAGGGRPSH 1400
CrRWP4              GGAT-----PMRLSNALSLLGAASAG--GGYMDLLGPGDMFMDS--GVDPLADILM-------DAMRPSL 1400
CiRWP4              GGGTGGGATPMRLSNALSLLRAASAG--TAYMDLLGPGDLFMDG--GVDPLADIMM-------DAMRPSL 1400
CsRWP4              GGGTGGGATPMRLSNALSLLRAASAG--TAYMDLLGPGDLFMDG--GVDPLADIMM-------DAMRPSL 1400
EdaRWP/KAG2495229   GGTS-----PLRLSNAMSILR---AGSMGGYMDIMTSNDFFMD-----DLHL-----------DGRLPSL 1400
Yu.g2625.t1         GACT-----PLRLSNAMSILRQASAGIGGAYIDMVLSTDFIFD---GHEGLL-----------DGRLPSI 1400
Eu.g6369.t1         EGPS-----PHRLSNAMSILRNPSSGLGPGDLDIIM-GNYFLD---GADPLLS----------DGRLPSI 1400
VcRWP2              DTGS-----PHRLSNAMSILRNRSAD--LGSLDMIMSNDYFMD---GVDPLLN----------DGRQPSI 1400
                       :     * *****:*:*    :.   . :*::   : ::*     :              ..  .*

CrRWP11             ADKFLMSIDALPPVPPLLPPSLIPPGYA--------------NALVMTTGDAGAGLSNSGPLSAAAGAA- 1470
CiRWP11             ADKFLMCIDALPPMPPLLPSSLIPPGYA--------------NALIMTTGDAGAGMSYSSGGGAMAVAA- 1470
CsRWP11             ADKFLMCIDALPPMPPLLPSSLIPPGYA--------------NALIMTTGDAGAGMSYSSGGGAMAVAA- 1470
CrRWP4              TEIPVVAMPAAAAAAAAA------AGGA----------------AAEPAGRSGVPRSSLATLNRMLMSE- 1470
CiRWP4              TDIPVTTAAALTAAPAPVPASA--AGGAAA------------APAAAELSTSGGGRSGGGSGSRSSLTTL 1470
CsRWP4              TDIPVTTAAALTAAPAPVPASA--AGGAAA------------APAAAELSTSGGGRSGGGSGSRSSLTTL 1470
EdaRWP/KAG2495229   ADKLLMSIDVLQPSLSARP-------------------------------SADKGPNGQQPGSAAAMRT- 1470
Yu.g2625.t1         ADKFLMSIDGGLPTLPSLPSLAFLHGHGQPHGHGYCNGHGHIPLPSMAPGTAPEAEGLTPGPGAAAGAA- 1470
Eu.g6369.t1         ADKFLMTIDGTQSLP--LAPTMAMPGGV--------------PLPGMPPDPANGGADADAAASASAAAA- 1470
VcRWP2              ADKFLMYIDGG-GHPGMQPGGLIGLASG------------------FPLGPIPSGTAGDAAAAAAAATA- 1470
                    ::  :

CrRWP11             ---------GYRRHQQGMEAHAALERVAEEQPDVDAEGTEEPEEEWGQGRR-------PRRGAASRAVTA 1540
CiRWP11             --------PAVGRRHVGAEAQAALDRV---VEEPQAEGAEDPAGEEGAQQGPGPKGRRQRRGPASRSVAA 1540
CsRWP11             --------PAVGRRHVGAEAQAALDRV---VEEPQAEGAEDPAGEEGAQQGPGPKGRRQRRGPASRSVAA 1540
CrRWP4              ------GRQQPQPLQSQLVQQMQQQSAGVSSAPPGSGGLAAACAAGSNTDP-------RHCLPSPAAVVA 1540
CiRWP4              NRVLMWDGAGNGQQQQQQLMHQHQQAAGASSAPPSAGSGLAAACAVSNTDP-------RHYQPTPADVVA 1540
CsRWP4              NRVLMWDGAGNGQQQQQQLMHQHQQAAGASSAPPSAGSGLAAACAVSNTDP-------RHYQPTPADVVA 1540
EdaRWP/KAG2495229   ----------------------TDNQIDSRAPPGLTSGA-------------------DGRIPSPAEVLA 1540
Yu.g2625.t1         -------SPGPDQHHPGTHGSCHSSPDGAAGAGGEPGGGPRTTDNGLPLMPQMPMGAGGHSPRAQQQQLK 1540
Eu.g6369.t1         ------AAATPQQRHAAQLAGCSNSHGSAAADSAVQNGASQPQRRQQQQQQQQQQQQQQQQEQQQLQDVG 1540
VcRWP2              --------GGPQR---------QLPAAGLHSSGAGGAGGSASAAAVDVFQN-------PHMPQAAVDGTC 1540
                                                         .

CrRWP11             RSGAPREVLQEGEDEGEDDGGTVAAAPWAA------RAAGSSRGVPSARRQLVPASPMSSPM-------M 1610
CiRWP11             HLAAA-TARAAGHDAGDVAVAAAAPAAPPEPTPAARRGNAHQSSGRQQQQLLLVQSPARSPM-------M 1610
CsRWP11             HLAAA-TARAAGHDAGDVAVAAAAPAAPPEPTPAARRGNAHQSSGRQQQQLLLVQSPARSPM-------M 1610
CrRWP4              AAAVAAAQQQQHQLSSPPLTG-----GQPQHQQQLLHGSRVVQAQAGSYESIMGGGLSAVAL-----MDL 1610
CiRWP4              AAAAA---AAAQHVASPPLPGRQPPPPHQQQQQQQQCGPAQRSAAVGSGPGGVGGGGAAVGLSAVAMTDL 1610
CsRWP4              AAAAA---AAAQHVASPPLPGRQPPPPHQQQQQQQQCGPAQRSAAVGSGPGGVGGGGAAVGLSAVAMTDL 1610
EdaRWP/KAG2495229   AVAAG-----------------------------------------------NVNDLRMAV-------DF 1610
Yu.g2625.t1         GGAAATAGAAGGAIAS--------------------AAQAAGCGGTGSGKGLHAGDPRMNMF------AL 1610
Eu.g6369.t1         NGDSA-------------------------------------HNGGGGGKGMMPYDMD------------ 1610
VcRWP2              GGDAT-------------------------------PGGGGGGGGGGGGNCLMGQDPRMIVY------DM 1610
                                                                           .

CrRWP11             TSPMRSPTAPARTFVEGPTAAAAVAA-AGAIMAPQCRAVAGVLGGGTTSIALGGNGH------------- 1680
CiRWP11             ISPIRSPAAPSSRAFAEGLAASAAMAAGGAIIAPPSRAMAGVLGGGTTSVASEDEDEGLDRMRRHQHQHQ 1680
CsRWP11             ISPIRSPAAPSSRAFAEGLAASAAMAAGGAIIAPPSRAMAGVLGGGTTSVASEDEDEGLDRMRRHQHQHQ 1680
CrRWP4              EAMSRHQLSAPAILTRLPSGSGCAAG-------------VKQFYQQQQQQQQQQQQQQQQQQPHQPHQQQ 1680
CiRWP4              EAISRHQLSAPTVFMTHPISSDDVKHFYQQQQPPIAQGGLGYLPTQHQQQQAQHMQQLHQQATFHQHQHQ 1680
CsRWP4              EAISRHQLSAPTVFMTHPISSDDVKHFYQQQQPPIAQGGLGYLPTQHQQQQAQHMQQLHQQATFHQHQHQ 1680
EdaRWP/KAG2495229   ETMMRQQISAPAVMISNAVPNAHGGAAVPNSHPHPHPQPPTAFQTNPNGFISSSQHH-----PFALHNGL 1680
Yu.g2625.t1         NTVMPHNNNNNNNNNNNNNNNNNNNN--------------------NNNNNNNNNDN---------HNNQ 1680
Eu.g6369.t1         HAMMRNQGTTPSGLCGGGVAGGGAPA---------------PYGRSSDHDAGAAAGS-----------QS 1680
VcRWP2              DAMMRSQAPTA---FGGGVGVTTSAK-------------PQSFCRPSDNTLNTGNGS-----------HT 1680
                     :

CrRWP11             -RGAPPQAAPGPSSFSAPFGGGGGGSS----LSGNSQGG----PVTARALFMMDGAGGLAGGGGGAGVTF 1750
CiRWP11             QAPPPPPAVAAHQSFSEPCQHGGGGSGAGRCLSGAGRTA----PTTARALFMMDSGGAGAGGCGGAT--F 1750
CsRWP11             QAPPPPPAVAAHQSFSEPCQHGGGGSGAGRCLSGAGRTA----PTTARALFMMDSGGAGAGGCGGAT--F 1750
CrRWP4              MQQMQQQAVQQQQQQHAAFAHQQQQQP------MPQQQQ----PHPQPQQTATSSLLQDVLSTARAVLGN 1750
CiRWP4              QQPQPQQCTATSALLQDVFS-------------TARAVL----GSATTVGNVFPDGGSSLTANAHGNNNY 1750
CsRWP4              QQPQPQQCTATSALLQDVFS-------------TARAVL----GSATTVGNVFPDGGSSLTANAHGNNNY 1750
EdaRWP/KAG2495229   VSPPYNPPAATSAAVQPSFNRNSSNQPSGPGLDGGPSMAQRFDALTKTAPTPPPHNGGGPNGTCVSVTGL 1750
Yu.g2625.t1         ESGTPPLYYQSDNASNSAYGHGH----------GTGAAAAASSPYNSSFALATAQYGNGASVDLAAA--- 1750
Eu.g6369.t1         LDALPGVDAASGGA-------------------GTPSAM----PFTSSFAMAMAQYGTVSGGPGGPA--- 1750
VcRWP2              REPLPSVDG------------------------GGSPAA----PFGSSFAFAMAQYGNCPAADGGGDVGS 1750


CrRWP11             ADGLG---GDCAPSLMPPPPPFQH----GHGHGHHNLFADPR-YGAAAHDLE--AMNRQQLSAPAVLEHP 1820
CiRWP11             ADGLADMDSKAPLTTQPPLPLHQHVLGHGHGHRQHHVIGDPR-YGVSLQDLE--AMNRQQLSAPAVLVHP 1820
```

Dataset S1

```
CsRWP11             ADGLADMDSKAPLTTQPPLPLHQHVLGHGHGHRQHHVIGDPR-YGVSLQDLE--AMNRQQLSAPAVLVHP 1820
CrRWP4              ATTVGSMFPAPAAGSHSNTAFANAHVNKAMLQSLPASAGSGVPLFTAHQQQQLMQHHQQLMATQHAS--- 1820
CiRWP4              GNIIAHAMGKSVPSFSAAPSA-------GSASGTAVGGGAGVQLFTAQQQQQLMQHHHQLLVTQHTSSGH 1820
CsRWP4              GNIIAHAMGKSVPSFSAAPSA-------GSASGTAVGGGAGVQLFTAQQQQQLMQHHHQLLVTQHTSSGH 1820
EdaRWP/KAG2495229   TNGAAAINGPAAMAARPGGPMGLP----PMGSGSISAGCSPS-LADQYQHHQ--VMYRLAGQHPGIATGP 1820
Yu.g2625.t1         YDVSAKVMSDPTPSPAAAAAALQRRMTPNQTHPLPMGGAKGL-YGSPYQETS--QLFMSSGTTATAWGS- 1820
Eu.g6369.t1         -DSLARITSAPPGTCLPDEAAFLPG---HMGHDQSLAHGMP--YNLSYGLQS--GHHLAMNSAPIGWGGN 1820
VcRWP2              VDGLARITSAP-PGVLGDESAFCPRMSSAHAHGLSMAHGKGMSYSVPYSHGQ--GHPLAMNSAPAIWTGL 1820
                        .              .                               .

CrRWP11             GAAM------LCAGAGDATATSTSGVSGGGCGGGVG-GGSVVSSAADPSGGLFAGPAAMLASLTAAAAA- 1890
CiRWP11             GAVA-------SSAAGAAGANTTSGFSAGGGGGVSS---------ADLTGGLFAGPAAMLASLTAAAAAA 1890
CsRWP11             GAVA-------SSAAGAAGANTTSGFSAGGGGGVSS---------ADLTGGLFAGPAAMLASLTAAAAAA 1890
CrRWP4              ------------SNSAGTTPTGGAAHGVPGSVPGGG--SAAIMQPGPVVSAPSPWAPQALVQLPPAAAPP 1890
CiRWP4              GAGT------TTPPAGGPQLGMGSSFTSAGGPPGSGSSLSAVMQPPGAVSAPSPWMPQALVQLSASPAP- 1890
CsRWP4              GAGT------TTPPAGGPQLGMGSSFTSAGGPPGSGSSLSAVMQPPGAVSAPSPWMPQALVQLSASPAP- 1890
EdaRWP/KAG2495229   AASL--------LGSSLPTAHGLSDWAGGGGGGGSA------------SGDSISAPLVMLPERPRSMQQQ 1890
Yu.g2625.t1         ----------LDAFGGLATITPLDQMASAPVTHATA--------GAGPGEDLMALQGDHFPGRSRSLHGH 1890
Eu.g6369.t1         YGESSGIMTGLESGGGGGGCYPGDHMNFHGNLSVHERLVMKGEIAAGGGGGGGGGGGGKMPMRCNSTQGF 1890
VcRWP2              G----------DSNLGGMDVYSMEHMG------------------------------------------- 1890
                                   .

CrRWP11             --GGGVMGMA-------------------------------------ASAPQPQVPVGLVSMGTGHSSGS 1960
CiRWP11             SSGGCCGGMG-----------------------------------ITASAPPPQARAGFGMGPGGHSSGS 1960
CsRWP11             SSGGCCGGMG-----------------------------------ITASAPPPQARAGFGMGPGGHSSGS 1960
CrRWP4              AGATGLYQLQ-----QTQQVQQSQHLQQTHTVGGPPALMQPQQQHLQQQHLHPMSLSQPLPSHNAPMMPT 1960
CiRWP4              --GTGVYQHQQQTQQQERLQRVQQMQQQAHTVAGEVALTQQLQQHLQH--LQPVSLSQPLPPIHNLAAAI 1960
CsRWP4              --GTGVYQHQQQTQQQERLQRVQQMQQQAHTVAGEVALTQQLQQHLQH--LQPVSLSQPLPPIHNLAAAI 1960
EdaRWP/KAG2495229   AING-----------------------------------------MMRSRTDASGPSVSMCPPRPFSTSS 1960
Yu.g2625.t1         CGGAGDDAVS-----------------------QQQHLLRQQHHHSPHHHHPYHGTTGNMLPHRQLSQYS 1960
Eu.g6369.t1         GGGGGGGGAG--------------------------------DEHLRHLPFLSHSGSLAAHPHSHQHQHH 1960
VcRWP2              -----------------------------------------------------------MPAPPGSAVER 1960

CrRWP11             QLYASAASVAPGGGGGLWAEQPQVQLQQHTLH---TPAPGMQQHAH-------------------SQAAP 2030
CiRWP11             QLYASAANSGGPASSGMWADPQQQQQQERLL---QPPPIMPQQA-------------------HAQQGTA 2030
CsRWP11             QLYASAANSGGPASSGMWADPQQQQQQERLL---QPPPIMPQQA-------------------HAQQGTA 2030
CrRWP4              ------SQAPPPHQLPLPQHQQHLLSQQQLQSGGIQRSISMGHYRMGPETVELVDAQR--LQKQAQQQQQ 2030
CiRWP4              MGNPHAPQQQAPHQ--QHQHQQHLQPQQPAAPGVIQRSISMGNYHMGPASVELVDA------HRLQQVQQ 2030
CsRWP4              MGNPHAPQQQAPHQ--QHQHQQHLQPQQPAAPGVIQRSISMGNYHMGPASVELVDA------HRLQQVQQ 2030
EdaRWP/KAG2495229   HGMMSMPQHAPHSPLGMIPTSQQAQHGVQFNHGAMQQPHGHSMHAM-----------------HLSGGSA 2030
Yu.g2625.t1         LQQQQQLQHQQQHQLQHYPQSPSQQALQVGQVSGSRRGRQEADLVNSCEALEVISKMRDASVQFAIAPAS 2030
Eu.g6369.t1         QHHQHHQQAHPYLRQNSQPGPLQQQQQQQQQMAAQHPHPQHHHQQ------------------NLQQHHQ 2030
VcRWP2              QQQVRQLQQQQQLQLEQLQQEQQQQQQQQLQQ---------------------------------QQLQQ 2030
                           .

CrRWP11             DVLSLQQLHSRDKASPLRSA------SFSRGSVGSSTR-------SPRRAHLNAAIQAVVKAASASPSPS 2100
CiRWP11             ADMSLQA--HRDKLSPLCST-----PSFNRGSVGSSTR-------SPRRAHLNAAIQAVVKAANASPSPS 2100
CsRWP11             ADMSLQA--HRDKLSPLCST-----PSFNRGSVGSSTR-------SPRRAHLNAAIQAVVKAANASPSPS 2100
CrRWP4              QVRSLQQPSHAQQQQQLHLH------HFQQQLPAPMMI-------SPVGPPISTMIQNHQQQQTLRPPSV 2100
CiRWP4              HHYQQQQLARSLQPPPPHAHLLQNLHHLQQMPPAPMII-------SPVGPPISTMMHQVQQQQQKTPQQQ 2100
CsRWP4              HHYQQQQLARSLQPPPPHAHLLQNLHHLQQMPPAPMII-------SPVGPPISTMMHQVQQQQQKTPQQQ 2100
EdaRWP/KAG2495229   HLVPMQQ--------PLTAN-----HHMQQGN-GHVTH-------YHVG---QGQVRVHIGAAGLHPGS- 2100
Yu.g2625.t1         HVDQQQQQHLVSQIFPDHLQ-----QQMQQQQQQQLQQ---------QQQQLQQQQQQVLQQQQQQQQQH 2100
Eu.g6369.t1         HQHQHPQQHQHHHPHPHQPQ-----SHFTQSPPPSLSTLAAQGLSASIGPTIEQQQQLQHQLQQQQQLFA 2100
VcRWP2              HQHQQQ-------------------QLFAQLFPGAQQA-------------------------------- 2100
                                               : :

CrRWP11             PLARGSD-----------------GRVIRRGAGAAAAAGFAGSMESPLAGS--AAAAAGGNSRAGSVAPQ 2170
CiRWP11             PLARDS------------------SRLRRRGGGGGGGGGSSGFGGGMDSPL------AGGSSRAGSVAPQ 2170
CsRWP11             PLARDS------------------SRLRRRGGGGGGGGGSSGFGGGMDSPL------AGGSSRAGSVAPQ 2170
CrRWP4              PPIFAS------------------DLAVGAAAHGGGAPSISTSTGNSSSAAIHIKVAAHAMPTGPLFVGS 2170
CiRWP4              VLMRHPTIPPLFNSTPPANIIMAHGVVISGTNSSTVGGGVATCNGGSAAGI-QVNVAAHELPAGPLFVGS 2170
CsRWP4              VLMRHPTIPPLFNSTPPANIIMAHGVVISGTNSSTVGGGVATCNGGSAAGI-QVNVAAHELPAGPLFVGS 2170
EdaRWP/KAG2495229   ---------------------------MSGSRRARAETDLLGSSDCLEAAI-AMAAANNGNGLNGNGHNG 2170
Yu.g2625.t1         LHMG--------------------GKSLMQRHGVSGGGSTDGTAAAMAMQAVANFAASGAGGLQPYMHGG 2170
Eu.g6369.t1         QLFPGA------------------QQVVCGVSGPGASGMVPTSNPMGQNGT----AVHPGCGAVPYMPHG 2170
VcRWP2              ---------------------------VCGGGGGSAGGSSISISPSTMMSP------TGNSPVMHFMPTS 2170

CrRWP11             LPP---------------------------------------LPTLPQLPQLPQLSPV------------ 2240
CiRWP11             LPP---------------------------------------------LPHFPQAAGG------------ 2240
CsRWP11             LPP---------------------------------------------LPHFPQAAGG------------ 2240
CrRWP4              YGPTQQQQQQQQQNLMRALEEARRSAGQQQPAGSSGGHGGGAGDNGVPCRGFARTSSGSTAGGGSAHHHQ 2240
CiRWP4              HAPSQQQQQQH---LLRALEEARRSAGQ---PGSSSGHGCATDGGATAAFGFARTSSGSIAAG------- 2240
CsRWP4              HAPSQQQQQQH---LLRALEEARRSAGQ---PGSSSGHGCATDGGATAAFGFARTSSGSIAAG------- 2240
```

Dataset S1

```
EdaRWP/KAG2495229   LPP--------------------------------------LAGGEPGLSLARISTN------------ 2240
Yu.g2625.t1         AIT-----------------------------QPEDSGSVNGGVAVGASGLSLSRISTG------------ 2240
Eu.g6369.t1         YSP-----------------------------SRPDDSCGVGLGLGPKGAPPTRILTG------------ 2240
VcRWP2              FPP--------------------------------QRDDSGCLSIGHKAHMLPRISTA------------ 2240
                      .                                                 .:

CrRWP11             -------------------------PQYAPHSDLGDGAAHHGLTAEMGLAAYDGVTAAGVAD-------- 2310
CiRWP11             ------------------------------EGAKLPGGELGGPGDDSAAGAAEERMPAKVAAIAAVAAAA 2310
CsRWP11             ------------------------------EGAKLPGGELGGPGDDSAAGAAEERMPAKVAAIAAVAAAA 2310
CrRWP4              QHQQQHQQQQPYQQQQQPALADGCNSPPAGPASRNPSGRSLLIRHTSGLARTTSAAATAAAG-------- 2310
CiRWP4              ------THPQSHNQQQHLAIADGCNSSSAGPASRNPSGRSL-LRHHSGLARTTSAAGAAAAAATAAAACG 2310
CsRWP4              ------THPQSHNQQQHLAIADGCNSSSAGPASRNPSGRSL-LRHHSGLARTTSAAGAAAAAATAAAACG 2310
EdaRWP/KAG2495229   --------------------------PSAGAAA--PPGHSHMPLTSFGSEPVGMNAAAN----------- 2310
Yu.g2625.t1         ------GQPQQQPQVQLQPQQHPQVQLPASPGV-LPRGLRPGSSNSANARAAGGSLSQG----------- 2310
Eu.g6369.t1         ----------------------AGQPPPASPSA--AGALRNGRGAAIGPGRAGGGAGAGGAMGSAGQAQS 2310
VcRWP2              -----------------VGLPPHFQPPPSPPSV--VRNMRGGCAAANSVRAGGGAAAAAAAG-------- 2310
                                                   .

CrRWP11             GAAAAAEEEAKEASTLAAAIAAATTGGGSVDGFTRKK--------------------------------- 2380
CiRWP11             AAMAATPPVFVAPSGALAMPVGTEAGRSE-GVMRDIAD-------DGDGQETAAIAAATMSSA------- 2380
CsRWP11             AAMAATPPVFVAPSGALAMPVGTEAGRSE-GVMRDIAD-------DGDGQETAAIAAATMSSA------- 2380
CrRWP4              GASGITSPRSPAAAAA-------AGGGPGTPRSRSFRQQQSQQQRSDVPPPSSLLGSGTFLAGDMGIKLE 2380
CiRWP4              SPGGITSPRSPAVVGGGGGGGGGGGGAPGTPRSRSFRQQQQQHRGDAPPPSSLLGSNTFLADGRVAIKSE 2380
CsRWP4              SPGGITSPRSPAVVGGGGGGGGGGGGAPGTPRSRSFRQQQQQHRGDAPPPSSLLGSNTFLADGRVAIKSE 2380
EdaRWP/KAG2495229   ----RQLQRSISSSNSRGSAGRSPRG-PATPRSRSRAA-------AGASLPSAAAVAAVMDEI------- 2380
Yu.g2625.t1         QSQGR-SPRASAPSTPRGSRAG-ATGTSRGGRVKSNASSLSLASPAPPLPSALAVAAALEEI------- 2380
Eu.g6369.t1         QTPGR-SPRLSAPGTPRSRPVPAAAGTPR-GRMKAGSTPLAAAGSPGASLPSSLAVRAALEEINAG---- 2380
VcRWP2              QAPAR-SPRSSAPGTPRSPRAP-ATGTPR-SRTKAGCT--------PLSAPSPGAVKAQLDEM-MRVDPV 2380
                               :             * .     :

CrRWP11             ----------GGAKAA------GGGRAGRGAKRGGAGSADIGSG--------SDDMDAGSAP-------- 2450
CiRWP11             ----------GGCSKKADSGKAGGGRGGRGAKRSTGGGADACSG-----SGSDDGMHEGPTA-------- 2450
CsRWP11             ----------GGCSKKADSGKAGGGRGGRGAKRSTGGGADACSG-----SGSDDGMHEGPTA-------- 2450
CrRWP4              DGS-TAPVGYLGAAAAAV---------PTGAAPTAALAAALAAGVMRRGGGSDDGGAGAPAAASGDSGEG 2450
CiRWP4              DGSGLAPLGYMGASAAAAAASMAGAGVATGTAPTAALAAAIAAGVVGRGAS-DDGAAGESGD-------- 2450
CsRWP4              DGSGLAPLGYMGASAAAAAASMAGAGVATGTAPTAALAAAIAAGVVGRGAS-DDGAAGESGD-------- 2450
EdaRWP/KAG2495229   ----------GGAPSAAAATSSGDGGAVSGAAPTVAVAAAVAAGVV---PA-PDGTASADGT-------- 2450
Yu.g2625.t1         ---------------------IRSDPTAPAAAAAVAAAAAVAVGLA---AG-HDGAASADGP-------- 2450
Eu.g6369.t1         ----------GGGGGASAESLAAAAAVAGSTAPEAALAAATAAGLI---VA-EDGSGGADSPFAGGGGGG 2450
VcRWP2              AAAAAAAGGGSGMSESLAAAAAAAAVAAVAAAPEAALAAAAAAGLI---VA-EDAGAAVDSP-------- 2450
                                                 .:    . .*    *         *

CrRWP11             -----SGGG---AAPTAERPSGVGRLG-------AAAPSGRGSQMDSGDEES-------GGGGGGDEEGD 2520
CiRWP11             ----GPWCG---GAMSKEEPAEEGQVG----AALALALGCIGGRTDSGNDNE---TGDDGGGGGGDGEGD 2520
CsRWP11             ----GPWCG---GAMSKEEPAEEGQVG----AALALALGCIGGRTDSGNDNE---TGDDGGGGGGDGEGD 2520
CrRWP4              SGADADAAG---GTVRRSRREPRGG------VKTSRSGGAGGRRRRAGDDDDAPYETDDAEEEESSDGSD 2520
CiRWP4              ASGGNAEAAAAAGGVRRSRREQRGADV----KSARGSGAGNGRRRRNGEDDDESYETDDAEEEEESSEGSD 2520
CsRWP4              ASGGNAEAAAAAGGVRRSRREQRGADV----KSARGSGAGNGRRRRNGEDDDESYETDDAEEEEESSEGSD 2520
EdaRWP/KAG2495229   SAGADDSAG---GGLRRSTRGTRANRA-------ASSRGGRG-RGRSREDDDEDYAAEDDDDASSEDEDD 2520
Yu.g2625.t1         --GPSSAGG---GSLRRSRRETRGSKV----SSLAAARGSRHSNRSRVDEDDDDYPSDADEEGLSSDDGE 2520
Eu.g6369.t1         GGGGGGGGG---GGQRRSRREARNGKGANTAAAAAAARGSRGGRSRGTDDDDEDYATDVDDDALTSEEGD 2520
VcRWP2              ----GASGG---GNLRRSRREARGARG----STLGTARTGRGGRSRGTDDDDEDYATDADEDALSSDDGD 2520
                            .   .    .                  :      .    :::.             .  .:

CrRWP11             ------------GGGV-----------SGGG-------EGV----SVVNLRKVTQDGAPRQLTKQSLKDV 2590
CiRWP11             ---DEDGGGSFSAGGF-----------SAGG-------GGVIEGPNVVNLRKVTLDGMPRQLTKQSLKDV 2590
CsRWP11             ---DEDGGGSFSAGGF-----------SAGG-------GGVIEGPNVVNLRKVTLDGMPRQLTKQSLKDV 2590
CrRWP4              SDLDDDFLGGGG-GGRRSRKL------GRDGGDDDVDGGYYVDDNSVLVVRAV-KDGQRREITKHALRKV 2590
CiRWP4              SDLEDDFLDGGCGGGRRARRSGGRNGDGGGGGSDDVDEGYYVDDDSVLVVRAV-KDGQRREITKQALRKV 2590
CsRWP4              SDLEDDFLDGGCGGGRRARRSGGRNGDGGGGGSDDVDEGYYVDDDSVLVVRAV-KDGQRREITKQALRKV 2590
EdaRWP/KAG2495229   ---SSGGRRGLGPGGE-----------DEEE-------EGFVDEDSVLVVRPV-KDGPRREITKMALRKV 2590
Yu.g2625.t1         ---EGSGRSGGGVGGD-----------GEGG------VDGFVDENSVLVVRAV-RDGVRREITKNALRNV 2590
Eu.g6369.t1         ---EEGGAAGGSGGCRS----------GVGG-------EGYVDEDSVLVVRAV-RDGVRREITKSALRKV 2590
VcRWP2              ---DEGGGGGGGAGRG-----------GGGGGGGTTSGEGYVDEDSVLVVRAV-RDGVRREITKSALRKV 2590
                                 *             .                 .*: :* *  **  *::** :*:.*

CrRWP11             YHLPINEAAAALNIGVTVLKKYCRKFSIPRWPYRKLNSVNKLMETFERYKRDALLGGNVTGGEECEVVLQ 2660
CiRWP11             YHLPINEAAAALNIGVTVLKKYCRKFSIPRWPYRKLNSVQKLSETFERYKRDALLNGNFTGGEECEVVLQ 2660
CsRWP11             YHLPINEAAAALNIGVTVLKKYCRKFSIPRWPYRKLNSVQKLSETFERYKRDALLNGNFTGGEECEVVLQ 2660
CrRWP4              YHLPINEAAAALNIGVTVLKKYCRKFRVPRWPYRKLQSMGKLIESFQRYKKGALGAGNTDEAERCEAVIQ 2660
CiRWP4              YHLPINEAAAALNIGVTVLKKYCRKFRVPRWPYRKLQSMAKLIESFQRYKKGALASGNPDEAERCEAVIQ 2660
CsRWP4              YHLPINEAAAALNIGVTVLKKYCRKFRVPRWPYRKLQSMAKLIESFQRYKKGALASGNPDEAERCEAVIQ 2660
EdaRWP/KAG2495229   YHLPINEAAAALNIGVTVLKKYCRKFRVPRWPYRKLQSMGKLIESFQKYKRAAIANADLEAAERCEVVIQ 2660
Yu.g2625.t1         YHLPINEAAAALNIGVTVLKKYCRKFRVPRWPYRKLQSLAKLIDSFQRYKRMAISNADLANAEHCEATIN 2660
Eu.g6369.t1         YHLPINEAAAALNIGVTVLKKYCRKFRVPRWPYRKLQSMAKLIESFERYKKNAFSNGDMEGAADCQDVIA 2660
VcRWP2              YHLPINEAAAALNIGVTVLKKYCRKFRVPRWPYRKLQSMAKLIESFERYKKNAFSNGDIEGAADCQDVID 2660
```

```
                    ************************** :********:*: ** ::*::**: *:  .:   .  *: .:

CrRWP11             SLGKMKVELYEDPDKDIDERIKKLRQANFKVEYRARQDTTGKQQQPQQQQHQDEFQQLLQQQLQQHLMQQ 2730
CiRWP11             SLGKMKMELYEDPDKDIDERIKKLRQANFKVEYRARQDTSGKQQQQQQQQQDDQHLQ-QQLQLQQHLMQQ 2730
CsRWP11             SLGKMKMELYEDPDKDIDERIKKLRQANFKVEYRARQDTSGKQQQQQQQQQDDQHLQ-QQLQLQQHLMQQ 2730
CrRWP4              SLYRFREEVYDNPDKDIDESVKKLRQANFKIEYRQRQGQT------------------------------ 2730
CiRWP4              SLYRFREEVYDNPDKDIDESVKKLRQANFKIEYRQRQGQP------------------------------ 2730
CsRWP4              SLYRFREEVYDNPDKDIDESVKKLRQANFKIEYRQRQGQP------------------------------ 2730
EdaRWP/KAG2495229   SLHRFREEVYDNPDKDIDEGVKKLRQANFKIEYRQRQGQS------------------------------ 2730
Yu.g2625.t1         SLMKFREELYDNPDKDIDESVKKLRQANFKIEYRQRQGQT------------------------------ 2730
Eu.g6369.t1         RLLKFREEVYDNPDKDIDESVKKLRQANFKIEYRQRQGQT------------------------------ 2730
VcRWP2              RLLKFREEVYDNPDKDIDESVKKLRQANFKIEYRQRQGQT------------------------------ 2730
                     * ::: *:*::******* :*********:*** **. .

CrRWP11             QQQQMFLPQQQQQQQQQQELPGAQGQQMLMP--VGDGFGM----HGGMPQGAAGMLPAMPPPFLLPPFNP 2800
CiRWP11             QQVFM-------QQQQLHGLFGGMSCAMGLPALASEGFAMAAGGSSGLLVGPGGILAPMPPPYLLPPLDP 2800
CsRWP11             QQVFM-------QQQQLHGLFGGMSCAMGLPALASEGFAMAAGGSSGLLVGPGGILAPMPPPYLLPPLDP 2800
CrRWP4              --------------------QGAPSTAAAAADI-----------LIEDGIGMEGV-----------AME- 2800
CiRWP4              --------------------LGAPSNAAAAADI-----------LLEDGIDMDGT-----------AMDM 2800
CsRWP4              --------------------LGAPSNAAAAADI-----------LLEDGIDMDGT-----------AMDM 2800
EdaRWP/KAG2495229   ---------------------GPGAAATAGSDILTGASGPMG--SADLGRELAGL-TVSPDGTILPALDG 2800
Yu.g2625.t1         --------------------QGPPSGSLAEPSP-----------TAADQHAAGGMVVVGSSNDLRRDFET 2800
Eu.g6369.t1         --------------------QGAPPTGIARTSLSGGAAVE----VEGAPVAVAGV---ASSNDLRRDFET 2800
VcRWP2              --------------------QGAPPSTLTRSSLNGE--------TPGAATGVGGA---PSSHDLRRDFET 2800
                                         *        .                      *             ::

CrRWP11             -AALPSQFL--PEGSAQGFGGGCPAA-NPTNF 2832
CiRWP11             TAALPPQFLPPPEGTPGVIGGSSAGSVMPTHF 2832
CsRWP11             TAALPPQFLPPPEGTPGVIGGSSAGSVMPTHF 2832
CrRWP4              -------------AGRGGAEGGSSSTSEAVAA 2832
CiRWP4              QAG----------GGGGGAEGGSSSTSETVAA 2832
CsRWP4              QAG----------GGGGGAEGGSSSTSETVAA 2832
EdaRWP/KAG2495229   AGG--------AAAAGGGSGGGAPGVL----- 2832
Yu.g2625.t1         LTV-----------NGASEGGQELSSL----- 2832
Eu.g6369.t1         LSV----------NSAAVEAAKEMTPT----- 2832
VcRWP2              LSV----------NAASAEVTEPPPA------ 2832
```
