## Supplementary material for "A conserved RWP-RK transcription factor VSR1 controls gametic differentiation in volvocine algae": Dataset S2

| name | gene | template | Forward primer name | Forward primer sequence | Reverse primer name | Reverse primer sequence | PCR condition | restriction fragment (backbone/insert) | assembly |
|---|---|---|---|---|---|---|---|---|---|
| pGAD-CrMID | CrMID | CC-125 mt- cDNA | P174-MID-EcoF | gaattcATGGCCTGTTTCTTAGCCAGGTTCCAGT | P175-MID-XhoR | aggatgctcgagCATGTGTTTCTTGACGCTGGCGACCTTTC | KOD FX Neo (TOYOBO), 2 step-cycle protocol, with 30 sec extension, 28 cycles | pGADT7 EcoRI/XhoI digest | DNA ligation high (TOYOBO) |
| pGAD-CrRWP11 | CrRWP11 | CC-125 mt- cDNA | P178-RWP11-EcoF | gaattcATGGACCTCGACGTGCCTGACCTCCTC | P179-RWP11-XhoR | aggatgctcgagGAAGTTGGTTGGATTCGCGGCAGGGCAG | KOD FX Neo (TOYOBO), 2 step-cycle protocol, with 5 min extension, 28 cycles | pGADT7 EcoRI/XhoI digest | DNA ligation high (TOYOBO) |
| pGAD-CrRWP11^(431-691) | CrRWP11^(431-691) | pGAD-CrRWP11 (this study) | P508-R11-28NEB-F | GCTCATATGGCCATGGAGGCCAGTGCGGGGCGCATCCTGGCAACC | P563-No38-R11-R | CGATTCATCTGCAGCtcgagGTCAGGGTTGCAGTCGTCTGCC | KOD FX Neo (TOYOBO), 2 step-cycle protocol, with 30 sec extension, 25 cycles | pGADT7 EcoRI/XhoI digest | NEBuilder (NEB) |
| pGAD-CrRWP11^(702-2123) | CrRWP11^(702-2123) | pGAD-CrRWP11 | P509-R11-38NEB-F | GCTCATATGGCCATGGAGGCCAGTGGCGAGCTGGAGTGCACGCTC | P510-R11-Nru30Rev | GCGGTCAACGGCAGGTCTGGGAAATCATCG | KOD FX Neo (TOYOBO), 2 step-cycle protocol, with 30 sec extension, 25 cycles | pGAD-CrRWP11 EcoRI/NruI digest | NEBuilder (NEB) |
| pGAD-VcMID | VcMID | pL23:VcMIDcDNA-6xFLAG (Geng et al 2018) | VcMID_Y2H_F | | VcMID_Y2H_R1 | | | pGADT7 BamHI/XhoI digest | DNA ligase |
| pGAD-VcRWP2 | VcRWP2 | pGADT7 | P429-VcRWP2-Ter-F | GAGCCGCCGCCGGCTctcgaGCTGCAGATGAATCGTAGAT | P428-VcRWP2-AD-R | CGCAAGGTCCTCCATgaattCACTGGCCTCCATGGCCATA | KOD FX Neo (TOYOBO), 2 step-cycle protocol, with 4 min extension, 25 cycles | pNitA-SF-VcVSR1cDNA MluI/XbaI digest | NEBuilder (NEB) |
| pGAD-VcRWP2^(485-694) | VcRWP2^(485-694) | pGAD-VcRWP2 | P571-VC2X8-F | CCATGGAGGCCAGTGAGTTCGTCAATTTGGCTTGGAACGCAAACT | P575-VC2Q9-R | CGATTCATCTGCAGCTCGAGTTATTGCTGAAACTGAGGCGCTTGC | KOD FX Neo (TOYOBO), 2 step-cycle protocol, with 30 sec extension | pGADT7 EcoRI/XhoI digest | NEBuilder (NEB) |
| pGAD-VcRWP2^(563-996) | VcRWP2^(563-996) | pGAD-VcRWP2 | P572-VC2X9-F | CCATGGAGGCCAGTGAAGGTGACAAGGGGACAGAGCTCCTAACCG | P577-VC2X9-R | CGATTCATCTGCAGCTCGAGTTACATGATCATGTCCAGGCTCCCT | KOD FX Neo (TOYOBO), 2 step-cycle protocol, with 1 min extension, 25 cycles | pGADT7 EcoRI/XhoI digest | NEBuilder (NEB) |
| pGBK-CrMID | CrMID | CC-125 mt- cDNA | P174-MID-EcoF | gaattcATGGCCTGTTTCTTAGCCAGGTTCCAGT | P175-MID-XhoR | aggatgctcgagCATGTGTTTCTTGACGCTGGCGACCTTTC | KOD FX Neo (TOYOBO), 2 step-cycle protocol, with 30 sec extension, 28 cycles | pGBKT7 EcoRI/SalI digest | DNA ligation high (TOYOBO) |
| pGBK-CrMID-N | CrMID-N | pGAD-CrMID | P547-BK-CrMID-F | TGGCCATGGAGGCCGaattcATGGC | P546-CrMIDNXho-R | aTGCGGCCGCTGCAGCtcgagCTTCTTGGCCGGCTGTGCGGGCAA | KOD FX Neo (TOYOBO), 2 step-cycle protocol, with 30 sec extension, 25 cycles | pGBKT7 EcoRI/SalI digest | NEBuilder (NEB) |
| pGBK-CrMID-C | CrMID-C | pGAD-CrMID | P548-BK-MIDC-F | TGGCCATGGAGGCCGaattcGCTGACTTGACACTTCATGACATC | P549-CrMID-BK-R | TGCGGCCGCTGCAGctcgagCATGT | KOD FX Neo (TOYOBO), 2 step-cycle protocol, with 30 sec extension, 25 cycles | pGBKT7 EcoRI/SalI digest | NEBuilder (NEB) |
| pGBK-CrRWP11^(431-691) | CrRWP11^(431-691) | pGAD-CrRWP11 | P592-R11-IA1-BK-F | TGGCCATGGAGGCCGCGGGGCGCATCCTGGCAACCGCCGCCGTGC | P590-R11IA1-BK-R | TGCGGCCGCTGCAGGTCAGGGTTGCAGTCGTCTGCCAGTGCCGCC | KOD FX Neo (TOYOBO), 2 step-cycle protocol, with 30 sec extension, 25 cycles | pGBKT7 EcoRI/SalI digest | NEBuilder (NEB) |
| pGBK-VcMID | VcMID | pL23:VcMIDcDNA-6xFLAG (Geng et al 2018) | VcMID_Y2H_F | | VcMID_Y2H_R2 | | | pGBKT7 BamHI/PstI digest | DNA ligase |
| pGBK-VcRWP2^(485-562) | VcRWP2^(485-562) | pGAD-VcRWP2 | P591-VC2X8-BK-F | TGGCCATGGAGGCCGAGTTCGTCAATTTGGCTTGGAACGCAAACT | P588-VcRWP2-X8-BK-R | TGCGGCCGCTGCAGGTAAATGGGCTTTATCCCTTTTGACTGGCAA | KOD FX Neo (TOYOBO), 2 step-cycle protocol, with 30 sec extension, 25 cycles | pGBKT7 EcoRI/SalI digest | NEBuilder (NEB) |
