## Supplementary material for "A conserved RWP-RK transcription factor VSR1 controls gametic differentiation in volvocine algae": SI

**Supporting Information for**
A conserved RWP-RK transcription factor VSR1 controls gametic differentiation in volvocine algae

Sa Geng [a,1], Takashi Hamaji [a,b,c,1], Patrick J. Ferris [a], Minglu Gao [a], Yoshiki Nishimura [b], James Umen [a,*]
a) Donald Danforth Plant Science Center, St Louis, MO, USA
b) Department of Botany, Graduate School of Science, Kyoto University, Kyoto, Kyoto, Japan
c) Research and Development Initiative, Chuo University, Bunkyo-ku, Tokyo, Japan
1) These authors contributed equally.
* corresponding author: James Umen

This PDF file includes:

Supporting text
Tables S1 to S6
Figures S1 to S12
Legends for Datasets S1 to S2
SI References

Other supporting materials for this manuscript include the following:

Datasets S1 and S2

### Supporting Information Text 1

**Cloning *VcVSR1* genomic DNA, *VcVSR1* cDNA and *CrVSR1* genomic DNA.** For Volvox VSR1, PCR was used to amplify genomic fragments and cDNAs of *VSR1* from wild type female strain EVE. Genomic DNA was isolated as described previously (1), and cDNAs were prepared as described above. The high GC content and low complexity of *VSR1* necessitated using a modified 10X dNTP mix for PCR amplifications containing 0.5 µM dGTP, 1.5 µM 7-deaza-2'deoxy dGTP, and 2 µM each dATP, dCTP, dTTP. Phusion enzyme (NEB) was used for amplifications according to the manufacturer's instructions. PCR products were amplified using 32 cycles of 98°C for 20 s, 60-68°C for 20s, and 72°C for 30s-120s.

Two separate fragments that together covered the whole *VcVSR1* gene were amplified from female genomic DNA (isolated as described above) using primer sets VcVSR1_F1-2(907)/VcVSR1-R23(904) and VcVSR1-F23(903)/ VcVSR1_R6-2(908). The two PCR fragments were assembled into Chlamydomonas vector AR:Luciferase:RBCS2 (2, 3) that was digested with XhoI/BamHI to remove the luciferase fragment and using Gibson Assembly® Master Mix (NEB, E2611) to create vector pAR-VcVSR1 containing a full length genomic clone of VcVSR1 from predicted ATG to stop codon. pAR-VcVSR1 was further modified by inserting FLAG(F) and StrepII(S) epitope tags at the 5'end and replacing the Chlamydomonas AR promoter with the Volvox NitA promoter. To do so, the vector pVcCas9-1 (4) was used as a template with amplification primer set NitA3'UTR_F4(sg44)/ NitA 5'UTR_R4(sg45) to generate a NitA promoter fragment. Epitope tags FLAG(F) and StrepII(S) block fragment were amplified with primer set gBlock_TAP_F1(sg46)/ gBlock_TAP_R2 (sg49) from the following synthetic fragment 5'-GGCTCCTGGAGCCACCCCCAGTTCGAGAAGGGCGGCGGCAGCGGCGGTGGTTCGGGCGGCGGCAGCTGGTCGCATCCCCAGTTCGAGAAGGGCAGCTACCCCTACGACGTGCCCGATTACGCGCACCACCACCACCACCATGACATCCCCACCACTGCGAGCGAGAACCTGTACTTCCAGGGCGAGCTGGACTACAAGGACCACGACGGCGACTACAAGGACCATGACATTGACTACAAGGACGACGACGACAAGGACATCCCCACCACCGCTTCG-3'. Two VcVSR1 gene fragments were amplified from the vector pAR-VcVSR1 with the primer sets VcVSR1-F41(sg48/))/VcVSR1-R23(904) and VcVSR1-F23(903)/VcVSR1-R35(sg43). Then the two VcVSR1 genomic fragments, tandem epitope tag fragment and NitA promoter fragment were assembled with the backbone of pVcCas9-1 digested by SpeI/BamHI following the recipe of Gibson assembly (NEB) to get vector pNitA-SF-VcVSR1. To generate the construct pSF-VSR1, the pNitA-SF-VcVSR1 vector backbone was amplified with the primer set HRcas9-vector_R(sg125)/NitA3'UTR_F4(44). The VcVSR1 native 5'UTR and promoter were amplified from genomic DNA by primer set VcVSR1_5'UTR_F(122)/ VcVSR1_5'UTR_R(123). The tandem epitope tag FLAG(F) and Strep II(S) was amplified with primer set gBlock_TAP_F4 (sg124)/ gBlock_TAP_R2 (sg49) and the vector pNitA-SF-VcVSR1 was cut with MluI and PacI to recover a 9.2kb VcVSR1 genomic fragment. Then the four fragments were assembled following the recipe of Gibson assembly (NEB) to get the vector pSF-VSR1. All the clones were verified by Sanger sequencing.

For Chlamydomonas *VSR1*, a 9.5 kb fragment containing the full-length genomic region of *CrVSR1* was obtained by digesting BAC clone 9498 using enzymes SspI/Mfe (http://www.chlamy.org/bac_details.html), ligated to vector pBlueScript II KS (5) amplified with primers pBlueCrVSR1SSP1R(782)/ pBlueCrVSR1MfeIF(783) (http://www.chlamy.org/bac_details.html) and digested with SspI/Mfe to get pKSCrVSR1 (Primers described in Table S7). Two fragments that cover the whole length of CrVSR1 genomic DNA were amplified from pKSCrVSR1 with primer set CrVSR1_XhoI_F(780)/CrVSR1-mid-R(795) and CrVSR1-mid-F(794)/CrVSR1_BamHI_R(781), and assembled using Gibson assembly (NEB) with the vector from pRPL23:Luc:RPL23 (6) digested with enzymes XhoI/BamHI to excise the luciferase gene and create pRPL23CrVSR1. To generate AphVII-RPL23CrVSR1, the AphVII expression cassette was amplified from pKS-aph7-lox (7) with primer set XbaI-Hyg-F and XbaI-Hyg-R. The amplicon was digested with XbaI and ligated to XbaI digested pRPL23CrVSR1 to generate the vector pAphVII-RPL23CrVSR1. To generate the vector pRPL23-3HA-CrVSR1, four fragments were assembled. First, vector pAphVII-RPL23CrVSR1 was digested with SpeI/NsiI to recover a ~11kb fragment with a small portion of VSR1 removed from the 5' region. Second, the 3xHA_F (sg234)/ 3xHA_R3 (sg238) primers were used to amplify the triple HA epitope tag from 3xHA-MAT3 (8). Third, Primers RPL23_SpeI-F(sg264) and RPL23_R (sg265) were used to amplify a fragment from AphVII-RPL23CrVSR1 between the SpeI site and ATG at the start site. Fourth, the primer set CrVSR1_atg_f (sg232)/ CrVSR1_NsiI_R (sg267) were used to amplify the fragment from AphVII-RPL23CrVSR1 between the ATG and the internal NsiI site as the fourth fragment. Then the four fragments were assembled following the recipe of Gibson assembly (NEB) to generate the construct pHA-CrVSR1. All the clones were confirmed by Sanger sequencing.

The plasmid pVcMID-GFP-Ollas (pVcMID-GO) was created as follows: the two oligos Ollas_VcMid_Ve, gcgaatgaattgggacctaggttgatgggcaagctttctagatagcggccgcGGATCCATTTGCGATTGCTTGCTCTTCGTA and Ollas-GFP_R, atcaacctaggtcccaattcattcgcaaagccactagtcatggtagatctggctgcCTTGTACAGCTCGTCCATGCCGT were annealed and used to create a double stranded fragment by running PCR program, 20 cycles of 98°C for 20 s, 68°C for 20s, and 72°C for 10s with Phusion enzyme (NEB) and HF buffer, and the product was used as a reverse primer to pair with primer VcMid-GFP_F to amplify GFP from pMF124cGFP (9) to create a GFP-OLLAS fusion fragment. The plasmid pVcMID-BH (2) was digested with NheI and BamHI, and the 3.5kb backbone was assembled with GFP-OLLAS fragments using Gibson assembly (NEB) to generate pVcMID-GFP-Ollas (pVcMID-GO).

Supporting Information Text 2

***V. carteri* transformation.** All transformations of Volvox were done as described previously (2) with 10 µg/ml Hygromycin B used for selection of edited transformants. The AichiM *vsr1* mutant candidates from the Hygromycin B selection medium were tested for sexual differentiation, and multiple independent candidates were found that did not develop sperm packets and were characterized further by amplifying the edited region using PCR (Table S7) and sequencing. Edited male mutant line *vsr1*$^{\#5\_4}$ (*vsr1-1*) was used for rescue experiments. Construct pSF-VSR1 was contransformed into *vsr1-1* along with pPmr3 which encodes a paromomycin resistance marker (10) with selection of rescued strains in medium containing 10 µg/ml of paromomycin. Eve *vsr1-1* mutants were generated by crossing Eve and a selected rescued male strain *vsr1::SF-VSR1*. Around 70 meiotic progeny were scored by genotyping (sex determining region, *VSR1* locus, and presence/absence of pSF-VSR1 construct) and phenotypic analysis of sexual development. Genotyping *vsr1-1* was facilitated by use of a *Hpy1*88I restriction site polymorphism created by the edited mutation (Figure S10, Table S4). Strains for Co-IP experiments with MID and VSR1 were generated by transforming either wild-type Eve or rescued female strain *vsr1::SF-VSR1*#1 with plasmid VcMID-GO and selecting for paromomycin resistance carried by the plasmid. Pseudo-male transformants were identified by sexual induction and scoring for sperm packet formation as described previously (2), and MID expression was confirmed by immunoblotting. Selected female VcMID-GO expressing and female *vsr1::SF-VSR1* VcMID-GO expressing transformants were used for Co-IP experiments.

**Supplementary Tables**

**Table S1.** Summary of the genetic segregation for crossing *rwp4 MT*$^-$ x CC-5155 *MT*$^+$

| *marker phenotype | numbers of progeny | Genotype | mating competence |
|---|---|---|---|
| Paro$^R$ | 10 | *rwp4-1 MT*$^+$ | Yes |
| Paro$^R$ | 15 | *rwp4-1 MT*$^+$ | Yes |
| Paro$^S$ | 12 | *RWP4 MT*$^+$ | Yes |
| Paro$^S$ | 11 | *RWP4 MT*$^-$ | Yes |

*paromomycin (Paro) resistance (R) or sensitivity (S)

**Table S2.** Summary of genetic segregation in Chlamydomonas cross *vsr1-1 HA-VSR1 MT*$^+$ x CC-5325 *MT*$^-$

| marker phenotype[1] | numbers of progeny[2] | genotype | mating competence |
|---|---|---|---|
| Paro$^S$+Hyg$^S$ | 4 | *VSR1 MT*$^+$ | yes |
| Paro$^S$+Hyg$^S$ | 8 | *VSR1 MT*$^-$ | |
| Paro$^R$+Hyg$^S$ | 5 | *vsr1-1 MT*$^+$ | no |
| Paro$^R$+Hyg$^S$ | 5 | *vsr1-1 MT*$^-$ | |
| Paro$^S$+Hyg$^R$ | 7 | *VSR1 HA-VSR1 MT*$^+$ | yes |
| Paro$^S$+Hyg$^R$ | 9 | *VSR1 HA-VSR1 MT*$^-$ | |
| Paro$^R$+Hyg$^R$ | 6 | *vsr1-1 HA-VSR1 MT*$^+$ | |
| Paro$^R$+Hyg$^R$ | 4 | *vsr1-1 HA-VSR1 MT*$^+$ | |

1. Indicates resistance (R) or sensitivity (S) to paromomycin (Paro) or hygromycin (Hyg).
2. Progeny from *vsr1-1::HA-VSR1 MT*$^+$ x CC-5325 *MT*$^-$.

**Table S3.** Summary of genetic segregation for *syp1* and *vsr1* mutations from rescued CLiP strain LMJ.RY0402.189640 crossed to wild type parent CC-5155.

| *marker phenotype | numbers of progeny | genotype | mating competence |
|---|---|---|---|
| Paro$^{S}$+Hyg$^{S}$ | 6 | *VSR1 SYP1 MT*$^{+}$ | Yes |
| | 6 | *VSR1 SYP1 MT*$^{-}$ | |
| Paro$^{S}$+Hyg$^{R}$ | 1 | *VSR1 HA-VSR1 SYP1 MT*$^{+}$ | |
| | 3 | *VSR1 HA-VSR1 SYP1 MT*$^{-}$ | |
| Paro$^{R}$+Hyg$^{S}$ | 2 | *vsr1-1 SYP1 MT*$^{+}$ | No |
| | 2 | *vsr1-1 SYP1 MT*$^{-}$ | |
| | 2 | *VSR1 syp1 MT*$^{+}$ | Yes |
| | 1 | *VSR1 syp1 MT*$^{-}$ | |
| | 5 | *vsr1-1 syp1 MT*$^{+}$ | No |
| | 2 | *vsr1-1 syp1 MT*$^{-}$ | |
| Paro$^{R}$+Hyg$^{R}$ | 6 | *VSR1 HA-VSR1 SYP1 MT*$^{+}$ | Yes |
| | 4 | *VSR1 HA-VSR1 SYP1 MT*$^{-}$ | |
| | 3 | *vsr1-1 HA-VSR1 syp1 MT*$^{+}$ | |
| | 3 | *vsr1-1 HA-VSR1 syp1 MT*$^{-}$ | |
| | 1 | *VSR1 HA-VSR1 syp1 MT*$^{+}$ | |
| | 2 | *VSR1 HA-VSR1 syp1 MT*$^{-}$ | |

**Table S4.** Summary genetic segregation in Volvox cross Eve (female) x *vsr1-1::SF-VSR1* (male).

| *genotype | numbers of progeny | gametogenesis/fertility |
|---|---|---|
| *VSR1 MTF* | 6 | yes |
| *vsr1-1 MTF* | 12 | no |
| *vsr1-1 SF-VSR1 MTF* | 7 | yes |
| *VSR1 SF-VSR1 MTF* | 8 | yes |
| *VSR1 MTM* | 4 | yes |
| *vsr1-1 MTM* | 11 | no |
| *vsr1-1 SF-VSR1 MTM* | 8 | yes |
| *VSR1 SF-VSR1 MTM* | 14 | yes |

**vsr1-1* is the allele designation for mutation in edited strain *vsr1*$^{\#5\_4}$ shown in Figure 3. *MTF*, female sex determining region. *MTM*, male sex determining region.

**Table S5.** Summary of RWP-RK transcription factors known or suspected to be sex-related.

| Name | Organism | Function | References |
|---|---|---|---|
| RKD1, RKD2 | Arabidopsis (Embryophyte) | Egg cell differentiation | (11–14) |
| RKD | Marchantia (Bryophyte) | Egg cell differentiation and spermatogenesis | (14–16) |
| Minus1 | *Closterium peracerosum–strigosum–littorale* (Zygnematophyceae) | Minus gamete differentiation | (17) |
| MLPa | *Ostreococcus* (Mamiellophyceae) | Unknown, but encoded in putative heteromorphic mating-type locus | (18, 19) |
| MLPb | *Micromonas* (Mamiellophyceae) | Unknown, but encoded in putative mating locus | (19) |
| RWP1 | *Ulva partita* (Ulvophyceae) | Unknown, present in male mating locus | (20) |
| UMSL057_0048 | *Ulva mutabilis* (Ulvophyceae) | Unknown, upregulated in gametogenesis | (21) |
| MID | Volvocine algae (Chlorophyceae) | *minus*/male dominant determination factor | (2, 22) |
| VSR1 | Volvocine algae (Chlorophyceae) | Gamete mating type/sex differentiation | This study |

**Table S6.** Oligonucleotides annealed to generate sgRNA vectors for targeting Volvox *RWP2*.

| Targeted Gene | Target Vector | Oligonucleotide sequences [a] | Protospacer/PAM strand |
|---|---|---|---|
| *RWP2* | RWP2-sgRNA-1 | 5’-tgacCTCCAGTACTGCTCTCGTGG-3’ | noncoding |
| | | 5’-aaacCCACGAGAGCAGTACTGGAG-3’ | |
| | RWP2-sgRNA-5 | 5’-tgacAGGTCCATTGCGCCACCTGG-3’ | noncoding |
| | | 5’-aaacCCAGGTGGCGCAATGGACCT-3’ | |

[a] lower case letters show BtgZI overhang nucleotides in sgRNA vector, and uppercase letters correspond to nucleotides in targeted region of *RWP2*.

**Table S7.** Oligonucleotides for cloning, genotyping and qRT-PCR.

| Name | Oligonucleotide Sequence (5' to 3') |
|---|---|
| Plasmid creation | |
| RWP2_F1-2 | GTTTCCATTTGCAGGATGCTCGAGATGAACACAGGCAAAGGTCTGAAAGCC |
| RWP2-R23 | CGTAAGGTTTTCGGCCAGTA |
| RWP2-F23 | TACTGGCCGAAAACCTTACG |
| RWP2_R6-2 | CTCCATTTACACGGAGCGGGGATCCCTAAGCCGGCGGCGGCTCTGTGACCTCAGCGGAAG |
| NitA3'UTR_F4 | CCGCCGGCTTAGTTAATTAAGAATTAATTCGGATCCCGATCCGACTC |
| NitA 5'UTR_R4 | TTGCCTGTGTTCATACGCGTACTAGTGAACTTCTGCAAATGGAAACGG |
| gBlock_TAP_F1 | ACGCGTATGAACACAGGCAAGGCTCCTGGAGCCACCCC |
| gBlock_TAP_R2 | GCAAGGTCCTCCATACGCGTCGAAGCGGTGGTGGGGATGT |
| RWP2-F41 | ACGCGTATGGAGGACCTTGCGGAAGTCCT |
| RWP2-R35 | TTAATTAACTAAGCCGGCGGCGGCTCTGTGACCTCAGCGGAAG |
| NitA3'UTR_F4 | CCGCCGGCTTAGTTAATTAAGAATTAATTCGGATCCCGATCCGACTC |
| HRcas9-vector_R | GTAGGCTGGTGGCTGGGCTCCCGGGAATCGAATTCCCGCGGCCGCC |
| RWP2_5'UTR_F | AGCCCAGCCACCAGCCTACTTC |
| RWP2_5'UTR_R | GAGCCTCCGGCAAGCACGC |
| gBlock_TAP_F4 | GCGTGCTTGCCGGAGGCTCATGGGCTCCTGGAGCCACCCCCA |
| gBlock_TAP_R2 | GCAAGGTCCTCCATACGCGTCGAAGCGGTGGTGGGGATGT |
| RWP2cDNA_R1 | CTGCTGCTGTTGTTGATGCTGC |
| rwp2_F34 | GCAGCATCAACAACAGCAGCAG |
| rwp2_R30 | GTTACGTACGACGGATGGTGGG |
| rwp2_F35 | CCCACCATCCGTCGTACGTAAC |
| pBlueRWP11SSP1R | TTCTTAATTCTTAATATTGCTACTTCACTGGCCGTCGTTTTACAACGTC |
| pBlueRWP11MfeIF | CTGTCTACGCACGATCAATTGcatggtcatagctgtttcctgtgtg |
| RWP11_XhoI_F | CTGCGCGCAGAGTCTCGAGATGGACCTCGACGTGCCTGACC |
| rwp11-mid-R | TCATGGGGCTTGACATCGGTGACGCAGGCACCAGTTGCCGC |
| rwp11-mid-F | CACCGATGTCAAGCCCCATGATG |
| RWP11_BamHI_R | CCCCGCCTCACCTGGATCCCTAGAAGTTGGTTGGATTCGCGGC |
| RPL23_SpeI-F | GCATATTCTTCAAGCCTGCTAGCGGAACTAGTAAGGCCCTCTGCGCAGA |
| RPL23_-R | CTCGAGACTCTGCGCGCAGACAAGCTCGAGACTCTGCGCGCAGAC |
| 3XHA_F2 | CTTGTCTGCGCGCAGAGTCTCGAGATGTCTAGTTACCCATACGATGTTCC |
| 3XHA_R3 | GGTCAGGCACGTCGAGGTCCATCCTAGGAGCGTAATCTGGAACGTCATATGG |
| rwp11_atg_f | ATGGACCTCGACGTGCCTGACC |
| RWP11_NsiI_R | GCTCAACGGTGACGGCGCCATGCATACGGCCAACCGACTACGGTATG |

| | |
|---|---|
| Sequencing | |
| M13R | CAGGAAACAGCTATGAC |
| genotyping | |
| 172 (RWP4_mc_F1) | CGCCGCTGCTACCAGACG |
| 173 (RWP4_mc_R1) | CGATAGCGCATTACTCAGACG |
| SYP1-F | TTTCCAGAACATTCATCCCC |
| SYP1-F2 | GGTATGGCGTGTCCTGGCGAAC |
| SYP1-R | GCTATTCACGTCGTTGCTCA |
| oMJ314 | TTGACCTGGAGGATCTGGAC |
| p359 | ACTGAGAATTCCTGCAGCCATGGCCTGTTTCTTAGCCAGGTTCC |
| p360 | ACTGAGGATCCCTAGCTAGCCATGTGTTTCTTGACGCTGGCGACC |
| p268 | GCTTACGAGGCTGTGACGCTTTTTG |
| p269 | TTTCCTAGCCGCTTGCGTTCGTTAC |
| P376 | AAGAACGATCGCCGACATAC |
| P377 | TCCTCCTCTTCCTCCTCCTC |
| oMJ913 | GCACCAATCATGTCAAGCCT |
| oMJ944 | GACGTTACAGCACACCCTTG |
| RPL23_P_F | GCGGCCCGCGACTGTTTC |
| sg175 | CAAGCGCTGACCAACCTGAG |
| 154 (RWP2_4197_F) | CTGACACAGCGGGCCTCAAGC |
| 335 (RWP2-4441_R) | CCCTCCATAGTCTACCAGTGG |
| sg223 (RWP2_E2R) | GTTGGGAAGTTCTCTTGCAATCGGC |
| sg96 (RWP2_5'UTR_F2) | GCAATGGTTCACGGAATACGAACC |
| 351 (VcMid3UTR.r1) | ACTGAGGTACCGAACAATTACTTTGCAACTGTCGCCATATAAAGC |
| 352 (VcMid3UTR.f1) | ACTGAGCTAGCTAAGGATCCATTTGCGATTGCTTGCTCTTCGTACCG |
| 139 (FSI1.SemiQ.f1) | TTGGCCTGTGTGCTGTTACTGAAGC |
| 140 (FSI1.SemiQ.r1) | CACCTTATGGTTGTGCAGGAAGCCC |
| qRT-PCR | |
| 18S_F1 | GAAGACGATTAGATACCGTCGTAGTCTC |
| 18S_R1 | TCCTTTAAGTTTCAGCCTTGCGAC |
| VcGSP1_qF6 | AGCCGCTGCCAACAGAATA |
| VcGSP1_qR6 | GTCATTCCCTCCAAGCCATCTAC |
| S0212FM_F1 | CGGAGATCAAATCACACCCTTGG |
| S0212FM_R1 | CTTGCTCTTGTAGGCCTTGTTCC |
| Vc0086_qR1 | CTTCGTCTCAGGTGCAGTAGTAA |
| Vc0086_qF2 | ACAATTGCTGCCCGACCTAT |
| VcGSM1_F4 | GTGTGTGTGTGTTGCTGTGC |
| VcGSM1_R4 | GCGAAATTTCCCCCAATAAT |
| RPL36a_F | GCCAAGACCACCAAGAAGATCG |

| | |
|---|---|
| RPL36a_R | TTAGTTGCCCTTCTTCTTGTCACC |
| GSP_Fq1 | ACCACGACGCAGAGGCAATTATCA |
| GSP_Rql | CGGCGTTGTTGAGGTGCTTAATGT |
| SAG 1_Fq 1 | CTGCCAATAGCGTGTTTGTGGTGT |
| SAG l_Rq1 | CTACTGCCGTTTGTGTTGTTGCCT |
| GSMl_Fq1 | ACGACAACTTCGTGTGGCCCTA |
| GSM1_Rq 1 | TTAATCTGCGTGTTGTTCAGCGCC |
| P446-SAD1_Fq1 | CAGCCGTTCAAACAGTCAACAGCA |
| P447-SAD1_Rq1 | TTGGGAACCGACATGTAGCGAAGA |

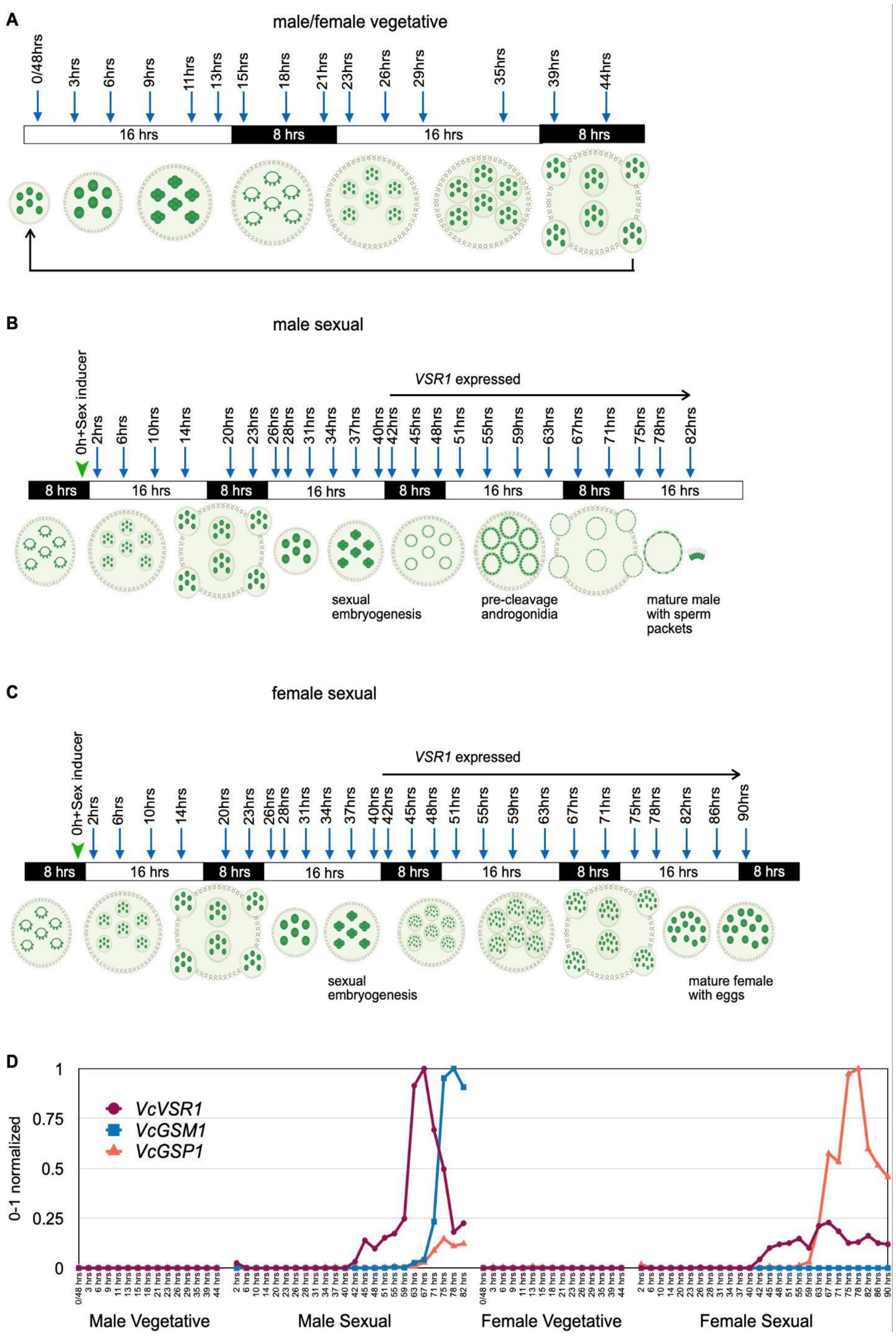


**Figure S1.** Time points and selected gene expression data for whole spheroids taken across vegetative and sexual development. A-C. Schematics showing developmental phasing for vegetative phase males and females (A), sexually induced males (B), and sexually induced females (C). Light/dark boxes show light and dark phases of life cycle, cartoons show spheroid morphology and downward arrows show sampling times. Addition of sex inducer in (B-C) is indicated by an arrowhead, and time of *VSR1* expression is shown by a horizontal arrow. D. Expression profiles of Volvox sex specific genes *VSR1*, *GSM1*, and *GSP1*. Vegetative and sexual phases with specific time points of female and male are labeled under the X axis. The Y axis shows expression normalized on a 0-1 scale with 1 being the maximum expression value observed.

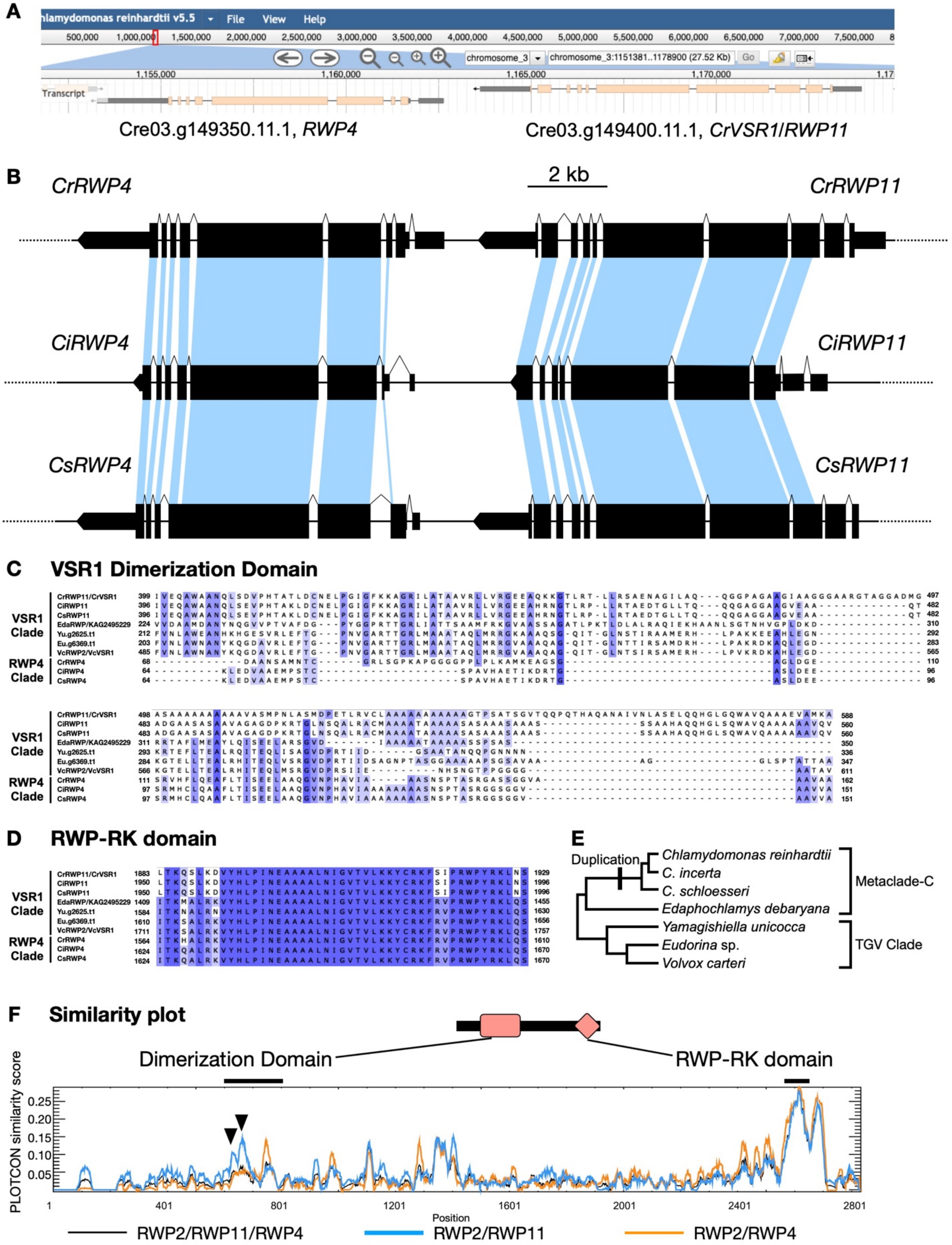


**Figure S2.** Sequence similarity among VSR1 homologs (RWP11 in Chlamydomonas, RWP2 in Volvox and other multicellular volvocine algae) and RWP4 homologs (found in Chlamydomonas and some close relatives). (A) Snapshot from Phytozome (26) genome browser showing adjacent location of *RWP4* and *RWP11* (*VSR1*) genes. (B) Synteny of *RWP11* and *RWP4* tandem structure among the core *Reinhardtinia* species *Chlamydomonas reinhardtii* (*Cr)*, *Chlamydomonas incerta* (*Ci)* and *Chlamydomonas schloesseri* (*Cs*). Corresponding coding exons are connected by blue shading. (C-E) Multiple sequence alignments of VSR1 and RWP4 predicted dimerization domains (C) and RWP-RK domains (D) aligned with MUSCLE (23). Residue conservation >50%, >70% and >90% identity shaded light, medium and dark purple. (E) Phylogram of the organisms from which the VSR1 homologs were aligned in B and C based on Craig *et al*. (24). Branch containing predicted RWP4/11 tandem duplication is marked with a thick vertical line. Branch length is not proportionate to the estimated divergence time. (F) Similarity score plot among RWP2/4/11 (black), VSR1 homologs only (RWP2 and RWP11) (blue), and between RWP2 and RWP4 (orange) homologs generated using PLOTCON (25). Regions spanning the dimerization domain and RWP-RK domains are shown. Inverted black triangles show peaks of similarity for the dimerization domain that are more conserved in VSR1 homologs (RWP2, RWP11) than in RWP4.

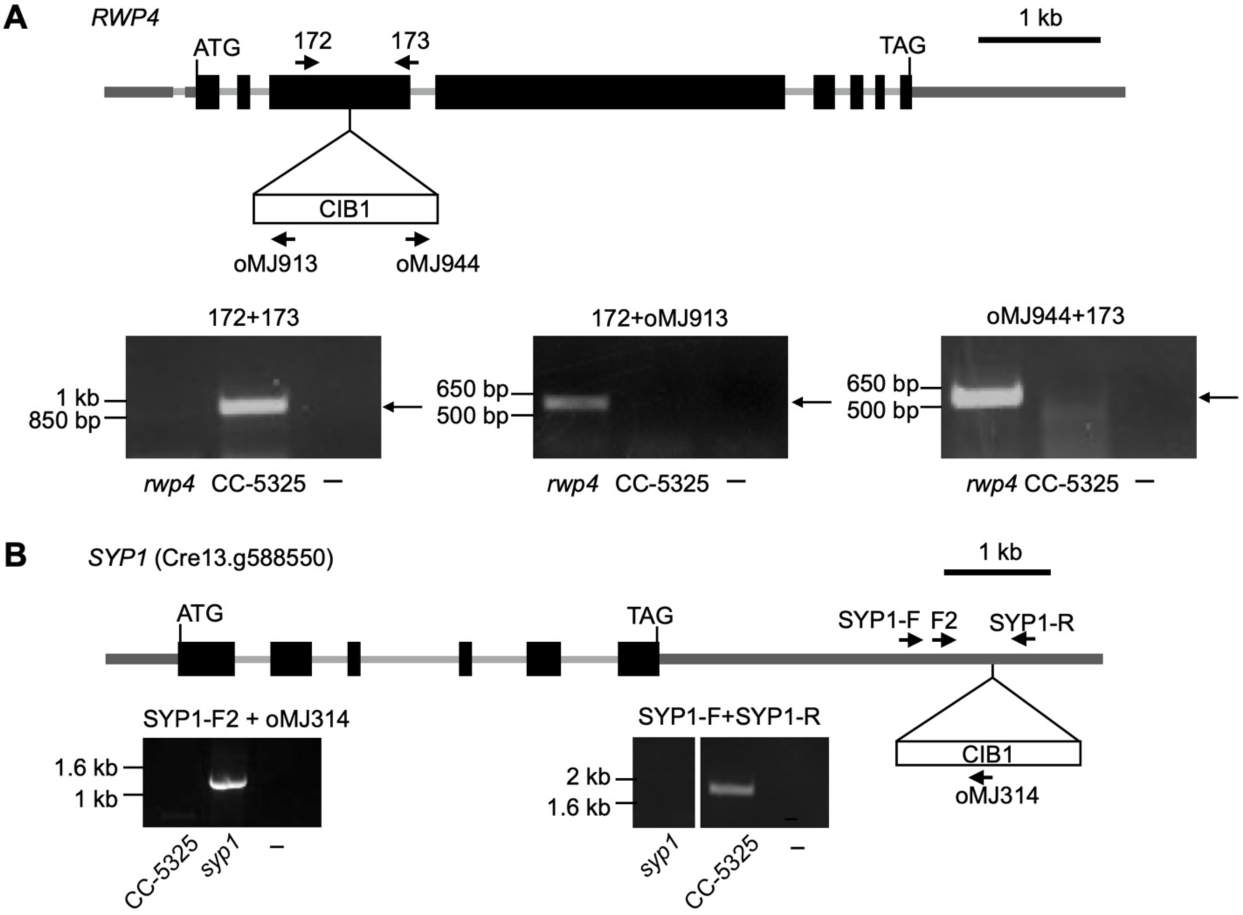


**Figure S3.** *RWP4* and *SYP1* loci in Chlamydomonas, and PCR genotype validation. A. Schematic of *rwp4* CLiP allele LMJ.RY0402.060132 with location of CIB1 cassette insertion and genotyping primers. Black boxes represent exons, thin gray lines, introns, and dark gray UTRs. Positions of start (ATG) and stop (TAG) codons are also indicated. PCR products used were separated on agarose gels and stained to visualize as in Figure 2B. B. While the CLiP strain LMJ.RY0402.189640 contains a CIB1 cassette insertion in the *VSR1* (*RWP11*) gene as shown in Figure 2A, it also has a second unlinked CIB1 cassette insertion in the 3' UTR of the *SYP1* gene as shown in the diagram and labeled similarly to that in panel A. Backcrossing experiments and genotyping of LMJ.RY0402.189640 strains that were rescued with a *VSR1* transgene were used to identify progeny lacking the *syp1* insertion mutation but which contained the *vsr1* mutation with or without a rescuing transgene as illustrated in Figure 2.

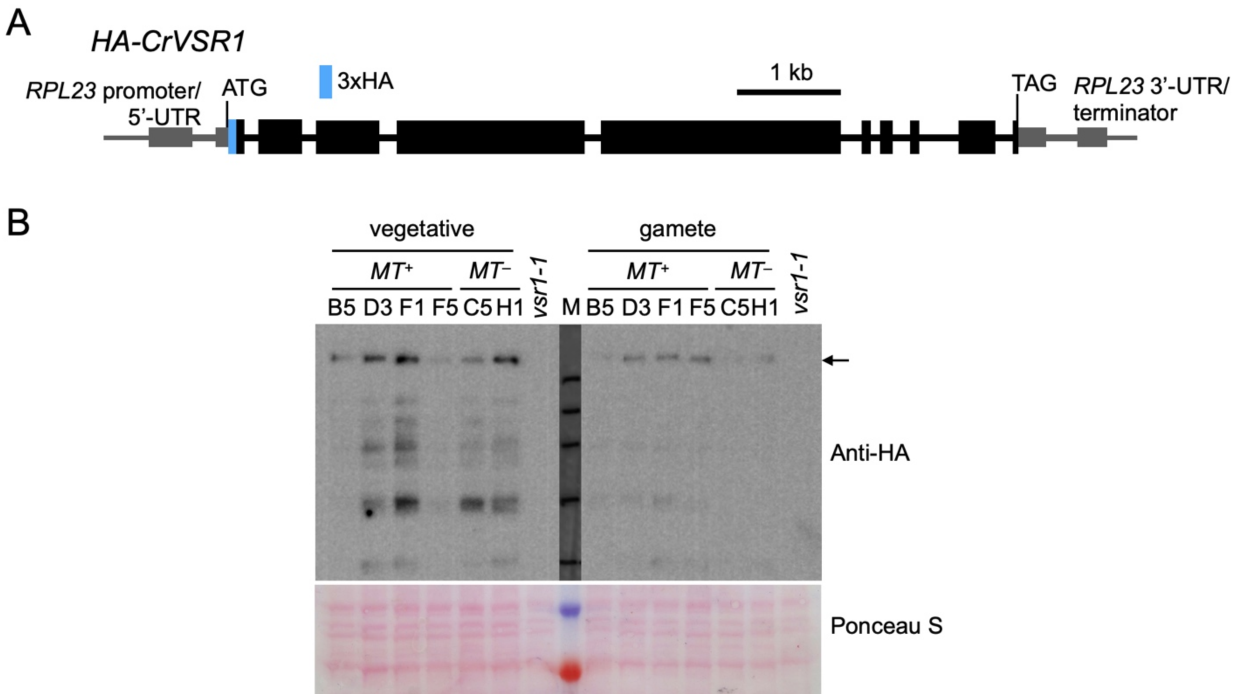


**Figure S4.** Schematic of the Chlamydomonas *HA-VSR1* construct (A) and anti-HA immunoblot (B) of SDS-PAGE fractionated protein extracts of *vsr1 HA-CrVSR1* and *vsr1* (control). The membrane was pre-stained with Ponceau S (bottom) as a loading control. Black arrow shows the predicted migration for full length 3xHA-CrVSR1 protein (210kDa). M, Marker, largest band size is 245kDa.

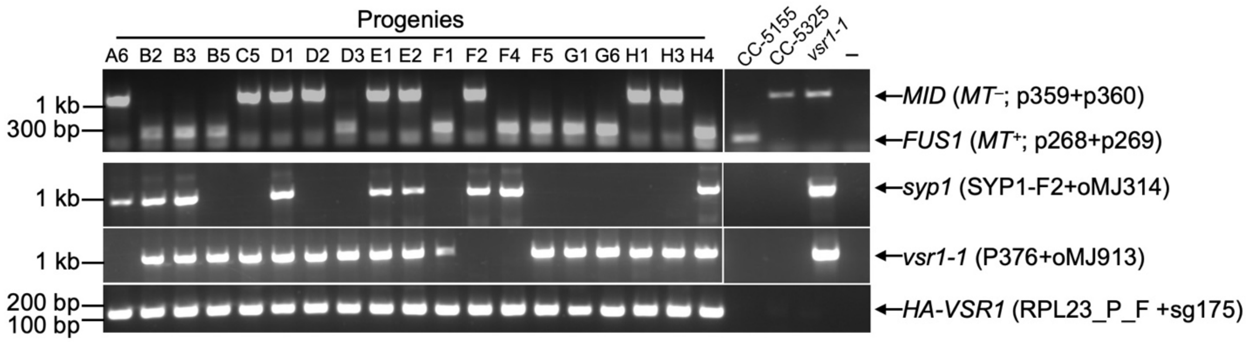


**Figure S5.** Genotyping progeny from backcross of rescued Chlamydomonas *vsr1* strain LMJ.RY0402.189640 (*vsr1-1 HA-VSR1 syp1*) described in Figure 2 and Figure S4B. *MID* and *FUS1* primers were used to determine mating type.

| | | Target #1 / PAM | | |
|---|---|---|---|---|
| wild type | +2272 | ......CTCTCCAGTACTGCTCTCGTGGGGGGGAGGGACTGTGGCGTCTC...... | +2229 | In/Del |
| *vsr1*#1_1 |  | ......CTCTCCAGTACTGCTCTCGATGGG--------CTGTGGCGTCTC...... |  | -8 bp |

| | | Target #5 / PAM | | |
|---|---|---|---|---|
| wild type | +4886 | ......GCTGCTGCTGGAGCTGTATGAGGTCCATTGCGCCACC-TGGCGGCGGCATCCCCACCATC...... | +4828 | In/Del |
| *vsr1*#5_1 | | ......GCTGCTGCTGGAGCTGTATGAGGTCCATTGCGCCACATTGGCGGCGGCATCCCCACCATC...... | | +1 |
| *vsr1*#5_2 |  | ......GCTGCTGCTGGAGCTGTATGAGGTCCATTGCGCC------------------CCACCATC...... |  | -14 |
| *vsr1*#5_3 |  | ......GCTGCTGCTGGAGCTGTATGAGGTCCATTGCGCCACC--------------------ATC...... |  | -19 |
| **vsr1*#5_4 | | ......GCTGCTGCTGGAGCTGTATGAGGTCCATTGCGCCACC (+20nt)(-10nt) ATCCCCACCATC...... | | +20,-10 |

**Figure S6.** Sequences of edited *VSR1* alleles from male Volvox strains in regions described in Figure 3A. Editing target sites (or protospacers) #1 and #5 are shown on the top and bottom respectively. Sequence positions relative to the start codon as +1 are shown for wild type. Note that these target sites are in the reverse complement orientation to the gene. **vsr1*#5_4 was designated *vsr1-1* and used for further experiments.

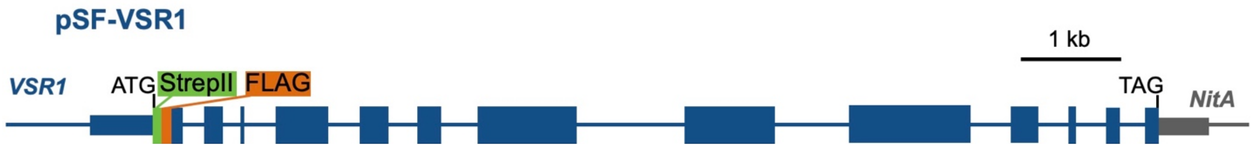

**Figure S7.** Diagram of pSF-VSR1 construct used to rescue the *vsr1-1* mutation. Coding sequences are wide bars, medium width bars are UTRs, and thin lines are intronic or non-coding regions. Native *VSR1* sequences are in blue. StrepII and FLAG epitope tag sequences at the N terminus are in green and orange, respectively. The *NitA* 3'UTR and terminator replace the native sequences past the stop codon and are shown in gray.

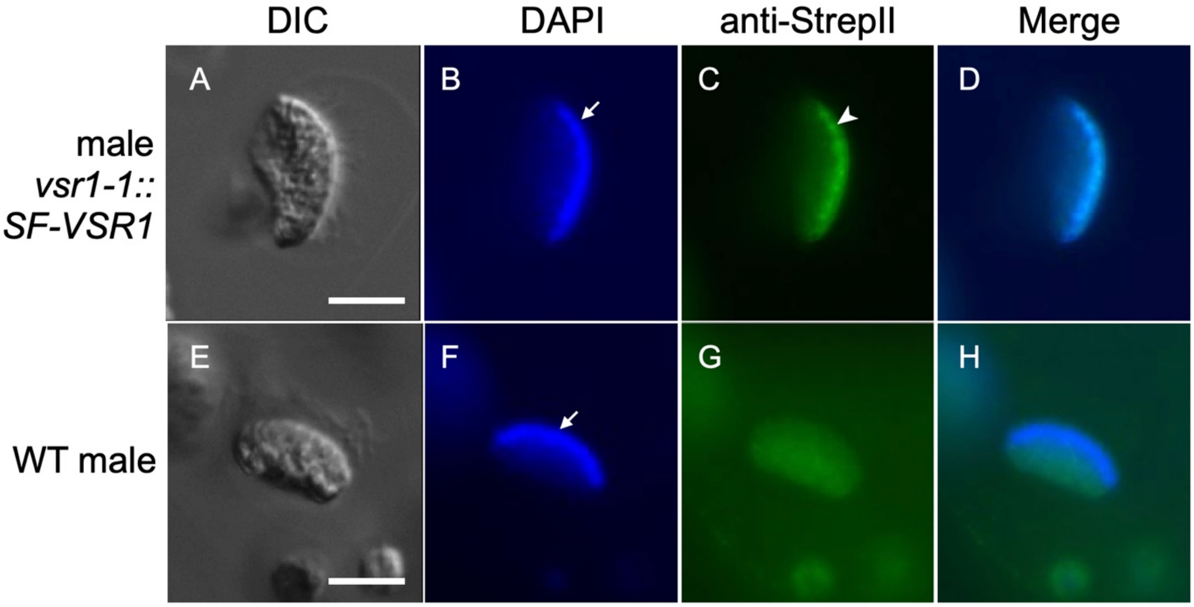


**Figure S8.** Immunolocalization of Volvox VSR1 in sperm nuclei. DIC images (A,E), DAPI staining false colored blue (B,F), StrepII tag immunofluorescence (IF) false colored green (C,G), and merged DAPI and StrepII (D,H) staining are shown for a rescued male *vsr1-1 SF-VSR1* sperm packet (B–D), or a wild type male sperm packet (F-H). Arrows and arrowheads show locations of a representative single nucleus from each image. Scale bars = 10 µm.

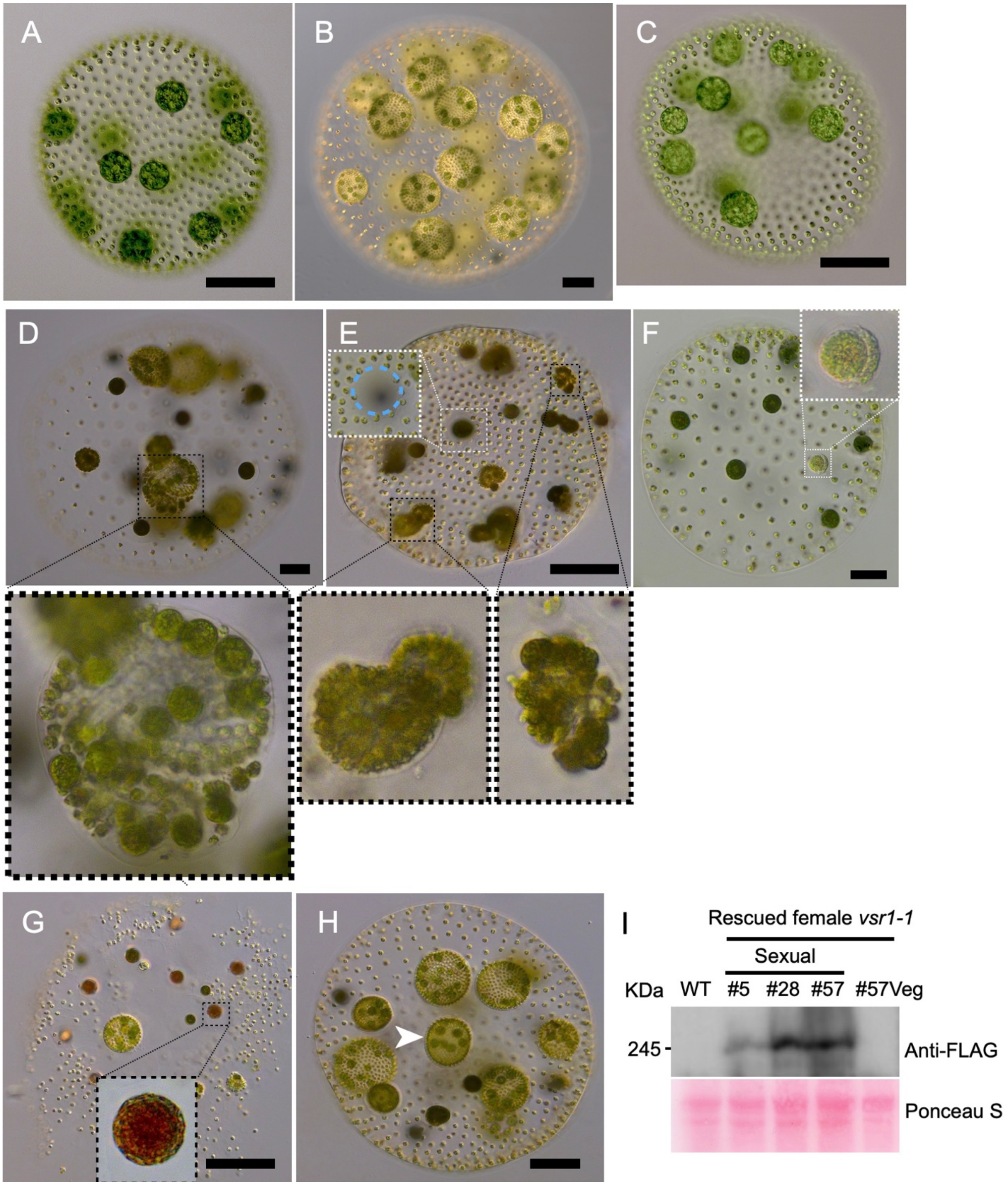


**Figure S9.** Characterization of Volvox female *vsr1-1* mutants and rescued strains in developmental stages not shown in Figure 3. A-B, Wild-type female spheroids in vegetative phase (A) and with unfertilized eggs that reverted to the vegetative stage (B). C-F. *vsr1-1* female mutant spheroids from vegetative phase (C), and after several days post sexual embryogenesis when the egg-like cells underwent aberrant vegetative embryogenesis with defects in completing inversion (D plus inset). When mature *vsr1* sexual females were mixed with wild-type male sperm (E), the sperm created fertilization pores (e.g. white dashed inset with blue dashed circle around the pore) through which they entered the *vsr1* female ECM but failed to complete fertilization. Instead, the egg-like cells behaved as unfertilized *vsr1-1* mutants and underwent vegetative embryogenesis with morphogenesis defects (insets with black dashed insets). Mature *vsr1-1* sexual females mixed with sperm sometimes had sperm attached to the outside of egg-like cells that persisted for days until the egg-like cell began de-differentiating (F). Sperm stuck on the outside of eggs were not observed in wild-type matings. G-H Rescued *vsr1-1 SF-VSR1* female spheroids can be fertilized successfully after sexual development and produce viable zygotes (G). If not fertilized (H) the eggs undergo normal vegetative embryogenesis without morphogenesis defects seen in *vsr1-1* mutants as in panels D and E. Scale bars are 50 μm. I. Immunoblotting similar to Figure 3J using a sexually induced female wild type (WT) strain, or three different rescued female segregants (#5, #28, #57). The right-most lane was loaded with a lysate from vegetative stage line #57 spheroids.

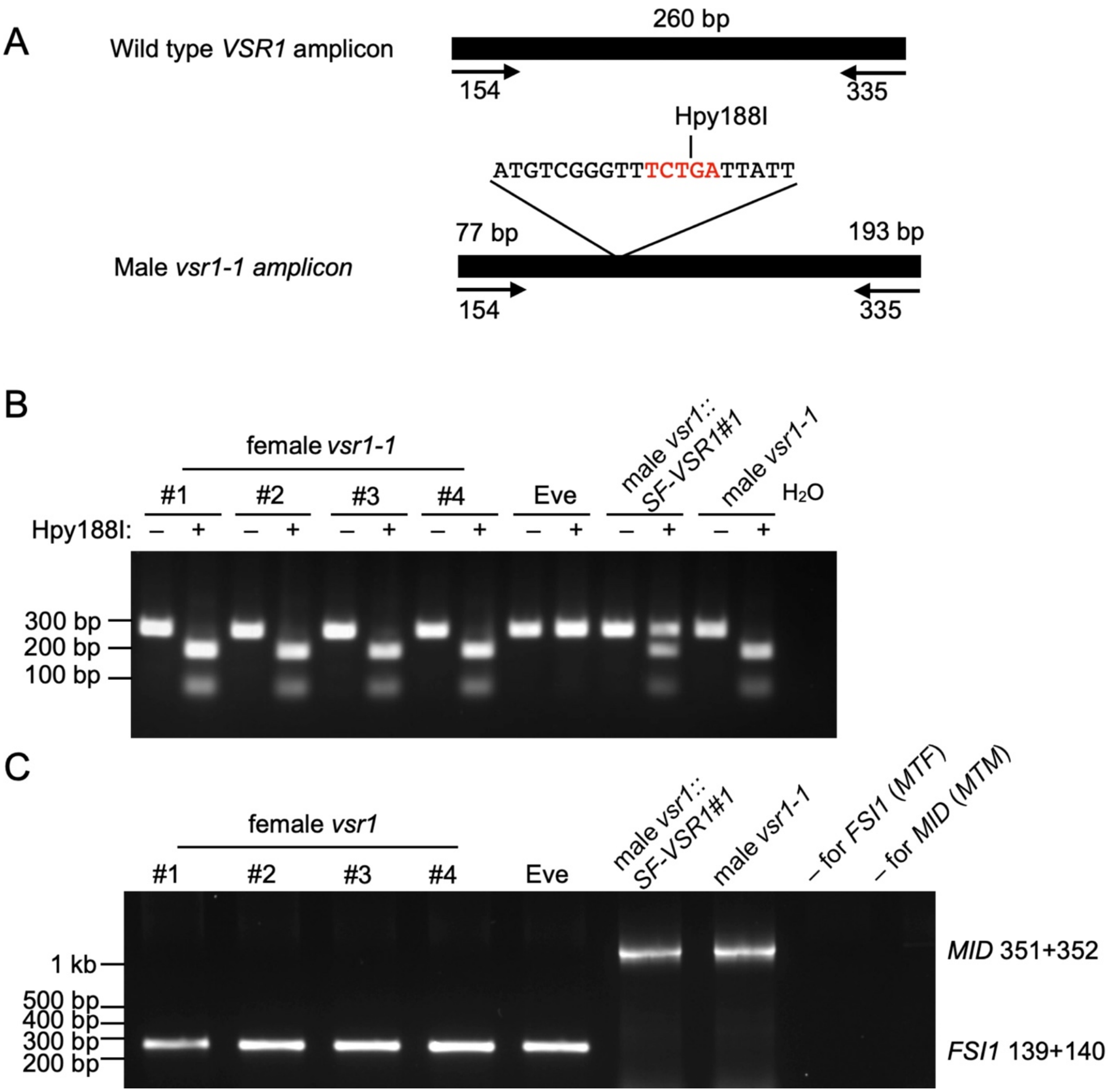
A
Wild type VSR1 amplicon
260 bp
154
335
Hpy188I
ATGTCGGGTTTCTGATTATT
77 bp
193 bp
Male vsr1-1 amplicon
154
335
B
female vsr1-1
#1
#2
#3
#4
Eve
male vsr1::SF-VSR1#1
male vsr1-1
H2O
Hpy188I:
300 bp
200 bp
100 bp
C
female vsr1
#1
#2
#3
#4
Eve
male vsr1::SF-VSR1#1
male vsr1-1
– for FSI1 (MTF)
– for MID (MTM)
1 kb
500 bp
400 bp
300 bp
200 bp
MID 351+352
FSI1 139+140

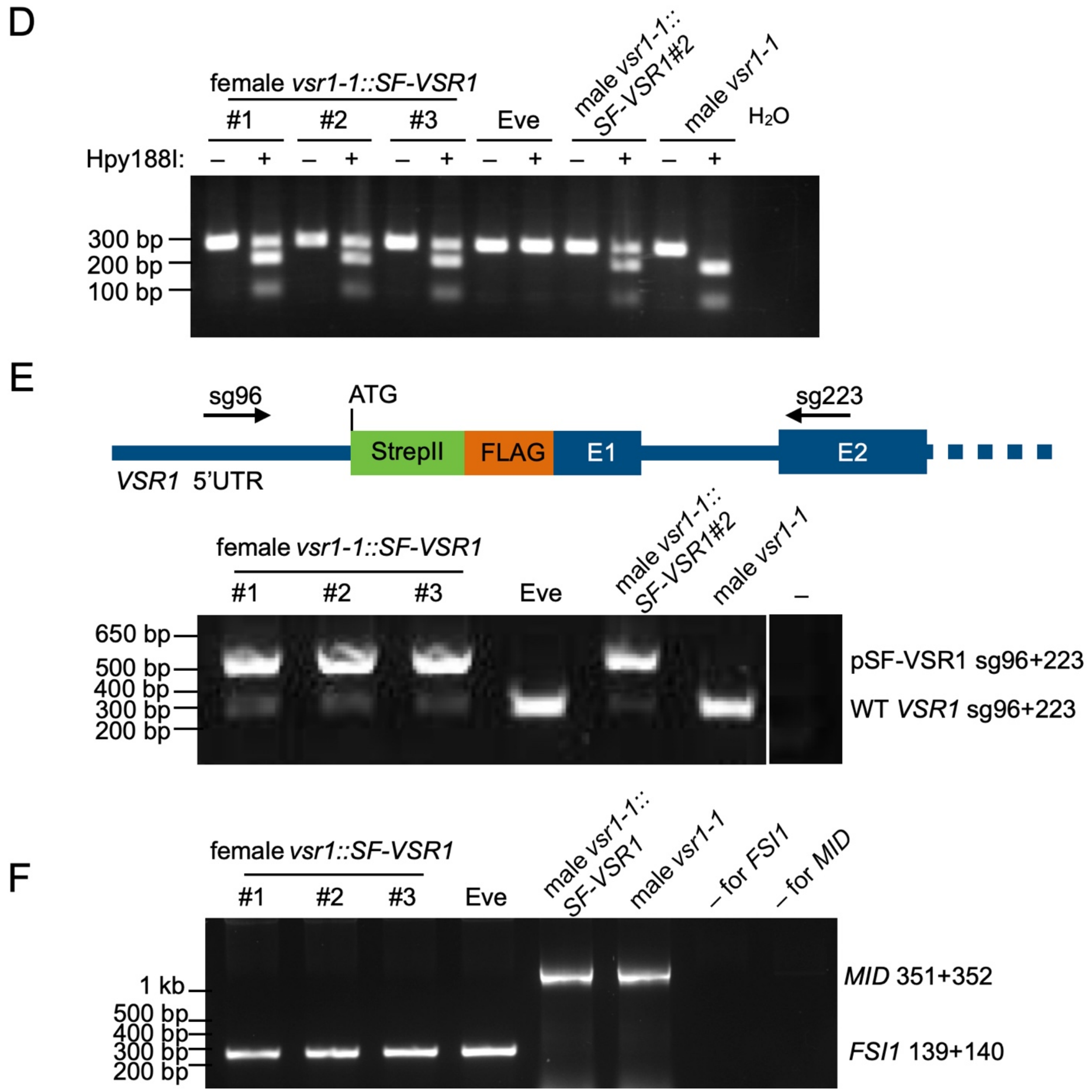


**Figure S10.** PCR genotyping of Volvox *vsr1-1* mutant. A. Diagram of wild type *VSR1* 260 bp amplicon produced with numbered primers shown by arrows (top) and the 270 bp amplicon produced in the *vsr1-1* mutant which adds 10 non-templated bases that contain a Hpy188I restriction site (shown in red lettering). B. Validation of genotyping primers for Volvox *vsr1-1* allele. Hpy188I digested (+) or undigested (-) PCR products were separated on an agarose gel where the mutant allele produces a 77 bp and 193 bp digestion product. Data are shown from four female *vsr1-1* segregants, wild-type female Eve, a rescued *vsr1-1* male strain, and an unrescued male strain. A no-template control ($H_2O$) was loaded in the last lane. C. Genotyping the sex determining region for males (*MTM*) based on amplification of the *MID* gene, and females (*MTF*) based on amplification of the *FSI1* gene (27). A no-template control for each primer pairs is designated as –. D. Genotyping rescued female *vsr1-1* mutant segregants similar to panel B. E. Top, diagram showing region of VSR1 where the epitope tag was inserted in the rescuing transgene and the location of genotyping primers (sg96: RWP2_5'UTR_F2; sg223: RWP2_E2R) for distinguishing the endogenous versus the rescuing allele. Bottom, validation of PCR genotyping primers for the SF-VSR1 transgene where the transgene produces a larger 560 bp amplicon than the endogenous *VSR1* product of 284 bp. F. Genotyping the sex determining region of rescued female *vsr1-1* mutants as in panel C.

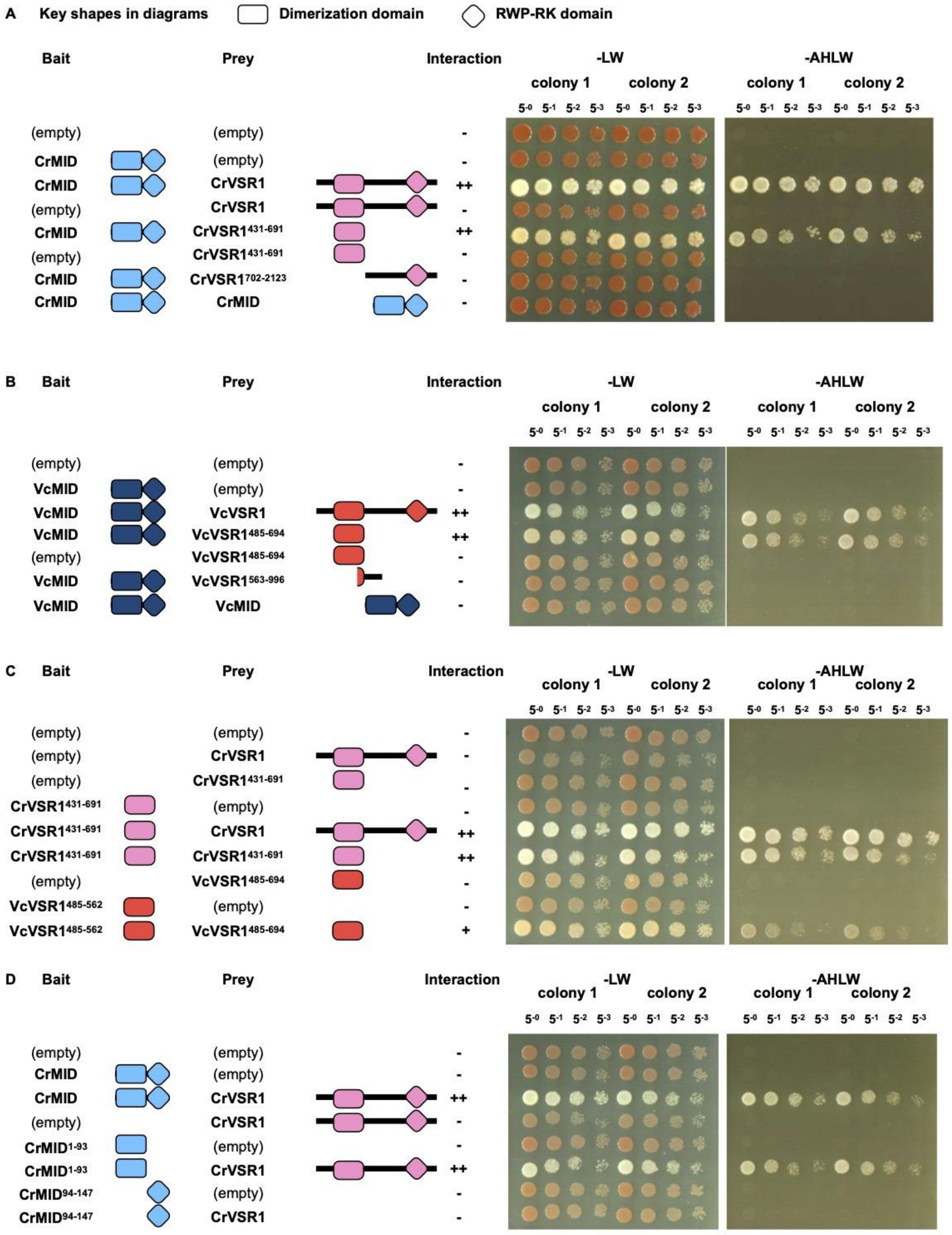


**Figure S11.** Yeast two-hybrid data with full plate images including independent replicate transformants.

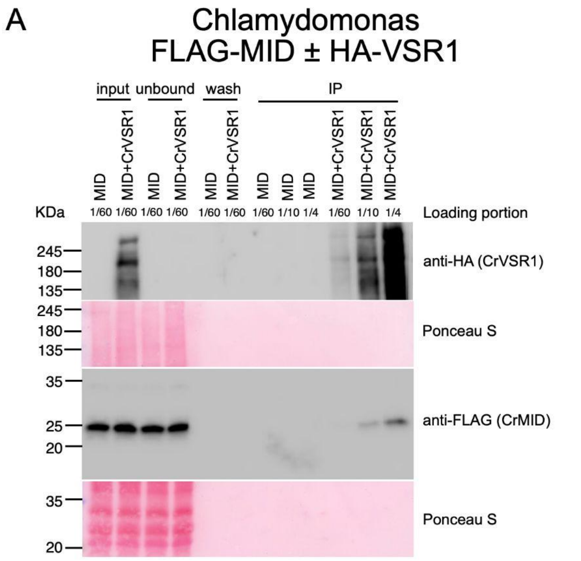


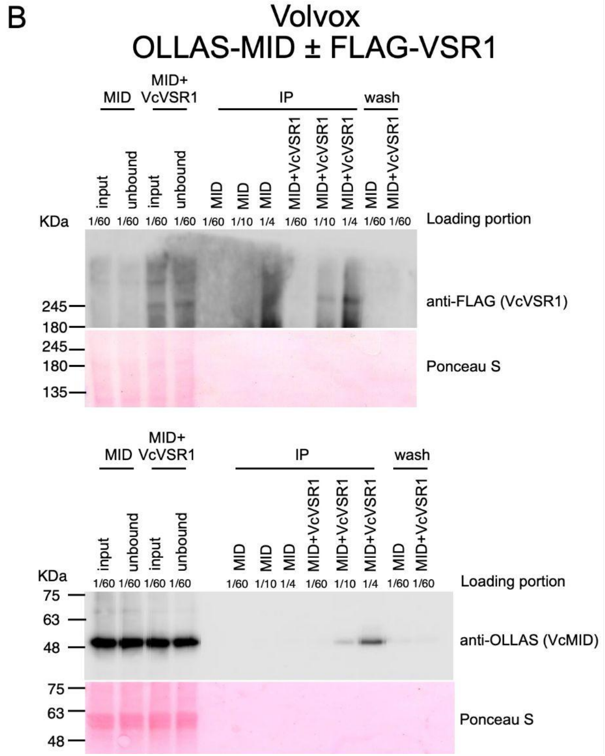


**Figure S12.** Full blot images of co-IP experiments from Figure 5. Immunoblot detection of CrVSR1 and CrMID Co-IP (A) and VcVSR1 and VcMID (B). Loading proportions based on total starting lysate are shown above each lane.

**Dataset S1 (separate file).** Amino acid multiple alignment of *VSR1* paralogs of core *Reinhardtinia*.

**Dataset S2 (separate file).** Summary of plasmid construction for yeast two-hybrid assay.

SI References


1. S. M. Miller, R. Schmitt, D. L. Kirk, Jordan, an active Volvox transposable element similar to higher plant transposons. *Plant Cell* **5**, 1125–1138 (1993).
2. S. Geng, P. De Hoff, J. G. Umen, Evolution of sexes from an ancestral mating-type specification pathway. *PLoS Biol* **12**, e1001904 (2014).
3. O. Ruecker, K. Zillner, R. Groebner-Ferreira, M. Heitzer, Gaussia-luciferase as a sensitive reporter gene for monitoring promoter activity in the nucleus of the green alga Chlamydomonas reinhardtii. *Molecular Genetics and Genomics* **280**, 153–162 (2008).
4. J. A. Ortega-Escalante, R. Jasper, S. M. Miller, CRISPR/Cas9 mutagenesis in Volvox carteri. *Plant J.* **97**, 661–672 (2019).
5. J. M. Short, J. M. Fernandez, J. A. Sorge, W. D. Huse, Lambda ZAP: a bacteriophage lambda expression vector with in vivo excision properties. *Nucleic Acids Res.* **16**, 7583–7600 (1988).
6. C. López-Paz, D. Liu, S. Geng, J. G. Umen, Identification of Chlamydomonas reinhardtii endogenous genic flanking sequences for improved transgene expression. *Plant J.* **92**, 1232–1244 (2017).
7. M. Heitzer, B. Zschoernig, Construction of modular tandem expression vectors for the green alga Chlamydomonas reinhardtii using the Cre/lox-system. *BioTechniques* **43**, 324, 326, 328 passim (2007).
8. B. J. S. C. Olson, *et al.*, Regulation of the Chlamydomonas cell cycle by a stable, chromatin-associated retinoblastoma tumor suppressor complex. *Plant Cell* **22**, 3331–3347 (2010).
9. M. Fuhrmann, W. Oertel, P. Hegemann, A synthetic gene coding for the green fluorescent protein (GFP) is a versatile reporter in Chlamydomonas reinhardtii. *The Plant Journal* **19**, 353–361 (1999).
10. T. Jakobiak, *et al.*, The bacterial paromomycin resistance gene, aphH, as a dominant selectable marker in Volvox carteri. *Protist* **155**, 381–393 (2004).
11. D. Koszegi, *et al.*, Members of the RKD transcription factor family induce an egg cell-like gene expression program. *Plant J.* **67**, 280–291 (2011).
12. F. Tedeschi, P. Rizzo, T. Rutten, L. Altschmied, H. Bäumlein, RWP-RK domain-containing transcription factors control cell differentiation during female gametophyte development in Arabidopsis. *New Phytol.* **213**, 1909–1924 (2017).
13. T. Waki, T. Hiki, R. Watanabe, T. Hashimoto, K. Nakajima, The Arabidopsis RWP-RK protein RKD4 triggers gene expression and pattern formation in early embryogenesis. *Curr. Biol.* **21**, 1277–1281 (2011).
14. K. Nakajima, Be my baby: patterning toward plant germ cells. *Curr. Opin. Plant Biol.* **41**, 110–115 (2018).
15. S. Koi, *et al.*, An Evolutionarily Conserved Plant RKD Factor Controls Germ Cell Differentiation. *Current Biology* https:/doi.org/10.1016/j.cub.2016.05.013 (July 4, 2016).
16. M. Rövekamp, J. L. Bowman, U. Grossniklaus, Marchantia MpRKD Regulates the Gametophyte-Sporophyte Transition by Keeping Egg Cells Quiescent in the Absence of Fertilization. *Curr. Biol.* (2016) https:/doi.org/10.1016/j.cub.2016.05.028.
17. H. Sekimoto, *et al.*, A divergent RWP-RK transcription factor determines mating type in heterothallic Closterium. *New Phytol.* **237**, 1636–1651 (2023).
18. R. Blanc-Mathieu, *et al.*, Population genomics of picophytoplankton unveils novel chromosome hypervariability. *Sci. Adv.* **3** (2017).
19. A. Z. Worden, *et al.*, Green evolution and dynamic adaptations revealed by genomes of the marine picoeukaryotes Micromonas. *Science* **324**, 268–272 (2009).
20. T. Yamazaki, *et al.*, Genomic structure and evolution of the mating type locus in the green seaweed Ulva partita. *Sci. Rep.* **7**, 11679 (2017).
21. X. Liu, *et al.*, Transcriptional dynamics of gametogenesis in the green seaweed Ulva mutabilis identifies an RWP-RK transcription factor linked to reproduction. *BMC Plant Biol.* **22**, 19 (2022).
22. P. J. Ferris, U. W. Goodenough, Mating Type in Chlamydomonas Is Specified by mid, the Minus-Dominance Gene. *Genetics* **146**, 859–869 (1997).
23. R. C. Edgar, MUSCLE: multiple sequence alignment with high accuracy and high throughput. *Nucleic Acids Res.* **32**, 1792–1797 (2004).
24. R. J. Craig, A. R. Hasan, R. W. Ness, P. D. Keightley, Comparative genomics of Chlamydomonas. *Plant Cell* **33**, 1016–1041 (2021).

25. P. Rice, I. Longden, A. Bleasby, EMBOSS: the European Molecular Biology Open Software Suite. *Trends Genet.* **16**, 276–277 (2000).
26. D. M. Goodstein, *et al.*, Phytozome: a comparative platform for green plant genomics. *Nucleic Acids Res.* **40**, D1178-86 (2012).
27. P. Ferris, *et al.*, Evolution of an expanded sex-determining locus in Volvox. *Science* **328**, 351–354 (2010).
